# Characterizing nascent transcription patterns of PROMPTs, eRNAs, and readthrough transcripts in the ENCODE4 deeply profiled cell lines

**DOI:** 10.1101/2024.04.09.588612

**Authors:** Ariel McShane, Ishwarya Venkata Narayanan, Michelle T. Paulsen, Mario Ashaka, Hailey Blinkiewicz, Nina T. Yang, Brian Magnuson, Karan Bedi, Thomas E. Wilson, Mats Ljungman

**Affiliations:** Cellular and Molecular Biology Program, University of Michigan Medical School, Ann Arbor, Michigan 48109, USA; Department of Radiation Oncology, University of Michigan Medical School, Ann Arbor, Michigan 48109, USA; College of Literature, Science, and Arts, University of Michigan, Ann Arbor, Michigan 48109, USA; Department of Pathology, University of Michigan Medical School, Ann Arbor, Michigan 48109, USA; Department of Biostatistics, School of Public Health, University of Michigan, Ann Arbor, Michigan 48109, USA; Rogel Cancer Center, University of Michigan Medical School, Ann Arbor, Michigan 48109, USA; Department of Human Genetics, University of Michigan Medical School, Ann Arbor, Michigan 48109, USA; Department of Environmental Health Sciences, School of Public Health, University of Michigan, Ann Arbor, MI 48109, USA; Center for RNA Biomedicine, University of Michigan, Ann Arbor, Michigan 48109, USA

## Abstract

Arising as co-products of canonical gene expression, transcription-associated lincRNAs, such as promoter upstream transcripts (PROMPTs), enhancer RNAs (eRNAs), and readthrough (RT) transcripts, are often regarded as byproducts of transcription, although they may be important for the expression of nearby genes. We identified regions of nascent expression of these lincRNA in 16 human cell lines using Bru-seq techniques, and found distinctly regulated patterns of PROMPT, eRNA, and RT transcription using the diverse biochemical approaches in the ENCODE4 deeply profiled cell lines collection. Transcription of these lincRNAs was influenced by sequence-specific features and the local or 3D chromatin landscape. However, these sequence and chromatin features do not describe the full spectrum of lincRNA expression variability we identify, highlighting the complexity of their regulation. This may suggest that transcription-associated lincRNAs are not merely byproducts, but rather that the transcript itself, or the act of its transcription, is important for genomic function.

## Introduction

Transcription of protein-coding genes (pc-genes) by RNA polymerase II (RNAPII) occurs in three distinct phases: initiation, elongation, and termination^1^. Transcription initiation is highly regulated by the binding of pioneer transcription factors (TFs), remodeling of chromatin to form nucleosome-depleted regions (NDRs), and binding of general TFs, cofactors, and RNAPII at promoters in NDRs to form the preinitiation complex (PIC)^2^. Following initiation, RNAPII promoter-proximal pausing and productive elongation into the gene body occur, regulated by elongation factors, histone modifications, and RNAPII carboxy-terminal domain (CTD) phosphorylation events^3,4^. Finally, 3’-end processing and transcription termination occurs upon slowing of RNAPII, recognition of a polyadenylation site (PAS), and recruitment of the cleavage and polyadenylation (CPA) complex^2^. Transcription can also be influenced by distal regulatory mechanisms and 3D chromatin conformation^3,5^. While these phases of transcription are highly regulated, nascent RNA-seq uncovered the prevalence of short-lived lincRNAs that arise from intergenic regions by transcription mechanisms that are less well defined and may differ from pc-genes^2^. Interestingly, some of these lincRNAs are produced during the process of canonical transcription^6–9^ from promoters in NDRs shared with genes (PROMPTs^10^) and enhancers (eRNAs^11^), as well as from readthrough transcription downstream of genes (RT transcripts^12^). While some view these transcription-associated lincRNAs as byproducts, these transcripts, or the act of their transcription, may be important for the expression of proximal genes^2,6^. Thus, further study of their expression patterns and functions could advance our understanding of transcriptional regulation. Utilizing transcriptomics and functional genomics data generated during the fourth phase of the ENCODE project (ENCODE4), we assessed the prevalence of PROMPT, eRNA, and RT transcription in 16 human cell lines. We describe their distinct patterns of expression relative to their associated genes, in addition to identifying sequence and chromatin features that correlate with their transcription. While we report features that distinguish the expression of these lincRNAs from their associated gene, the patterns of histone modifications, 3D genome architecture, and sequence motifs identified only highlight some of the potential mechanisms that may regulate their transcription.

## Results

### Exploring the transcriptional landscape with the ENCODE4 deeply profiled cell lines

For this study, we used data from the ENCODE4 deeply profiled cell lines (DPCL). The DPCL is a unique sample collection composed of data from 16 human cell lines (2 biological replicates) that were grown in one laboratory to minimize batch effects and distributed to 8 ENCODE consortium laboratories to perform 13 diverse biochemical assessments (Fig. 1a). Represented in the DPCL are 6 assays with 9 unique modalities probing different features of transcription, including the steady-state transcriptome (total RNA-seq), RNA synthesis and turnover (Bru-seq and BruChase-seq 2h/6h, respectively), transcript isoform diversity (long-read RNA-seq), transcription initiation (BruUV-seq, PRO-cap), and small non-coding RNA expression (miRNA-seq), as well as the transcriptional response to a perturbation (Bru-seq with ionizing radiation). Additionally, there are 5 assays with 6 unique modalities describing the chromatin landscape of the cell, including accessibility (ATAC-seq, snATAC-seq, DNase-seq) and 3D organization (Intact Hi-C, POLR2A ChIA-PET, CTCF ChIA-PET). Together, these bioassays provide a unique opportunity to explore the form and function of the human genome in a set of matched biosamples (ENCODE4 Flagship 2024).

**Figure 1.**
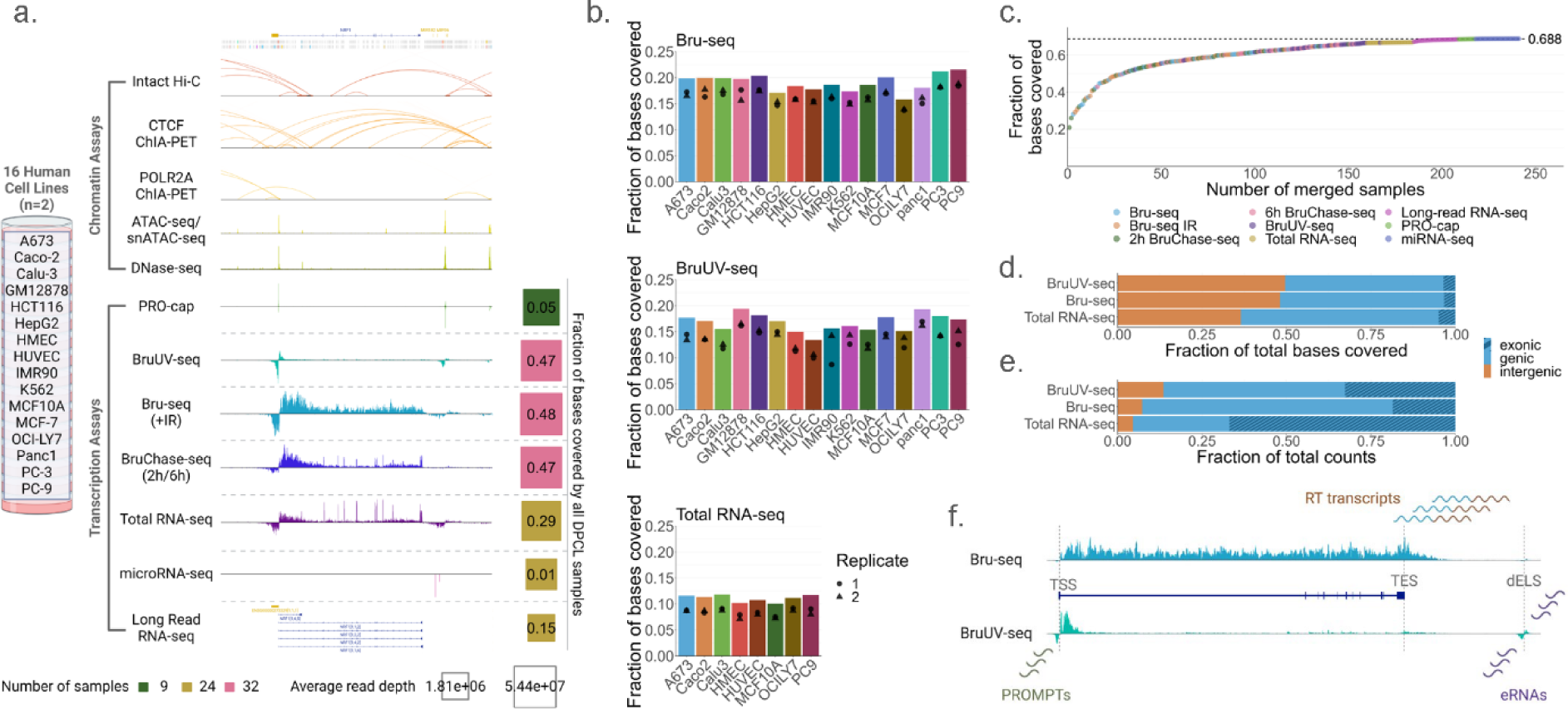
The ENCODE4 DPCL data can be used to study functional genomics in 16 human cell lines. **a.** Schematic describing the ENCODE4 DPCL collection of data (N = 2 biological replicates). Signal tracks are included for a 274.5 kb region containing the NRF gene, MIR182, and MIR96, in Caco-2 (ENCBS095ZZD) or HepG2 (ATAC-seq, ENCBS193JTP, ENCBS041EUO) for all assays except long-read RNA-seq where known and novel transcript isoforms are displayed^14^. Bulk ATAC-seq, untreated Bru-seq, and 6h BruChase-seq data is shown where multiple assays are listed. The percentage of bases covered cumulatively per DPCL transcription assay is included, with number of libraries and average read depth of all samples represented. **b.** The fraction of bases covered for each cell line is shown for selected assays. Cell line coverages are calculated from merged replicate data, thus cell lines with 2 replicate DPCL samples per assay are shown. Individual replicate coverages are indicated by the points within the bars. **c.** The fraction of bases covered by all DPCL samples, where each point indicates the cumulative coverage after each sample was added. **d,e.** For selected assays (16 cell lines, 24 libraries), the cumulative fraction of bases covered (**d**) and total uniquely mapping reads (**e**) was calculated for genic and intergenic compartments of the genome (based on GENCODE v29), with the striped bar indicating the proportion of genic coverage over exons. **f.** Schematic representation of Bru-seq and BruUV-seq signal (ENCBS826LEP) at the DYRK1A gene. Characteristic differences between the two assays can be appreciated, including a 5’-bias and low gene body coverage in BruUV-seq along with signal enrichment at highly unstable RNA species. Annotated TSS, TES, and dELS locations are shown, and PROMPT, eRNA, and RT transcription can be seen up- and/or downstream of annotated genomic elements.

To explore the global transcriptional landscape, we examined the RNA coverage of the genome using the DPCL transcription assays. For each DPCL transcription assay, we calculated the cell line specific and cumulative proportion of bases covered on both strands, exploring RNA synthesis in nascent data alongside the genomic coverage of diverse RNA pools (Fig. 1a). 18-22% of bases were covered by nascent Bru-seq reads per cell line, due in part to epigenetic regulation, with 47.6% of the genome covered cumulatively by all cell lines (Fig. 1a-b). In contrast to nascent assays (Bru-seq, BruUV-seq), the proportion of bases covered by the other transcription assays was predictably lower (Fig. 1a-b, Extended Data Fig. 1a), due to their more mature RNA pool (total RNA-seq, longread RNA-seq) or more targeted nature (PRO-cap, miRNA-seq). However, 68.8% of bases were covered after compiling all DPCL data to increase depth and transcript diversity, approaching previous estimates of 75-85%^6,13^ which relied on unstranded total RNA-seq data to infer RNA synthesis from a primarily mature RNA pool (Fig. 1c, Supplementary Note 1).

Given the extensive transcription potential of the genome, we assessed the distribution of transcriptional activity in genic and intergenic compartments. We found that ∼50% of all bases covered by Bru-seq data are in the unannotated portion of the genome (∼30% per cell line), however, the fraction of reads is predictably higher in genes (Fig. 1d-e, Extended Data Fig. 1b-c). Despite the smaller fraction of RNA sequencing reads, a significant proportion of genomic space traversed by RNAPII is between genes, and this intergenic activity is more variable among cell lines (Extended Data Fig. 1d).

Intergenic transcription has partly been attributed to lincRNA co-products of canonical transcription^6,8,9^; particularly PROMPTs, eRNAs, and RT transcripts. We identified and characterized transcription-associated lincRNAs using Bru-seq and BruUV-seq, which provide more intergenic coverage due to the temporal proximity of nascent Bru-labeled RNA to synthesis, than datasets with more mature, processed RNA pools (e.g. total RNA-seq, Fig. 1d-e). Bru-seq allowed the investigation of nascent RT transcription, and BruUV-seq revealed the initiation patterns at highly unstable RNA species, including PROMPTs and eRNA, in part due to suppression of RNA exosome-mediated degradation^15,16^ (Fig. 1f, Supplementary Note 2).

### PROMPT transcription is regulated distinctly from genes

The observed signal profile at genic transcription start sites (TSSs) is typically bidirectional with transcripts arising from distinct promoters generating an asymmetrical output due to productive elongation of the pre-mRNA and rapid termination of the PROMPT^10,17–27^ PROMPT expression was described in the 2 kb upstream divergent region of 12939 annotated pc-gene TSSs (Extended Data Fig. 2a). Transcriptional profiles at gene TSSs were classified based on both PROMPT and pre-mRNA signal (Fig. 2a, Extended Data Fig. 2b). In all cell lines, PROMPT and gene pairs with promoters in an NDR displayed concordant expression with transcription of both features (bidirectional, 37.2%) or neither (negligible signal, 47.3%, Fig. 2a, Extended Data Fig. 2c-e). However, specific regulation could dictate the transcriptional output for these different regions. ∼14% of assayed TSSs displayed gene expression accompanied by very low PROMPT signal (low-confidence bidirectional, Fig. 2a, Extended Data Fig. 2d), with few unidirectional TSSs (PROMPT RPKM = 0, Extended Data Fig. 2e). In addition, only a moderate correlation was observed between PROMPT and gene expression for bidirectional TSSs^28^ (Pearson’s r 0.33-0.43, Extended Data Fig. 2f). To further evaluate TSS-proximal expression patterns at bidirectional TSSs in all cell lines, we calculated the PROMPT:Gene (P:G) RPKM ratio of 2 kb regions flanking the TSS. Expression over the gene body was typically higher^25–28^, with similar P:G ratio distributions across cell lines (Extended Data Fig. 2g). However, ∼27% of TSSs displayed higher PROMPT expression in at least one cell line (Fig. 2b), further suggesting independent regulation of the PROMPT.

**Figure 2.**
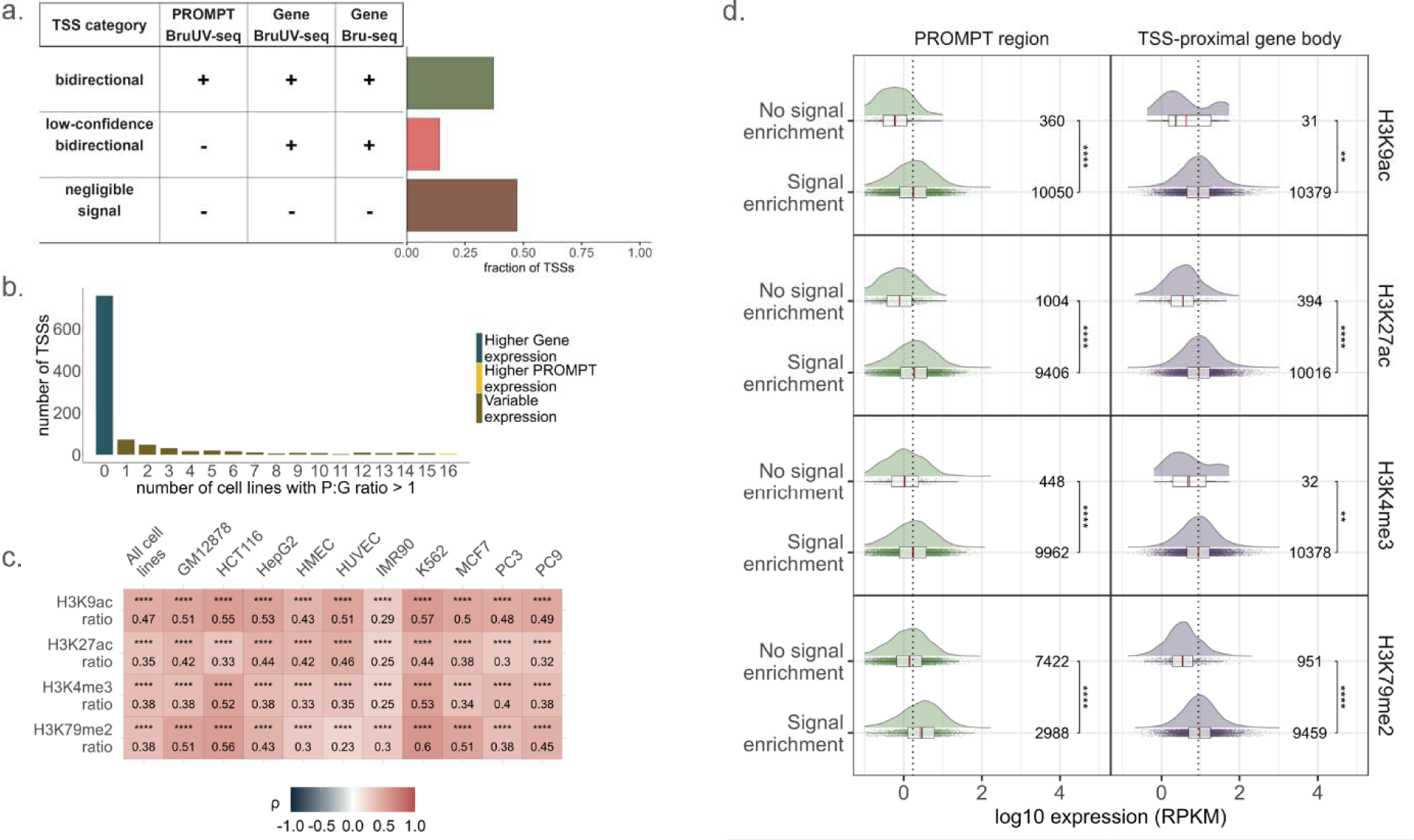
Discrete expression patterns at bidirectional Transcription Start Sites (TSSs) associate dynamically with activating histone modifications. **a.** Predominant TSS categories and an abridged set of parameters used for their classification. “+” or “-” indicates above or at/below the established parameter thresholds (see Methods: TSS category generation). Bar plot shows the fraction of TSSs for the categories for all cell lines (N_unambiguous_ TSS = 132799) **b.** Counts of TSSs with distinct expression patterns determined by TSS-proximal PROMPT:Gene (P:G) RPKM ratios for bidirectional TSSs common in all cell lines, N_TSS_ = 1041/12939. Higher Gen or PROMPT expression is indicated for patterns consistent across all 16 cell lines, Variable expression indicates P:G ratio > 1 for 1-15 cell lines. **c.** Heatmap of Spearman correlation coefficients (Iii) between selected histone modification ratios and expression (RPKM) ratios over PROMPT and gene regions flanking the TSS. False discovery rate (FDR) adjusted p-values are indicated by *p_adj_ < 0.05, **p_adj_ < 0.01, ***p_adj_ < 0.001, ****p_adj_ < 0.0001. **d.** Distribution of PROMPT or TSS-proximal gene expression (RPKM) based on histone signal enrichment status (determined by peak overlaps). The number of observations is denoted per signal enrichment group. Dotted vertical lines denote the median log10 expression values for PROMPT (0.237) and genic RNA (0.943), vertical lines in the boxplot represent the following: solid black line = median, solid red line = mean. Significance levels, determined from Wilcoxon signed-rank test FDR adjusted p-values, are represented as in **c**.

To discern factors influencing expression of a PROMPT and a gene at bidirectional TSSs, we assessed various genomic and chromatin features of these two TSS-proximal regions relative to each other (Extended Data Fig. 3a). For selected histone post-translational modifications, we observed varying deposition (signal and peaks) in PROMPT and gene regions, as well as a lower aggregated signal over the PROMPT for most modifications in all cell lines assayed (Extended Data Fig. 3b-c). Moderate positive correlations were observed between our P:G RPKM ratios and the corresponding P:G ratios of histone signal for 4 modifications–H3K9ac, H3K27ac, H3K4me3, and H3K79me2–in all cell lines, with others displaying low or variable correlations (Fig. 2c, Extended Data Fig. 3a,d-f). Relative to the other modifications, enrichment of H3K79me2 was found at a comparatively small subset of highly expressed PROMPTs, despite all four histone modifications being similarly enriched in the gene body (Fig. 2d), and this was true in most cell lines (Extended Data Fig. 4a-d). This selective distribution of H3K79me2 may indicate that it plays a more specific role in TSS-proximal regulation of PROMPT transcription.

### Asymmetrical transcription at enhancer-like elements is associated with distal promoter interaction direction

The regulatory duality of promoters and enhancers has been well described, however there remain distinctions in their activity and regulation^10^. Active enhancers are bidirectionally transcribed, like promoters, however, enhancer transcription is not expected to be inherently directional. Despite this, many enhancers exhibit gene-like characteristics, including asymmetrical transcription^28–31^. Because transcription at regulatory elements is influenced in part by distal interactions, we explored the expression patterns at enhancers and enhancer-like gene TSSs (eTSSs) engaged with distal promoters.

To identify enhancers with asymmetrical transcription, peaks of BruUV-seq signal (BruUV-peaks) were called with MACS2^32^, and those overlapping ENCODE4 distal enhancer-like candidate cis-regulatory elements^33^ (dELS cCREs) were denoted as regions of eRNA transcription. Enhancers were designated bidirectional when proximal, divergent BruUV-peak pairs were identified and unidirectional where individual BruUV-peaks had no detectable divergent signal. Unpaired peaks with signal on the divergent strand were designated as low-confidence bidirectional enhancers (Extended Data Fig. 5a). 82-99% of all BruUV enhancers were bidirectionally transcribed, with larger proportions of unidirectional peaks in cell lines with fewer BruUV-peaks (Extended Data Fig. 5b), which may result from technical or biological variability (Supplementary Note 2). Bidirectional enhancers tended toward signal symmetry^28,31,34^, and were classified into symmetrical and asymmetrical classes based on their BruUV-peak RPKM ratio (Extended Data Fig. 5a,c), excluding low-confidence bidirectional eRNA peaks. The asymmetrical groups were divided roughly evenly into enhancers where the plus-strand or minus-strand peak was dominant (asymmetrical plus and asymmetrical minus, respectively), as were unidirectional peaks (individual plus and individual minus). To assess eRNA signal relative to their distal interactions, we obtained enhancers that were shown to interact with an expressed gene TSS (RPKM > 0.1) in DPCL intact Hi-C or POLR2A ChIA-PET data (enhancer-promoter loops, Fig. 3a, Extended Data Fig. 5d).

**Figure 3.**
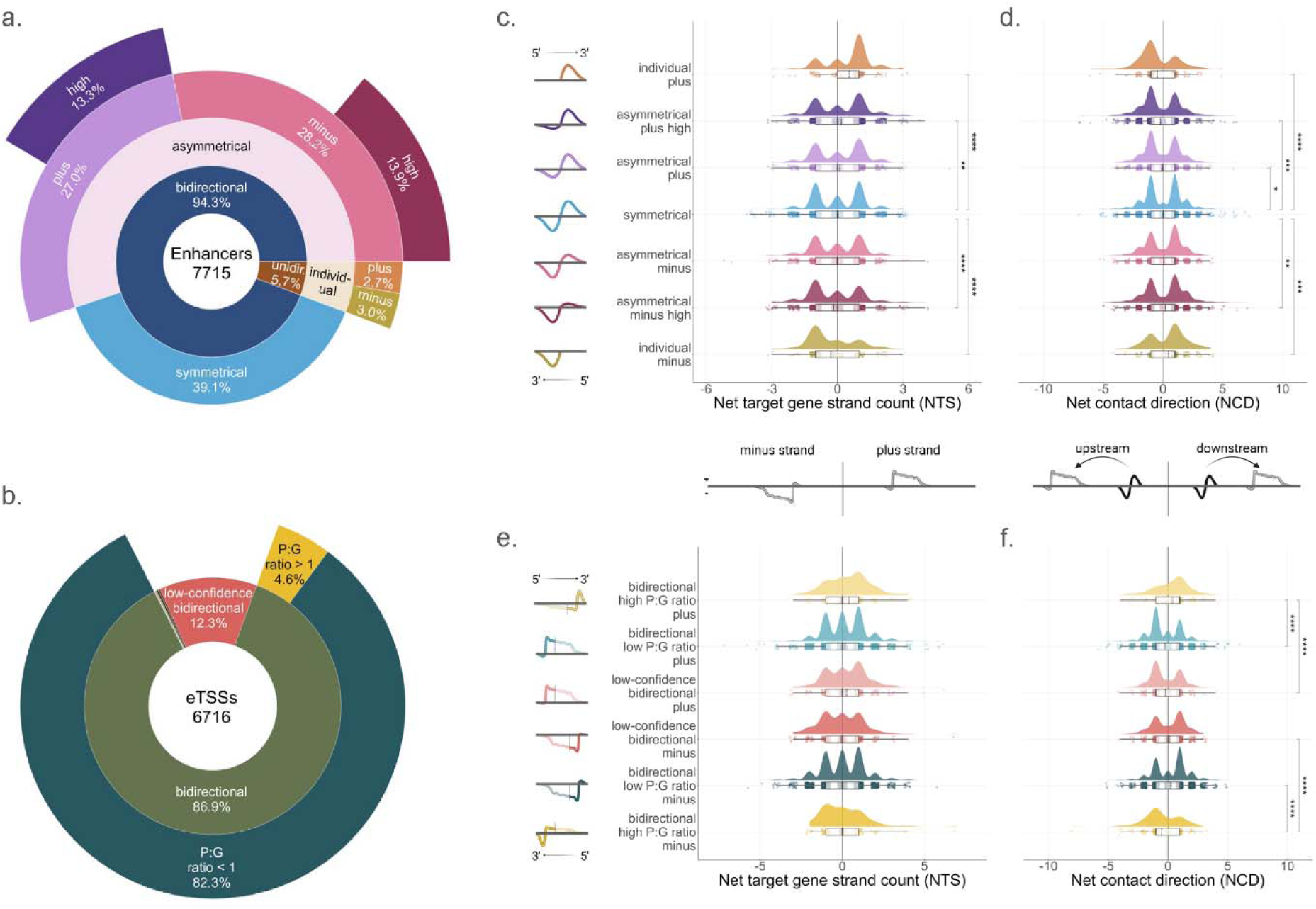
Enhancer-promoter and eTSS-promoter interaction patterns at asymmetrically transcribed lincRNAs. **a,b.** Class distributions for enhancers (**a**) and eTSSs (**b**) that interact with expressed gene promoters in either intact Hi-C or POLR2A ChIA-PET. Bidirectional eTSSs were also broken down according to their P:G RPKM ratio, designating eTSSs with higher PROMPT signal (P:G ratio > 1) and those with higher TSS-proximal gene signal (P:G ratio < 1). **c-f.** Distributions of nTS and nCD scores calculated for enhancers (**c-d**) and eTSSs (**e-f**) interacting with expressed genes via intact Hi-C loops. Schematics represent the patterns of enhancer or eTSS signal per group (left) and the interpretation of positive and negative nTS and nCD scores (middle), where the target gene (gray) is oriented relative to the DNA strand or about the regulatory element (black, e.g. enhancer/eTSS). For gene and PROMPT schematics, the sharp peak represents the PROMPT and the broad peak represents the gene. FDR adjusted p-values were determined from Wilcoxon signed-rank tests (*p_adj_ < 0.05, **p_adj_ < 0.01, ***p_adj_ < 0.001, ****p_adj_ < 0.0001).

eTSSs were defined from all assayed gene TSSs (N = 12939) by their overlap with proximal enhancer-like (pELS) cCREs and H3K4me1 histone signal peaks, and we retained those engaging in distal looping interactions with another expressed gene TSS (eTSS-promoter loops) as described for enhancers (Fig. 3b). Per cell line, fewer than 40% of assayed TSSs displayed any of these enhancer-like features and 1-12% had all three features (Extended Data Fig. 5e). eTSSs were primarily expressed in both the PROMPT region and gene (bidirectional TSSs) or enriched in the gene (low-confidence bidirectional TSS). Additionally, 4.6% of eTSSs had higher PROMPT expression (P:G ratio > 1, Fig. 3b, Extended Data Fig. 5f), highlighting that gene TSSs making distal interactions have varying expression patterns.

Although enhancer-promoter and eTSS-promoter pairwise interactions are independent of orientation^29^, we found that asymmetrical transcription of enhancers and eTSSs could be linked to patterns in their distal interactions. To investigate the relationship between enhancer and eTSS signal symmetry, we calculated two metrics, the net target strand (nTS) and net contact direction (nCD), to describe the looping interactions made by a regulatory element. The nTS describes the predominant target gene strand being contacted (N_plus-strand genes_ - N_minus-strand genes_), whereas the nCD describes the direction of the distal interaction from the enhancer or eTSS (N_downstream contacts_ - N_upstream contacts_). Assessing both metrics for each enhancer class and bidirectional eTSSs grouped by P:G ratio (Fig. 3b), we observed slight patterns in the interactions of asymmetrical regulatory elements. Asymmetrical enhancers preferentially interacted with target genes on the same strand as the more abundant eRNA (*p_adj._* < 0.01, Fig. 3c). They also have a preferred position relative to the gene, downstream for minus-strand and upstream for plus-strand enhancer classes (*p_adj._* < 0.05, Fig. 3d), which corresponds to transcription of the enhancer proceeding away from the gene. Interestingly, the patterns at enhancers where regions of higher signal correlate with nTS and nCD are not exactly reciprocated by eTSSs. A similar, but insignificant, nTS trend was detected for eTSSs, however the expected nCD pattern was observed relative to the gene strand, regardless of the eTSSs expression pattern (Fig. 3e-f). Importantly, these observations were consistent for contacts from both chromatin assays and for most cell lines individually (Extended Data Fig. 6).

nTS and nCD were not linearly correlated, although they may co-occur in a subset of asymmetrical elements (Extended Data Fig. 7a-b). Additionally, when evaluating the tendency of enhancers identified in two or more cell lines to follow these interaction patterns, we found that enhancers generally engage in similar net contacts in all cell lines, with no evidence that nCD correlates with changing expression patterns (Extended Data Fig. 7c-d). Together these results indicate that the transcription of some enhancer-like elements may relate to their orientation to and interactions with their target gene, however, these relationships do not exclusively dictate the directionality of enhancer transcription.

### Readthrough transcription is found downstream of most expressed genes

Termination of RNAPII can occur far beyond transcription end sites (TESs), resulting in RT transcripts, also known as downstream-of-gene (DoG)-containing RNAs. RT transcripts are detected in the homeostatic total RNA pool^6,35^, and are expressed in response to various stress conditions^36,37^. However, nascent RT transcripts have not been comprehensively characterized genome-wide in multiple human cell lines. We implemented an algorithm based on hidden Markov model (HMM) segmentation of Bru-seq data to identify regions of nascent homeostatic RT transcription downstream of genes (RTsegs), revealing that ∼75% of all expressed genes per cell line display nascent RT, ∼85% of which are pc-genes (Extended Data Fig. 8a-b). Collectively, all identified RT genes either had RTsegs in one or all cell lines, where 67.1% of ubiquitously expressed genes had RTsegs in all (Fig. 4a), suggesting that this is a general transcriptional phenomenon.

**Figure 4.**
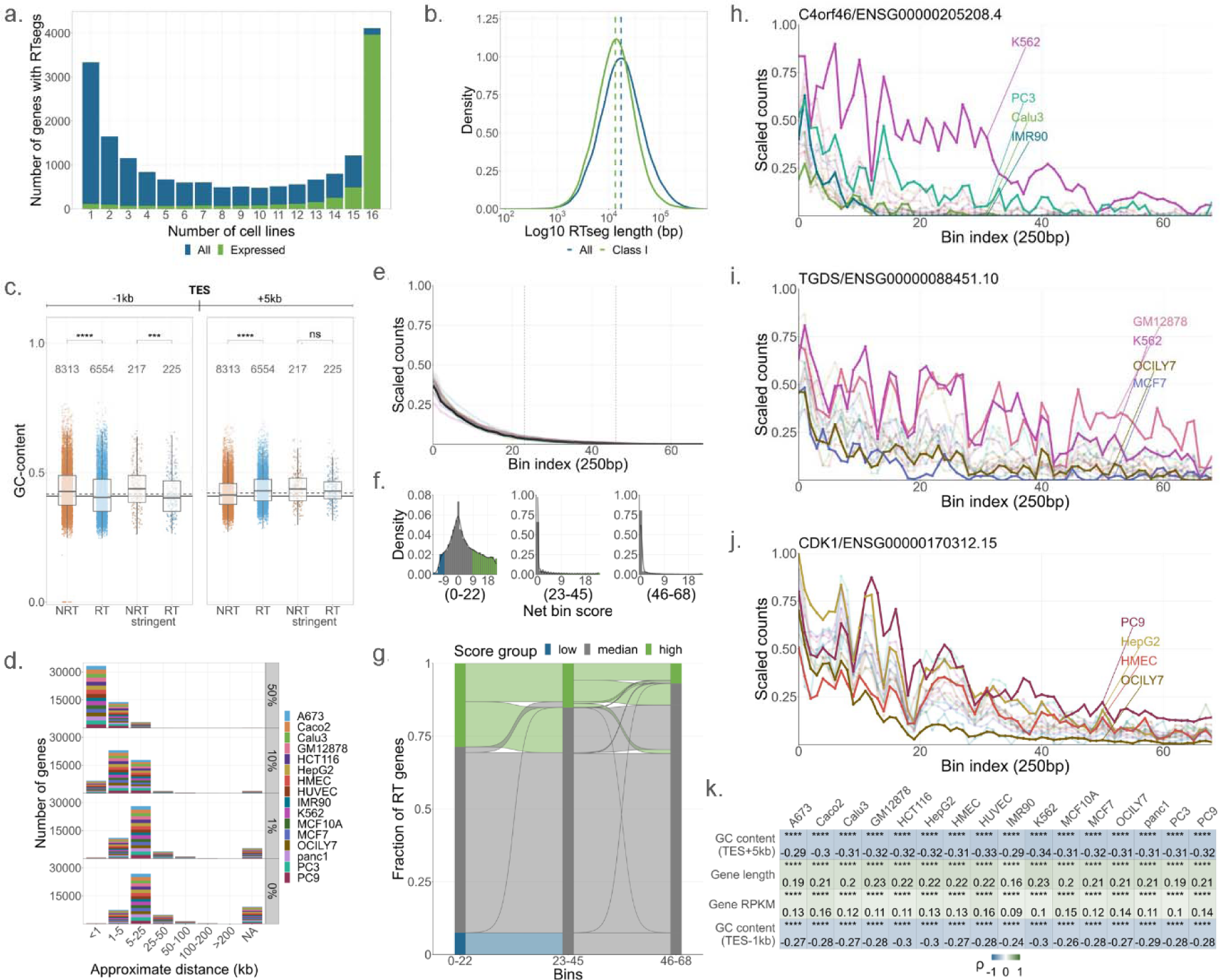
RT transcription downstream of expressed genes is ubiquitous and variable. **a.** The frequency of all (N = 18244) or expressed (RPKM > 0.25 in all cell lines, N = 5910) RT genes with an RTseg in 1-16 cell lines. **b.** Log10 length distributions of all and class I RTsegs, where the dashed line indicates the median of each distribution. **c.** The distribution of GC-content for all and stringent RT and NRT genes for the sequences 1 kb upstream (left) an 5 kb downstream (right) of gene TESs. Vertical lines represent the genome-wide GC-content^42^ (solid), and th median GC-content of all 1 kb or 5 kb regions (dashed). FDR adjusted p-values were determined from Wilcoxon tests. **d.** Distributions of genes with RT signal that reaches 0-50% of the 3’-gene signal within the approximate distance ranges shown. NA indicates the number of genes with RT signals that never reach the threshold, and th color shows the even distribution of genes per cell line. **e.** The median aggregated scaled signal of all 17.3 kb RT signals (black) and cell line specific RT signals. Scaled counts are calculated for 250 bp bins (Bin index) and vertical lines demarcate 23-bin sectors used downstream. **f,g.** The distribution of net bin scores (nBS) is shown for each sector (**f**). RT signals were grouped by their score–high (nBS ≥ 9) or low (nBS ≤ −9)–describing the magnitude and direction of their variability from the aggregate signal (**e**), and nBS trajectories between sectors are shown (**g**). **h-j.** Examples of scaled RT signal downstream of 3 genes (gene symbol/ENSEMBL ID) are shown for all cell lines. Selected cell lines are highlighted to display interesting patterns of variability. **k.** Representative correlations between RT signal length and gene features. Spearman’s rank correlation coefficients (ρ) and FDR adjusted p-values ar displayed. Adjusted p-values (**c**, **k**) are denoted as: *p_adj_ < 0.05, **p_adj_ < 0.01, ***p_adj_ < 0.001, ****p_adj_ < 0.0001.

Nascent, homeostatic RT events were found to occur similarly to known stress-induced and homeostatic RT transcription^6,35,36^. A wide range of RTseg lengths was observed, with a median of 17.3 kb (Fig. 4b, Extended Data Fig. 8c). Because RTsegs can overlap downstream genes (see Methods), each was classified accordingly into 4 main groups, where class I segments do not overlap a gene on the same strand and classes II-IV describe different patterns of gene overlap (Extended Data Fig. 8d). Paralleling all segments, class I RTsegs displayed a wide range of lengths (median = 13.5 kb, Fig. 4b, Extended Data Fig. 8c). Comparing all RT genes and genes with no RTseg identified in any cell line (NRT), we found that RT genes: 1.) tended to be slightly more proximal to their nearest downstream neighbor^36^ than NRT genes (Extended Data Fig. 8e), 2.) were enriched for the elongation mark H3K36me2^35,36,38^, both downstream (TES+5kb) and upstream of their TES (TES-1kb, Extended Data Fig. 8f-g), and 3.) had a lower 3’-end GC-content than NRT genes^35^, but a slightly higher GC-content downstream (TES+5kb, Fig. 4c). These results were validated with stringent RT/NRT and pc-gene subsets (see Methods), substantiating the similarities between nascent RTsegs and previously identified RT events.

### Readthrough transcription varies among genes and cell lines

Rather than terminate in a defined manner, readthrough signal tends to taper off gradually^36,39–41^. To better understand this behavior, we scaled RT signal (downstream of TESs) to gene signal (upstream of TESs). We assessed the approximate distance required for each RT signal to reach 50, 10, 1, and 0% of the 3’-gene signal (Fig. 4d), and found that within 1 kb of the TES, the RT signal for most genes reduced below 50%, and typically decreased to 0 within 25 kb, aligning with the median RT signal and RTseg lengths (∼11-17 kb, Extended Data Fig. 9a).

The extent of RT signal variability was assessed by identifying genes displaying signal patterns differing from the median aggregated signal of all genes (Fig. 4e). Scaled RT signals were obtained for a fixed 17.3 kb region downstream of RT genes and normalized to the aggregate signal per cell line. RT signal regions were assigned a net bin score, describing the degree of variation between the RT signal and aggregated signal. Based on their score, RT signals were designated as having significantly “high” or “low” signal relative to the median aggregated signal for each third of the 17.3 kb region (23 bin or 5.8 kb sectors, Fig. 4f) due to the larger degree of variability seen in the first ∼5 kb downstream of the TES (Extended Data Fig. 9b-d). RT signals that were significantly higher than the aggregate signal were found in each sector, including 5.9% that were higher throughout the entire 17.3 kb region (Fig. 4g). Negative scores were present in the first 5.8 kb only, but interestingly a handful of these initially low signals ended up above the aggregate signal downstream (Fig. 4g). Importantly, signal variability was found for individual genes across cell lines, with differences in relative RT signal amounts (Fig. 4h-j, Extended Data Fig. 9e-g) and occasionally distinct patterns of signal decay (Fig. 4j), suggesting that there may be gene and cell line specific regulation of RT termination.

RT termination variability was examined by correlating RT signal length with features shown to distinguish RT and NRT genes and basic gene properties that can impact RNAPII elongation (e.g. length, RPKM, co-transcriptional splicing index^43^). A weak negative correlation between RT length and GC-content was found (TES-1kb, TES+5kb), with other parameters exhibiting negligible correlations for all and class I RT regions (Fig. 4k, Extended Data Fig. 10a-b). Despite the low correlation with RT length, RT genes with higher scaled signal downstream typically had lower GC-content at their 3’-ends (TES-1kb, TES+5kb, Extended Data Fig. 10c-d), suggesting that GC-content may influence RT transcription.

RT genes had slightly higher GC-content downstream (TES+5kb) than NRT genes (Fig. 4c). Upon surveying all possible hexamers downstream of RT and NRT genes, we found an enrichment of GC-rich k-mers downstream of RT genes that increased slightly at regions further from the TES, although GC-content did not increase concordantly (Extended Data Fig. 10e-f). Interestingly, a similar pattern in GC-content was found upstream of the TSS, where the region beyond the peak PROMPT signal (TSS+500bp, Extended Data Fig. 2a) was more GC-rich, contrasting the pattern in genes (Extended Data Fig. 2h). This suggests that GC-content or G-rich motifs may play a general role in termination of transcription outside of genes, although these sequence features do not explain all variability seen in RT signal between genes and cell lines.

## Discussion

Canonical gene transcription gives rise to PROMPTs, eRNAs, and RT transcripts, however, little is known about their regulation and functions. Using the diverse transcriptomic and functional genomic data provided by the ENCODE4 DPCL, we have explored the characteristics of these co-products of transcription in 16 human cell lines and described sequence and chromatin features that correlate with their expression.

Initiation at pc-gene TSSs is highly regulated by proximal and distal elements, yet both PROMPTs and eRNAs can arise from transcription at NDRs. We found varying transcriptional patterns flanking a TSS and determined that regulation of PROMPT transcription was largely independent of the associated gene. Differential expression patterns of PROMPTs and genes dynamically correlated with histone modifications associated with transcription, consistent with previous studies^25,44,45^. Despite overall lower aggregate signal over PROMPT regions versus genes, a subset of expressed PROMPTs were highly enriched for H3K79me2 signal, which suggests that this modification could be associated with more specific regulation of PROMPT transcription. Due to methodological limitations (Supplementary Note 4), we were unable to accurately assess PROMPT length, but H3K79me2 may demarcate elongating PROMPTs with distinct functions or properties. Interestingly, H3K79me2 has been implicated in maintaining distal enhancer-promoter interactions for some enhancers^46^, and may play a similar role at TSSs to facilitate various regulatory element interactions (enhancer-promoter, eTSS-promoter). In line with this, we found that enhancers and eTSSs are enriched for H3K79me2 over the transcribed region (Extended Data Fig. 5g).

Highly asymmetrical enhancers and eTSSs engaged in putative chromatin contacts at levels proportional to their prevalence in our dataset, suggesting that symmetrical bidirectional transcription is not required for regulatory activity. Strikingly, asymmetrically transcribed enhancers often interacted with an inferred target gene on the same strand and with a preferred position relative to the gene, where the enhancer was transcribed away from the gene. eTSS-promoter interactions were distinct in their patterns, with less clear nTS trends and an nCD preference that mirrored enhancer-promoter interactions, when assessed relative to the gene strand. A current model describing enhancer-promoter interactions predicts that dynamic phase separated hubs are formed by high local concentrations of transcription-associated proteins that facilitate transcriptional bursts at both elements^47–49^ (‘hub’ model). Consequently, it has been suggested that the rate of initiation and productive elongation of enhancers and promoters may influence their mobility and contact frequencies within a domain^49,50^. Additionally, negative torsional tension within topological domains containing transcribed enhancers and promoters may influence their interactions^51–53^. Thus, the link between transcription directionality at these elements relative to loop orientation (nCD), coupled with the observation that at eTSS-promoter interactions the elongating element (i.e. the gene) corresponds to nCD, may align with the hypothesis that transcription regulates these interactions.

Many enhancers and eTSS do not show the described trends, however, nor do these interaction patterns explain cell line specific differences in transcription without changes in nCD (Extended Data Fig. 7c-d). This variability suggests that other factors, such as histone modifications or TF binding^11^, are involved in the regulation of transcription directionality at these elements. Recent studies suggested that eRNA and PROMPT sequence complementarity may contribute to enhancer-promoter interaction selectivity via their direct association^54^, perhaps with histone modifications, such as H3K79me2^55^, contributing to the stabilization of these RNA species and promoting their interaction. In line with the hub model, both enhancers and eTSSs can engage in interactions with multiple target genes and regulatory elements^50^ (Extended Data Fig. 7e), although these interactions are unlikely to occur simultaneously in individual cells^56^ where enhancer transcription is unidirectional^57^. Despite bulk data being ill-suited to the assessment of multiplex chromatin interactions, these observations display the complexities inherent to distal enhancer-promoter interaction patterns and highlight the utility of more targeted methods of evaluating genome-wide distal interaction patterns alongside nascent transcription^58^.

Readthrough transcription is a known phenomenon that results in RT transcripts that are stably expressed under homeostatic conditions and induced upon cellular stress for some genes. We showed that nascent, homeostatic RT transcription is widespread, occurring downstream of at least 75% of expressed genes, which is higher than current estimates based on RT transcripts found in the total RNA pool^35,36^. This suggests that while RT transcription occurs downstream of most genes, RT transcripts may be dynamically regulated.

By profiling nascent RNA signal downstream of RT genes, we found that most RT regions had less than 50% of the 3’-gene signal remaining immediately downstream of the TES (Fig. 4e), consistent with past observations^36,39–41^. This may indicate that most polymerases terminate efficiently but at various sites beyond the TES, however, nascent long-read sequencing will be required to determine the frequency of these events^40^. Nevertheless, we report instances of higher and lower RT signal relative to the gene, and variable RT signal patterns between cell lines, suggesting cell line specific RT termination for some genes.

The nascent RT genes we identified shared many sequence and chromatin features with known RT parent genes^35–37^. However, we found little to no correlation between these features and the distance downstream of the TES that RNAPII can travel, aside from a mild negative correlation with GC-content. While the 3’ GC-content of RT genes is lower than non-RT, corresponding with the hypothesis that RNAPII speed may influence RT transcription^35^, we noted a slight increase in GC-content and enrichment of GC-rich sequences in the 5 kb downstream of RT genes. Analogously, an increase in GC-content was detected in PROMPT regions after the peak of RNA signal (TSS+500bp). Increased GC-content and/or GC-rich sequences may be involved in slowing RNAPII to promote termination^59–61^. Given the correlation between lower GC-content and increased RT signal, transition from lower to higher GC-content may be a general mechanism to aid in termination, whereas RT transcription is regulated by dynamic termination factors, like CTD phosphorylation, Xrn2, BRD4, mediator, or integrator^62–66^.

Our data describing PROMPT, eRNA, and RT transcription, generated during ENCODE4, allowed for the exploration of regulatory patterns associated with their expression. Accompanying the potentially specific regulation of transcription-associated lincRNAs, there may also be general mechanisms that modulate their expression. For example, integrator is involved in 3’-end processing of pc-genes^66^, as well as both PROMPTs and eRNAs^67^. CTD tyrosine1 phosphorylation was previously shown to influence termination of both PROMPTs and genes and plays a role in RNAPII initiation or pausing at both promoters and enhancers^62^. These parallels suggest general regulatory mechanisms of intergenic, non-coding RNAs, though, further investigation will be required to delineate the different modes of transcription that occur within and outside of genes.

## Data Availability Statement

The datasets analyzed in the current study are products of the most recent ENCODE consortium (ENCODE4) and are available on the ENCODE project portal^68–70^ (encodeproject.org). Deeply profiled cell line (DPCL) sample accessions, ENCODE histone ChIP-seq accessions, ENCODE4 cCRE accessions, and all other file accessions and metadata are detailed in Supplementary Data File 1.

## Code Availability Statement

The ENCODE Data Coordinating Center (DCC) have developed Uniform Processing Pipelines for most major assay types generated by the project^71^. Details about the DCC Uniform Processing Pipelines can be found on the ENCODE portal (https://www.encodeproject.org/pipelines/) and code is available on the ENCODE DCC GitHub (https://github.com/ENCODE-DCC). More information regarding the specific mapping pipelines used for each assay can be found in the Supplementary Methods (section 2). Custom code used to perform analyses in this publication is available on GitHub (https://github.com/LjungmanLab/ENCODE-DPCL-paper). A list of all software and versions used to perform these analyses can be found in Supplementary Table 2.

## Methods

### Cell lines, sequencing, and data mapping

DPCL cell lines were grown and collected in two batches (N = 2 biosamples) for all assays together according to standard methods established for each cell line and then distributed to participating ENCODE consortium labs for completion of experimental protocols and sequencing. Growth conditions of each cell line can be found in Supplementary Table 1 and details on cell collection and experimental protocols can be found in the Supplementary Methods (section 1). In addition, experimental guidelines and data standards for most assays can be found on the ENCODE portal (https://www.encodeproject.org/data-standards/).

Sequencing parameters for each sample are available on the ENCODE portal, and are briefly detailed in the Supplementary Methods (section 2). All data was aligned to GRCh38 and gene-centric analyses used the GENCODE annotation, version 29^72^ (v29, https://www.GENCODEgenes.org/human/release_29.html). The ENCODE Data Coordinating Center (DCC) has developed Uniform Processing Pipelines for most major assay types generated by the consortium^71^. Details about the DCC Uniform Processing Pipelines can be found on the ENCODE portal (https://www.encodeproject.org/pipelines/) and code is available on the ENCODE DCC GitHub (https://github.com/ENCODE-DCC). Not all data processing pipelines are currently among the Uniform Processing Pipelines developed by the DCC, but these are described in the Supplementary Methods (section 2) in more detail, along with additional details for all DPCL assays.

### Genome coverage calculations

To calculate the proportion of the genome covered by at least one uniquely mapping sequencing read for each assay and cell line, we generated unnormalized, stranded bigwigs from alignments using deeptools^73^ bamCoverage for each sample using the following parameters (see Supplementary Data File 2 for file and parameter details):

--normalizeUsing None

--minMappingQuality 255 (where applicable)

--filterRNAstrand forward/reverse

--binSize 1

--skipNAs

We then merged bigwigs per strand using UCSC utilities bigWigMerge for each application that assessed the coverage of more than one sample, including cell line replicate samples (Extended Data Fig. 1a), all samples per assay (Fig. 1a,c; Extended Data Fig. 1b-c), and all DPCL samples (Fig. 1b). The resulting bedgraph files were then sorted (UCSC utilities bedSort) and reverted to bigwigs (UCSC utilities bedGraphToBigWig). Coverages were calculated for each of the canonical chromosomes (1-22, X, Y) per strand from merged bigwigs using the pyBigWig (deeptools) bw.stats(type=“coverage”), and the total bases covered on both strands was calculated for all chromosomes to yield a signal fraction of bases covered per bigwig.

For genic, intergenic, and exonic coverage calculations, merged bigwigs were generated as described above for Bru-seq, BruUV-seq, and total RNA-seq using only the 24 libraries for which we had data in all three assays (8 samples are missing from DPCL total RNA-seq), to keep these data as consistent as possible. Similarly, these data are only shown for the 8 cell lines (16 libraries) which had two biological replicates in all three assays (Extended Data Fig. 1b-c). To calculate the fractions of bases covered in the genic versus intergenic regions of the genome, we first obtained an annotation of these regions (.bed format). We merged (bedtools^74^ merge) gene boundaries from the GENCODE v29 basic annotation in a stranded fashion to establish the genic region, and then assigned the remaining regions, up to the length of each chromosome (GRCh38), as intergenic. To obtain the exonic regions, we first created a collapse exon annotation (GENCODE v29 basic) per gene using a custom python script (available on GitHub) and then merged any overlapping exonic regions on the same strand with bedtools as described above for genes. The coverages in each of these regions were calculated with pyBigWig.stats(type=“coverage”) as described above and the total bases covered was calculated per region, yielding 3 coverage values per bigwig (genic, intergenic, exonic).

Finally, genic versus intergenic counts were calculated for each bigwig using featureCounts^75^ (subread) using the following commands:

-F SAF

-p (excluded for total RNA-seq)

-Q 255

-s 2

--countReadPairs (excluded for total RNA-seq)

Counts were then summed per region, and the fraction of total counts was calculated using the total number of uniquely mapping reads per bigwig (sum of all region counts). To compare cell lines, Pearson’s correlations (r) were calculated from a counts matrix and hierarchical clustering was performed using pheatmap(clustering_distance_rows=“correlation”, clustering_distance_cols=“correlation”, clustering_method = “complete”, cluster_rows = T, cluster_cols = T). Importantly, this method of counting is different than the method used from all other count-based analyses performed with Bru-seq or BruUV-seq data (Supplementary Methods, section 2.2). However, these two methods were compared for Bru-seq data and were found to be well-correlated (Supplementary Fig. 1a-b). All data described in this section can be found in Supplementary Data File 2.

### Genic Transcription Start Site (TSS) curation for PROMPT characterization

An annotation-dependent method was used to characterize PROMPT regions. A fixed 2 kb divergent region upstream of an annotated gene TSS, that represented the majority of the PROMPT signal, was used to characterize the PROMPTs (Extended Data Fig. 2a). Annotated protein-coding genes and transcripts in the GRCh38 GENCODE v29 basic annotation were used for this analysis, resulting in multiple TSSs per gene. Using bedtools merge -s, TSSs belonging to the same gene that were within 2 kb of each other were merged into a single unit, with the most upstream coordinate serving as the TSS. Genes and transcripts greater than 2 kb, along with any merged transcript/gene greater than 2 kb, were kept. TSSs having an overlapping annotated feature (any biotype, GENCODE v29 basic annotation) in 2 kb regions up- and downstream of the TSS on the antisense strand were removed from the analysis. Mitochondrial genes were removed from this analysis. To remove RNA signals from alternate sources, TSSs were removed if the fixed 2 kb PROMPT region overlapped a readthrough segment originating from an upstream gene in any cell line (readthrough regions described in Methods: RT segment identification). However, PROMPT regions that overlapped a readthrough segment but had a BruUV-seq peak detected by our peak calling algorithm (see Methods: BruUV-seq peak calling and eRNA identification) in all cell lines were kept. Finally, any TSS within merged regions larger than 2 kb produced during the TSS merging step were manually evaluated to determine unambiguous initiating TSSs in BruUV-seq data to be retained. A resulting total of 12939 PROMPT regions/TSSs were generated for downstream analyses.

### TSS category generation

BruUV-seq and Bru-seq nascent RNA assays were used to classify the 12939 TSSs. We utilized a total of 5 metrics, thereby generating 8 working categories and 1 ambiguous category (Supplementary Table 3). PROMPT counts and RPKM (upstream divergent TSS - 2 kb region) were obtained from BruUV-seq data. For higher genic signal confidence, we used both BruUV-seq and Bru-seq data to get gene counts and RPKM values for a comparable TSS-proximal 2 kb region (TSS + 2 kb). Per cell line, replicate expression values for all regions of interest were averaged for downstream signal analysis. For classifying the signal around gene TSSs, we considered only expression (RPKM) for the PROMPTs, whereas for a gene, we considered both the expression (RPKM) of the TSS-proximal 2 kb region as well as its enrichment over the preceding 2 kb region on the same sense strand (TSS enrichment ratio = TSS+2kb RPKM / TSS-2kb RPKM). The criteria thresholds were set as the following: RPKM threshold = 0.1, TSS enrichment = 2. A parameter tag of “+” was indicated if that region’s metric was above the criteria threshold or “-” if it was equal to or below the threshold. We required the two genic parameter tags, i.e. the TSS-proximal gene RPKM and the TSS enrichment, to be the same per assay. Since the ambiguous category was predominantly a result of the two genic parameters being nonconcordant per assay, it was excluded from all downstream analyses. Additional rationale and validation methods are expanded upon in the Supplementary Methods (section 5.2). This resulted in total TSSs for all cell lines = 207024, total unambiguous TSSs for all cell lines = 132799.

### PROMPT: gene ratio calculation

PROMPT:Gene (P:G) expression ratios were calculated using BruUV-seq RPKMs in the 2 kb PROMPT and the TSS-proximal gene regions (P:G expression ratio = divergent TSS-2kb PROMPT RPKM / TSS+2kb gene RPKM). Log10 transformed ratio values were used for downstream analyses.

### Histone post-translational modification signal determination and ratio calculation

We explored the relationship between the TSS-proximal P:G expression and selected histone post-translational modifications namely H3K9ac, H3K27ac, H3K4me1, H3K4me2, H3K4me3, H3K27me3, H3K36me3, H3K79me2 for the cell lines that had data for all the modifications (10/16). TSS-proximal histone signal (counts) was obtained from signal p-value bigwigs for the various histone post-translational modifications (Supplementary Data File 1) using deeptools multiBigwigSummary for gene (TSS+1kb) and PROMPT regions (TSS-1.1kb, 100 bp is an average estimate of the Nucleosome-depleted region (NDR) for the TSSs in this analysis). These regions were determined from the histone aggregate plots (Extended Data Fig. 3b). Normalized counts (CPM) were generated and the histone P:G ratios were calculated as follows: Histone P:G ratio = (TSS-1.1kb PROMPT CPM / TSS + 1 kb gene CPM). Log10 transformed ratio values were used for downstream analyses.

Regions of histone signal enrichment (peak overlaps) were determined by overlapping signal peaks with gene (TSS+1kb) and PROMPT regions (TSS-1.1kb) using bedtools intersect with default overlap fraction options. The histone signal and enrichment/peak overlap groups were substantially in agreement with each other for the PROMPT regions across all cell lines (Supplementary Fig. 7a-e, 8a-c). Wilcoxon-signed rank tests were performed between the signal enrichment groups (No signal enrichment and Signal enrichment) per histone modification in the gene and PROMPT regions using the compare_means function (method = “wilcox.test”, p.adjust.method = “BH”, R/rstatix). False Discovery Rate (FDR) adjusted p-values were calculated from p-values and represented by ^ns^p_adj_ > 0.05, *p_adj_ < 0.05, **p_adj_ < 0.01, ***p_adj_ < 0.001, ****p_adj_ < 0.0001. Information about all the sequence and chromatin parameters that we tested, along with additional validation methods are expanded upon in the Supplementary Methods (section 5.4).

### P:G ratio correlation analysis

Spearman correlation coefficients were calculated between log10 transformed P:G expression and histone signal ratios using the cor.test function (method = “Spearman”, exact = FALSE) in R. FDR adjusted p-values were calculated from p-values in R using the p.adjust function (method=”BH”), and represented by ^ns^p_adj_ > 0.05, *p_adj_ < 0.05, **p_adj_ < 0.01, ***p_adj_ < 0.001, ****p_adj_ < 0.0001.

### Enhancer-like gene TSS (eTSS) identification

The 12939 TSSs were further evaluated for active enhancer-like features. TSSs were designated as eTSSs if they overlapped an H3K4me1 signal peak (Supplementary Data File 1) and a proximal enhancer-like candidate cis-Regulatory Element^33^ (pELS cCRE, determined by high DNase and H3K27ac signal, with low H3K4me3) in 500 bp regions up- and downstream of the TSS using bedtools intersect with default overlap fraction options. These requirements align with what has been shown previously pertaining to enrichment of histone or chromatin features for genic promoters that have enhancer-like regulatory potential^76,77^. The cell lines lacking H3K4me1 or pELS data (Calu3, MCF10A, HUVEC and HMEC) were removed from this analysis. Additionally, we required the TSSs to make at least one contact with an expressed protein-coding or lincRNA gene (TSS-proximal 2 kb BruUV-seq RPKM > 0.1). These contacts were determined from using bedtools pairToBed to overlap loop anchors obtained from intact Hi-C or POLR2A ChIA-PET data with regions 1kb up- and downstream of the two sets of TSSs. Self-interacting promoters were removed and reciprocal promoter-promoter interactions were retained.

### BruUV-seq peak calling and eRNA identification

To identify regions of enhancer transcription, i.e. eRNA signal, from BruUV-seq data, we first called peaks of BruUV-seq signal (BruUV-peaks) in an annotation-independent manner using the MACS2^32^ peak calling algorithm. For each library, properly oriented read pairs were obtained per strand using sambamba^78^, and narrow peaks were called on the plus (+) and minus (-) strands separately with the subcommand ‘callpeak’ and the following parameters: --p 0.01, --keep-dup 1, --call-summits. Peak calling was similarly carried out using alignments from all replicates to generate a pooled peak file (--p 0.01 --keep-dup 1). Consensus peaks, denoted as replicated peaks, are then derived from the BruUV-seq pooled replicated peaks using single library peaks for filtering. Peaks were considered replicated if they overlapped a peak summit called in each individual replicate library. These signal peaks were identified for all cell lines and validated using complementary data (Supplementary Methods, section 6.2). Since we expect BruUV-seq signal to be concentrated at both stable and highly unstable RNA species (Fig. 1e), this method can be used to identify transcription initiation at various RNA species throughout the genome without the requirement of an annotation, meaning this could also be used to identify potentially novel regions of transcription. For the purposes of this study, BruUV-peaks were then subjected to filtering based on their intersection (bedtools) with putative distal enhancer elements (dELS) from the ENCODE4 cCREs^33^. In addition, we removed peaks overlapping proximal enhancer cCREs (pELS) due to possible ambiguity in the origin of RNA signal that is proximal to a TSS. These cCRE annotations were specific to each cell line, thus cell lines without enhancer-like cCRE annotations (HMEC, HUVEC) were excluded from this analysis, meaning data for only 14 cell lines is presented here. The resultant peaks are BruUV enhancer peaks that were then subjected to downstream signal classification.

Using bedtools closest (-S -D a -nonamecheck) BruUV enhancer peaks were paired with the nearest divergent peak, and peak pairs with 5’-ends within 600 bp of one another were designated as arising from bidirectional enhancers. Bidirectional peak pairs and individual peaks with no divergent pair were then grouped by their relative divergent signal. RPKMs and RPKM ratios were calculated for all peaks and used for their classification (Extended Data Fig. 5a). For all cell lines, the mean and standard deviation (stdev) of bidirectional peak-to-peak ratio (plus strand peak/minus strand peak) distributions were calculated, and the overall mean values across all cell lines (mean = 0.0032, stdev = 0.4434) were used to divide the distribution into groups based on their signal symmetry. A peak pair was designated as asymmetrical if it had ratio greater than 1.678 or less than 0.605, i.e. 0.5 stdev away from the mean, with highly asymmetrical classes demarcated by RPKM ratios greater than 1 stdev from the mean (Supplementary Data File 4). The resultant class distribution is similar to previously published data^28^, however due to our more inclusive enhancer subset, it reflects higher proportions of asymmetrically transcribed enhancers. For unpaired (individual) peaks, the RPKM ratio of the peak and a divergent region of the same size were calculated, and peaks with no signal (RPKM ratio = 0) in the divergent direction were determined to arise from unidirectional enhancers. All individual peaks with any BruUV-seq signal (RPKM ratio > 0) were designated as low-confidence bidirectional enhancers and were excluded from downstream analyses. Low-confidence bidirectional enhancers were excluded primarily due to the incompatibility of their RPKM ratios and those of bidirectional peak-pairs, preventing them from being classified in the same manner. Furthermore, it is more likely for these unpaired peaks to result from regions of noisier BruUV-seq signal, e.g. within a gene body or at super-enhancers, where it is difficult to resolve peaks of nascent RNA signal arising from a single enhancer element.

### Enhancer-promoter and eTSS-promoter looping analysis (nTS/nCD)

Enhancer-promoter and eTSS-promoter loops were obtained by overlapping (bedtools pairToBed) identified enhancer-like elements and expressed protein-coding or lincRNA gene TSSs with intact Hi-C and POLR2A ChIA-PET loop anchors (see Supplementary Methods, sections 2.3 and 2.5). Loops in either assay that overlapped an enhancer-like element at one anchor and an expressed protein-coding or lincRNA gene at the other anchor were retained for downstream analyses. Gene expression was assessed in the BruUV-seq RPKM of the 2 kb region immediately downstream of the TSS (RPKM > 0.1) to mirror thresholds used to establish TSS-proximal gene expression in the PROMPT analysis. For each enhancer-like element, we collected information about all gene contacts that it made, summing the counts of genes on each strand and chromatin contacts in each direction, and used this information to calculate its net target strand (nTS = N_plus-strand genes_ - N_minus-strand genes_) and net contact direction (nCD = N_downstream contacts_ - N_upstream contacts_) indices. The direction of contacts (upstream and downstream) used in the nCD calculation was established using the genomic coordinates of the enhancer and target gene, where a target gene with a larger genomic coordinate than the enhancer was considered a downstream contact, and vice versa (i.e. the direction reported is relative to the enhancer). For loop anchors that were found to overlap multiple target genes–typically the result of genes sharing or having very close TSSs–we counted each target gene uniquely when considering the strand but counted the loop uniquely when considering the contact direction. Importantly, it is possible that reciprocal eTSS-promoter interactions are present if both TSSs are enhancer-like and both genes are expressed. Details on validation of these results with an orthogonal technique and the analysis of nCD at enhancer consensus loci can be found in Supplementary Methods section 6.2-3.

### RT segment (RTseg) identification

To infer regions of readthrough transcription, we performed genome segmentation using a previously described Hidden Markov model (HMM) with 10 logarithmically distributed output states^79–81^. Here we used 250bp binned RPKM values as an input to this model, which establishes the inferred transcription states of genome segments (HMM index 0-9), and excluded segments with an HMM index less than three^81^. We then merged adjacent segments with equal or decreasing HMM indices and intersected these merged segments with gene transcription end sites (TESs). Using the GENCODE v29 annotation, one TES was obtained per gene by selecting the 3’-most end site from all annotated gene isoforms. Each segment that overlapped multiple TESs was assigned to the most upstream TES that it overlapped, and the 5’-coordinate of all segments was defined by the TES of the associated gene. These segments were considered possible readthrough segments for their associated gene and were subject to further filtering. Readthrough segments were not reported for genes that were less than 1kb in length or were lowly expressed (RPKM < 0.25). These length and expression cutoffs were taken due to the inability to accurately resolve Bru-seq signal and RT segments at small and lowly expressed genes. Coverage over readthrough segments was obtained, and all segments with fewer than 10 counts were not reported. The remaining segments were reported as readthrough segments (RTsegs) and were subject to downstream classification and refinement based on their overlaps with annotated genes (Supplementary Methods, section 7.1). RTsegs were identified for each sample individually but were merged to create a cell line specific annotation of RT regions for most analyses. To merge replicates, the longest segment from either sample was retained per gene, relying on the assumption that each replicate would have identical RT potential downstream of a particular gene (Supplementary Methods, section 7.3). These data can be found in Supplementary Data File 5.

### RT versus NRT gene definitions

Due to the restriction of our analysis to genes that are greater than 1 kb in length we assayed a total of 38006 genes (GENCODE v29) for read through. RT genes were identified as genes that were assigned an RTseg in at least one cell line (18244) and NRT genes were not assigned an RTseg in any cell line (19762). Given that these analyses are independent of the length of a RTseg, genes with ambiguous RTseg classifications are included here (Supplementary Methods, section 7.1). However, to remove the possibility that technical ambiguity could confound our results, we also obtained a subset of stringent NRT genes (N = 239). Stringent NRT genes: 1.) were required to be expressed (RPKM > 0.25) in at least one cell line, 2.) have little to no RNA signal in the 5 kb downstream of their TES (<10 Bru-seq reads), 3.) do not have another gene within 5 kb downstream, and 4.) their TESs are not overlapped by an RTseg coming from an upstream gene on the same strand. Gene expression was not enforced in all cell lines due to the presence of very few ubiquitously expressed genes that met all other listed requirements. Stringent RT genes are a subset of 239 genes where: 1.) at least 1 cell line had a class I RTseg (does not overlap a downstream gene on the same strand), 2.) no cell lines have ambiguous RTseg classifications (Supplementary Table 3), and 3.) do not have another gene downstream within 5kb. The distribution of biotypes between the two subsets–and the overall distribution of RT gene biotypes–was kept as similar as possible to remove the potential for differences between coding or non-coding gene features to influence our results in either subset (Supplementary Fig. 13a). The distribution of gene lengths between the two stringent subsets were also confirmed to be similar (Supplementary Fig. 13b). In addition to the stringent subset, all results were verified using a subset of only protein-coding genes due to the imbalance of biotypes in the full RT/NRT gene sets and to more closely resemble the results of previous studies which have typically attributed RT transcripts to protein-coding genes exclusively. These gene lists were used for all analyses comparing nascent RT transcription to known steady-state homeostatic or stress-induced RT transcripts/DoGs (Supplementary Methods, section 7.4-6).

### Scaled RT signal calculation

Nascent readthrough signal was evaluated from scaled binned counts (250 bp bins) downstream of an RT gene TES. These binned counts were either obtained for the entire RTseg identified in our segmentation analysis, or for a fixed 17377 bp region downstream of RT genes. RT signal length was evaluated using binned counts for the full-length segments, and comparisons between RT signal patterns found across cell lines were performed using the fixed 17.3 kb regions. These two specific use cases are described in the sections below, and further explanation on the rationale of distinguishing these methods can be found in the Supplementary Methods (section 7.7). For all RT genes, the binned counts in the RT region were scaled to the signal at the end of the gene to allow comparison of RT signal patterns between genes and cell lines. To do this, we obtained the counts of four 250 bp bins upstream of the TES (∼1 kb into the gene body) and divided the binned counts in the readthrough region by the maximum binned count value at the end of the gene. Because multi-isoform genes, particularly those with multiple annotated TESs, could result in inaccurate measurements of the end-of-gene signal used for scaling, we restricted this analysis to single-isoform genes and genes where all annotated TESs are within 250 bp (1 bin size) of each other. These scaled RT signals were then further filtered for downstream analyses as needed.

### Approximate distances traveled by RT signal

Scaled counts from full-length RTsegs were used to calculate approximate distances or lengths to a particular signal level (number of bins * bin length) for most RT genes. RT genes were excluded from these analyses if their RTseg length was less than 250 bp or if they had RTsegs with ambiguous classifications (Supplementary Methods, section 7.1). The number of bins to each signal level was determined by setting a scaled signal threshold (e.g. 0, 0.01, 0.1, 0.5) and counting how many bins it took until the signal was maintained at or below that threshold for four consecutive bins (1 kb). The distribution of approximate RT signal lengths to zero scaled counts was reported (Extended Data Fig. 9a) and compared with RTseg lengths (Supplementary Methods, section 7.8) and the approximate distance to different signal levels (50, 10, 1, 0% of the gene signal) allowed quantification of the different signal trajectories downstream of RT genes (Fig. 4d). A threshold of zero counts was considered appropriate for these analyses based on the use of segmentation data to define the assayed regions before counting, thus we do not expect to be capturing noise.

### RT signal length correlations

The RT signal lengths to zero scaled counts were correlated with gene parameters to study if various sequence and chromatin features impact the distance downstream of a TES that RT occurs (Extended Data Fig. 10a-b). Spearman correlation coefficients were calculated using cor.test(method = “Spearman”, exact = FALSE) in R (v4.0.4), between log10 transformed RT signal lengths and each of the following gene features: gene length (log10 bp), gene RPKM, distance to the nearest downstream gene (log10 bp, Supplementary Methods, section 7.4) on either strand or the same strand (ss), exon density (i.e. the number of exons), co-transcriptional splicing index^43^ (Supplementary Methods, section 7.9), GC-content of the sequences 1 kb upstream and 5 kb downstream of the TES (Supplementary Methods, section 7.6), the enrichment of the canonical poly-A signal (AATAAA) in the 5 kb downstream of the TES (PAS ratio, Supplementary Methods, section 7.6), and peaks of H3K36me3 and H3K79me2 signal downstream of the TES (TES+5kb), upstream of the TES (TES-1kb), and downstream of the TSS (TSS+1kb, Supplementary Methods, section 7.5). Each of these correlations was performed for all cell lines for which data was available. This analysis was also performed using RTsegs to compare results with an orthogonal method (Supplementary Methods, section 7.8).

### Comparisons of scaled RT signals between cell lines

To compare RT signal for the same gene across different cell lines, we obtained scaled counts as described above for 17377 bp regions (equal to the median length of all RT segments, Extended Data Fig. 8c). In addition to the filters applied to all RT signal regions, we removed RT genes that: 1.) had RTseg lengths less than the fixed region (17.3 kb), 2.) had scaled RT signals where one or more bins had a higher count value than the gene body maximum (i.e. scaled signal must be between 0 and 1), and 3.) that had a downstream gene on the same strand within 17.3 kb. These measures were taken, to avoid ambiguities in the downstream RT signal that are attributable to gene expression that is unrelated to the parent gene, however it is still possible that signal artifacts could exist in this subset (e.g. highly expressed enhancers or unannotated genes). This final subset of fixed-length RT signals was then aggregated per cell line and each RT signal was normalized to the median aggregated signal by subtracting the median value per cell line from each bin. The normalized signals from this analysis were then used to describe the variance between RT signals, where positive values indicate binned scaled counts higher than the median and vice versa.

To summarize RT signal variances, we calculated median normalized signals and net bin scores (nBSs) for the 17.3 kb RT regions divided into three 23 bin (∼5.8 kb) sectors. RT regions were divided into sectors due to the increased variability seen in the first sector (Extended Data Fig. 9b), due primarily to the higher signal found immediately downstream of the TES (Fig. 4e), as well as to provide additional resolution when assessing patterns of signal trajectories (Fig. 4g). The nBS was calculated by assessing the number of bins that were significantly above or below the median binned signal. Bin signals were significantly different from the median signal if their normalized count value was more than one standard deviation (stdev) from the median (i.e. 0) of the distribution of all normalized binned counts (Supplementary Fig. 16a-b). Each bin was assigned a score of +1 if the value is significantly higher than the median, −1 if it is significantly lower than the median, or 0 if it is similar to the median. These values were then summed per sector to calculate the nBS (Fig. 4f), with 23 being the highest possible signal and −23 the lowest, indicating that all bins are higher or lower than the median signal, respectively. To compare nBSs of genes with RT across all cell lines, hierarchical clustering was performed using pheatmap(clustering_method = “complete”, cluster_rows = T, cluster_cols = T). Examples of genes that have variable RT signals between cell lines were discovered by calculating the standard deviation of the nBS distribution (Fig. 4f) and the median normalized signal distribution (Supplementary Fig. 16c) in the first 5.8 kb sector and capturing those with the highest stdev in either or both metrics (Supplementary Fig. 16d-e).

## Acknowledgements

This body of work was supported by grants from the National Institute of Health’s National Human Genome Research Institute (NIH NHGRI, UM1HG009382). We thank the personnel at the University of Michigan Advanced Genomics Core for their professional technical assistance, as well as Andy Lin, Manhong Dai, and Fan Meng for the administration and maintenance of the University of Michigan Molecular and Behavioral Neuroscience Institute (MBNI) computing cluster.

We would like to thank each ENCODE consortium investigator, their laboratory, and all core services at their respective universities that contributed to the sequencing experiments and data processing that led to the generation of the ENCODE4 DPCL dataset. This includes Barbara Wold (California Institute of Technology, UM1HG009443), Ali Mortazavi (University of California, Irvine, UM1HG009443), Erez Lieberman Aiden (Baylor University, UM1HG009375), Charles Lee (The Jackson Laboratories, 3UM1HG009409-04S1), Michael Snyder (Stanford University, UM1HG009442), John Stamatoyannopoulos (University of Washington, UM1HG009444), and Haiyun Yu/John Lis (Cornell University, UM1HG009393).

Research reported in this publication was also supported by the National Cancer Institutes of Health under Award Number P30 CA046592 by the use of the following Cancer Center Shared Resources: Cancer Data Sciences (K.B.). Figure schematics were created with BioRender.com.

## Author Contributions

M.T.P. cultured and collected all DPCL samples, with support from H.B. and supervision from M.L., and distributed them to the ENCODE consortium laboratories that were involved in the DPCL effort. M.T.P. carried out all experiments and library preparations involved in the generation of the Bru-seq and BruUV-seq data used heavily in this study. B.M. ensured proper documentation and sharing of all Bru-seq data to the ENCODE portal, with support from I.V..

A.M. and I.V. conceived of and performed the majority of the analyses presented in this study, with direct support from M.A., K.B., B.M., and N.T.Y.. A.M. and I.V. wrote the manuscript with input from all authors.

## Extended Data

**Extended Data Fig. 1.**
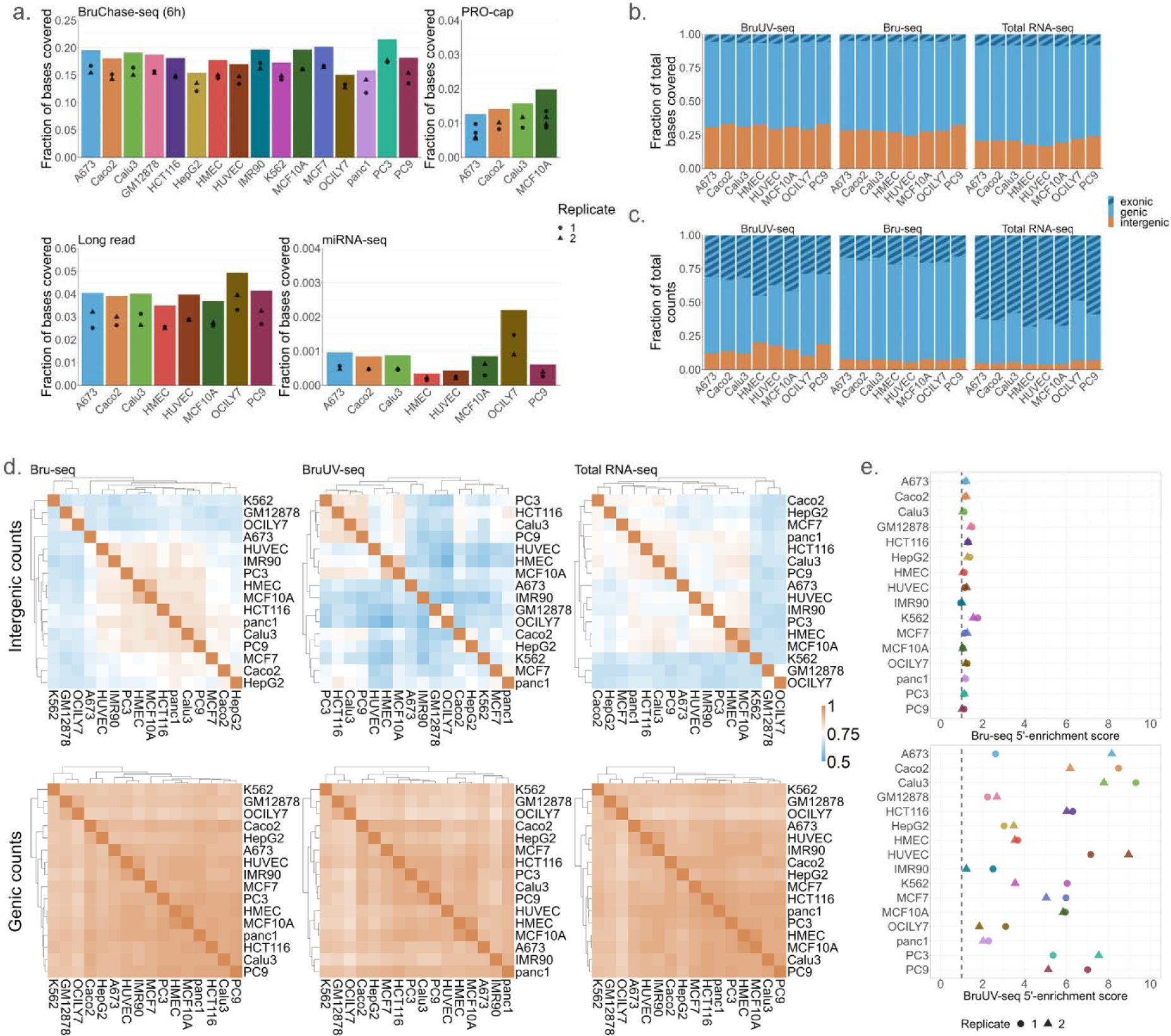
Genomic coverage distinguishes DPCL transcription assays. **a.** Fraction of bases covered per cell line for each DPCL transcription assay. Bars indicate the cumulative values for each cell line (N = 2), and dots indicate the individual replicate coverages. **b,c.** For select assays, the fraction of bases covered (**b**) and total uniquely mapping reads (**c**) per cell line (N = 2) was calculated for genic and intergenic compartments of the genome (based on GENCODE v29), with the striped bar indicating the proportion of genic coverage over exons. Eight cell lines for which replicate data was available in all assays are shown. **d.** Intergenic versus genic counts correlations (Pearson’s r) and hierarchical clustering of cell lines are shown for select assays. Color scale shows shift from moderate (0.5) to high (1) correlations between cell lines, highlighting increased variability in the intergenic space. **e.** Quantification of 5’-signal enrichment of Bru-seq and BruUV-seq in the 15 kb downstream of the TSS. BruUV-seq shows a higher 5’-enrichment and is also more variable between cell lines (Supplementary Note 2-3).

**Extended Data Fig. 2.**
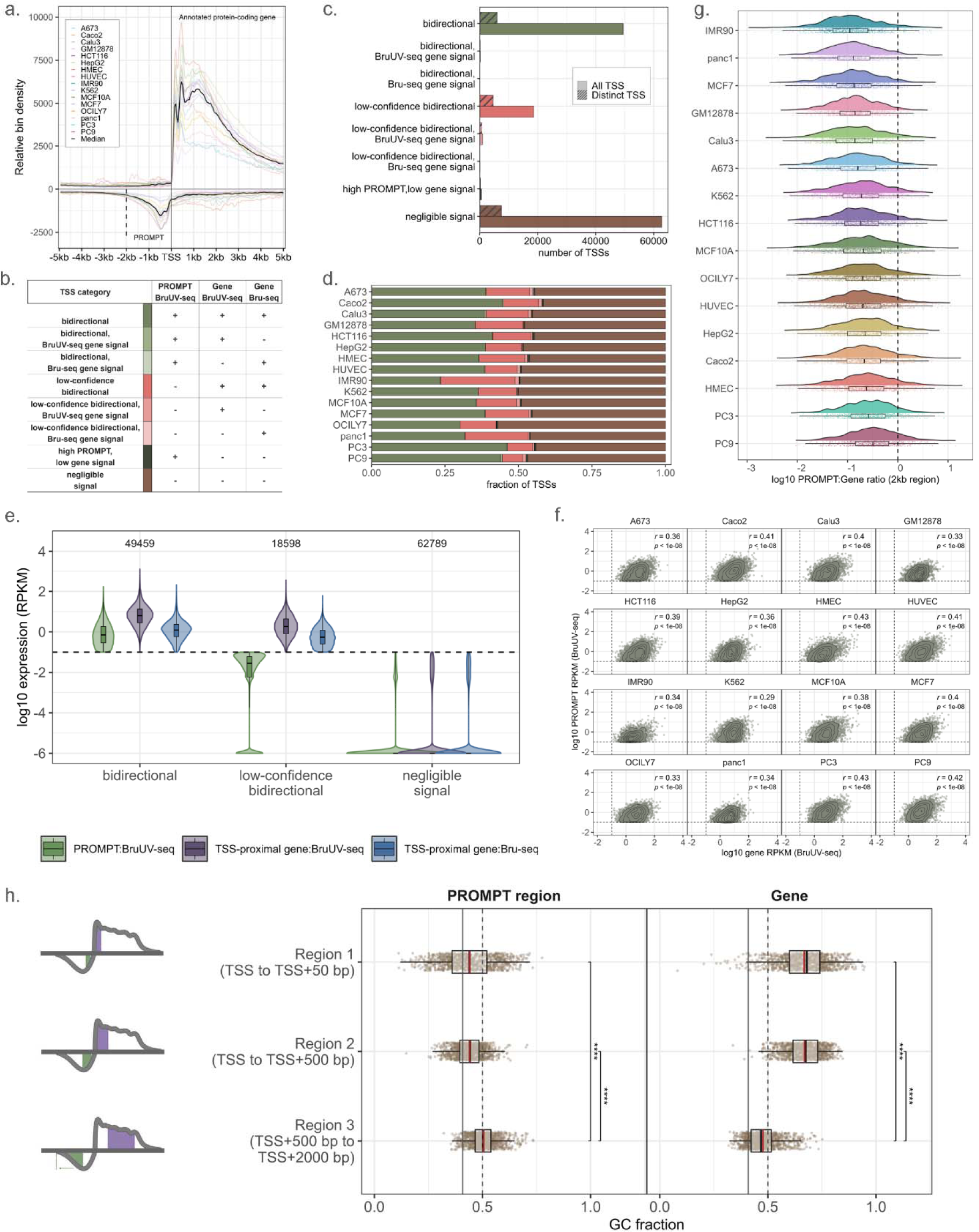
Characterization of TSS-proximal profiles. **a.** Metagene plot for 16 cell lines shows an enrichment of 5’-end signal flanking the TSS of annotated protein-coding genes (GENCODE v29 annotation, N_TSS_ = 5615, BruUV-seq). Each cell line’s signal profile is a combination of two replicates, with the black solid line denoting median binned signal for all cell lines. The black dotted line indicates the bounds of the 2 kb region used for describing PROMPT regions. **b.** All unambiguous TSS categories and an abridged set of parameters used for their classification. “+” or “-” indicates above or below the established parameter thresholds (see Methods: TSS category generation). **c.** Combined number of all TSSs (plain bars, N_total_ = 207024) or distinct TSSs (striped bars, N_total_ = 12939) for all cell lines per category. **d.** Distribution of TSS categories per cell line. **e.** Distribution of BruUV-seq and Bru-seq RPKMs for the PROMPT and TSS-proximal 2kb regions for the largest categories. Zero RPKM values are represented by a log10 value of −6. The total number of observations per category is indicated at the top of the plot. **f.** Scatterplots showing the relationship between PROMPT and TSS-proximal gene expression (2 kb regions, BruUV-seq RPKM) for all cell lines. Pearson’s correlations (r), associated p-values and 2D density contour plots are displayed. **g.** Distribution of PROMPT:Gene (P:G) expression ratios of 2kb regions flanking the TSS, with the cell lines ordered per increasing median ratios. The dotted line indicates a P:G ratio of 1. **h.** Distribution of GC-content (fraction) for three regions based on transcriptional signal over the PROMPT and gene (Region 1: TSS to TSS+50 bp, Region 2: TSS to TSS+500 bp, Region 3: TSS+500 bp to TSS+2000 bp) excluding the nucleosome-depleted region for the PROMPT regions. Within the boxplot: solid black line = median, solid red line=mean. For the entire plot: solid black line = genome-wide GC-content^42^ (0.408961), dotted black line = reference line (0.5). False discovery rate (FDR) adjusted p-values were determined from Wilcoxon signed-rank tests: ^ns^p_adj_ > 0.05, *p_adj_ < 0.05, **p_adj_ < 0.01, ***p_adj_ < 0.001, ****p_adj_ < 0.0001.

**Extended Data Fig. 3.**
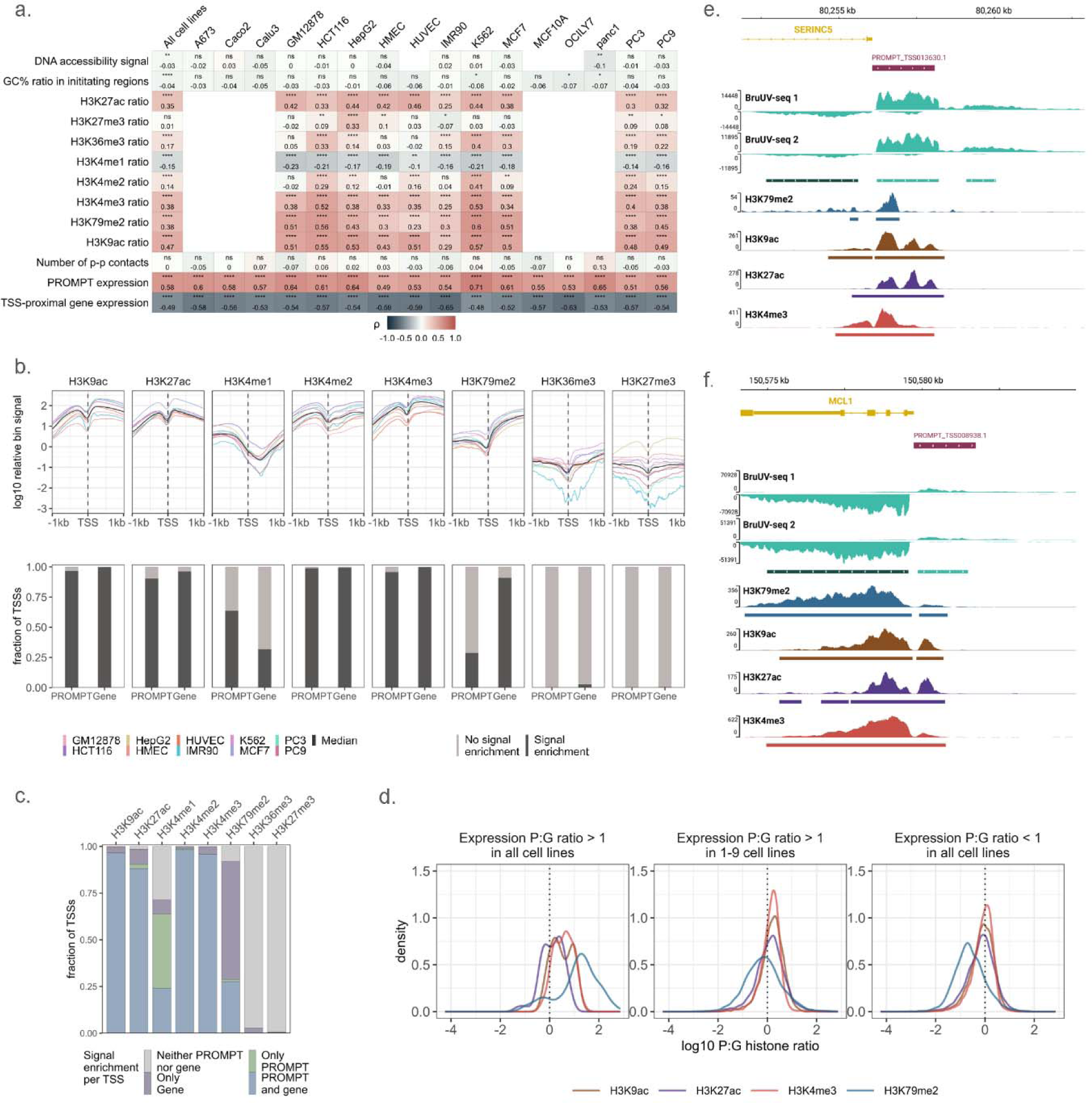
Factors associated with bidirectional TSS profiles in all cell lines. **a.** Heatmap of Spearman correlation coefficients (Iii) between various genomic and chromatin features and expression (RPKM) ratios over PROMPT and gene regions flanking the TSS. FDR adjusted p-values are indicated by ^ns^p_adj_ > 0.05, *p_adj_ < 0.05, **p_adj_ < 0.01, ***p_adj_ < 0.001, ****p_adj_ < 0.0001. **b.** Aggregated signal profile (top panel) and signal enrichment (determined by peak overlaps, bottom panel) over PROMPT and gene regions for selected histone post-translational modifications. **c.** Signal enrichment (peak overlap) status for PROMPT and gene regions per TSS. **d.** Density plots of the distribution of P:G histone ratios per P:G expression ratio profiles for selected histone modifications. **e-f.** Examples of bidirectional TSSs displaying differential expression (BruUV-seq, 2 biological replicates shown separately) and histone signal patterns (ChIP-seq, merged replicates for signal and peaks) over the gene an PROMPT regions in K562.

**Extended Data Fig. 4.**
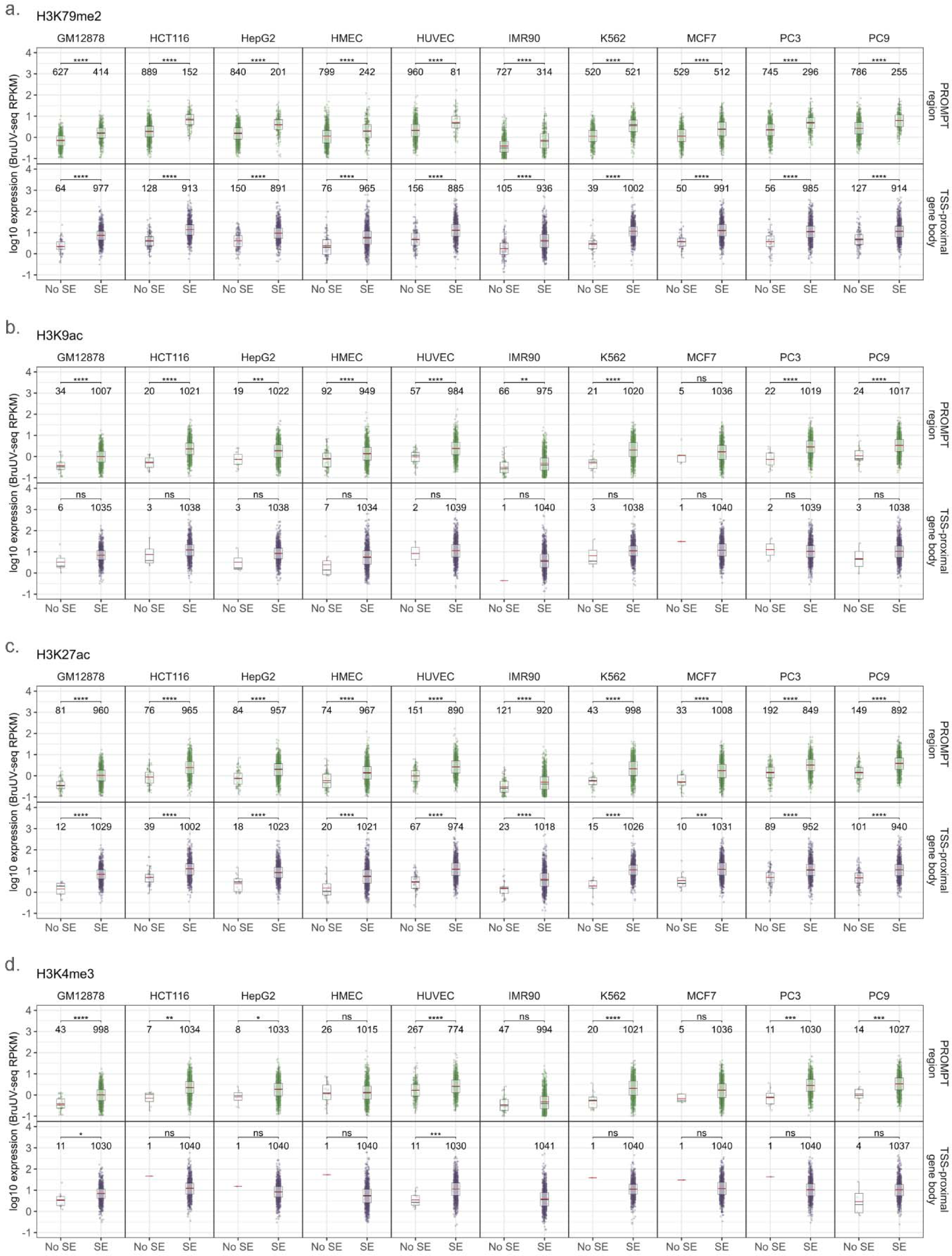
Signal enrichment of histone modifications at expressed PROMPTs and genes. **a-d.** Distribution of PROMPT or TSS-proximal gene expression (RPKM) based on histone signal enrichment status (determined by peak overlaps) for all the cell lines assayed (10/16). The number of observations is denoted per signal enrichment group. Wilcoxon signed-rank test FDR adjusted p-values are represented by ^ns^p_adj_ > 0.05, *p_adj_ < 0.05, **p_adj_ < 0.01, ***p_adj_ < 0.001, ****p_adj_ < 0.0001. Horizontal lines in the boxplot represent the following: solid black line = median, solid red line = mean.

**Extended Data Fig. 5.**
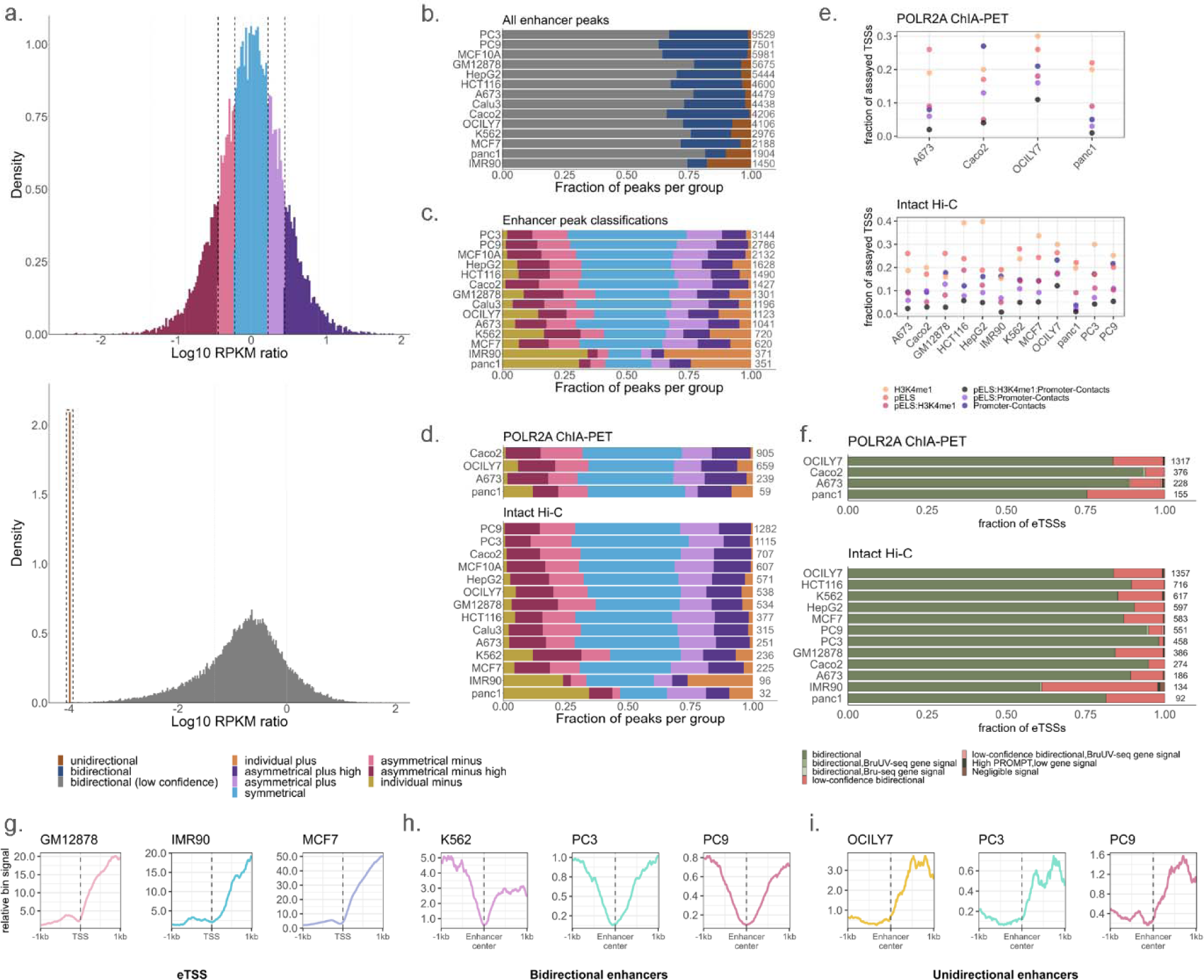
Enhancer and eTSS class distributions are generally similar between cell lines. **a.** Log10 BruUV RPKM ratio distributions for bidirectional peak pairs (top) and individual peaks and their divergent region (bottom). These distributions were used to classify bidirectional peaks into symmetrical and asymmetrical groups, where dotted lines indicate demarcations between groups at 0.5 and 1 standard deviation from the mean (see Methods). Unidirectional enhancers were extracted from the individual peaks (RPKM ratio = 0; Log10 ratio = −4) and the rest were considered low-confidence bidirectional. **b,c.** Peak classification distributions per cell line, with the total number of enhancer peaks indicated alongside each bar. For all enhancer peaks, the proportions of bidirectional versus unidirectional enhancers are shown (**b**), along with the classification breakdown for all enhancers (excluding low-confidence bidirectional) is shown (**c**). **d.** Peak classifications per cell line for enhancers interacting with expressed genes via POLR2A ChIA-PET or intact Hi-C loops. **e.** Fraction of assayed TSSs (N = 12939) displaying features associated with enhancers. Promoter-promoter contacts were obtained from POLR2A ChIA-PET or intact Hi-C datasets. **f.** Distribution of TSS categories for enhancer-like TSSs (eTSSs) per cell line, with the total number of eTSSs displayed adjacent to the bars. **g-i.** H3K79me2 aggregate signal profiles for selected cell lines for eTSSs (**g**), bidirectional enhancers (**h**), and unidirectional enhancers (**i**).

**Extended Data Fig. 6.**
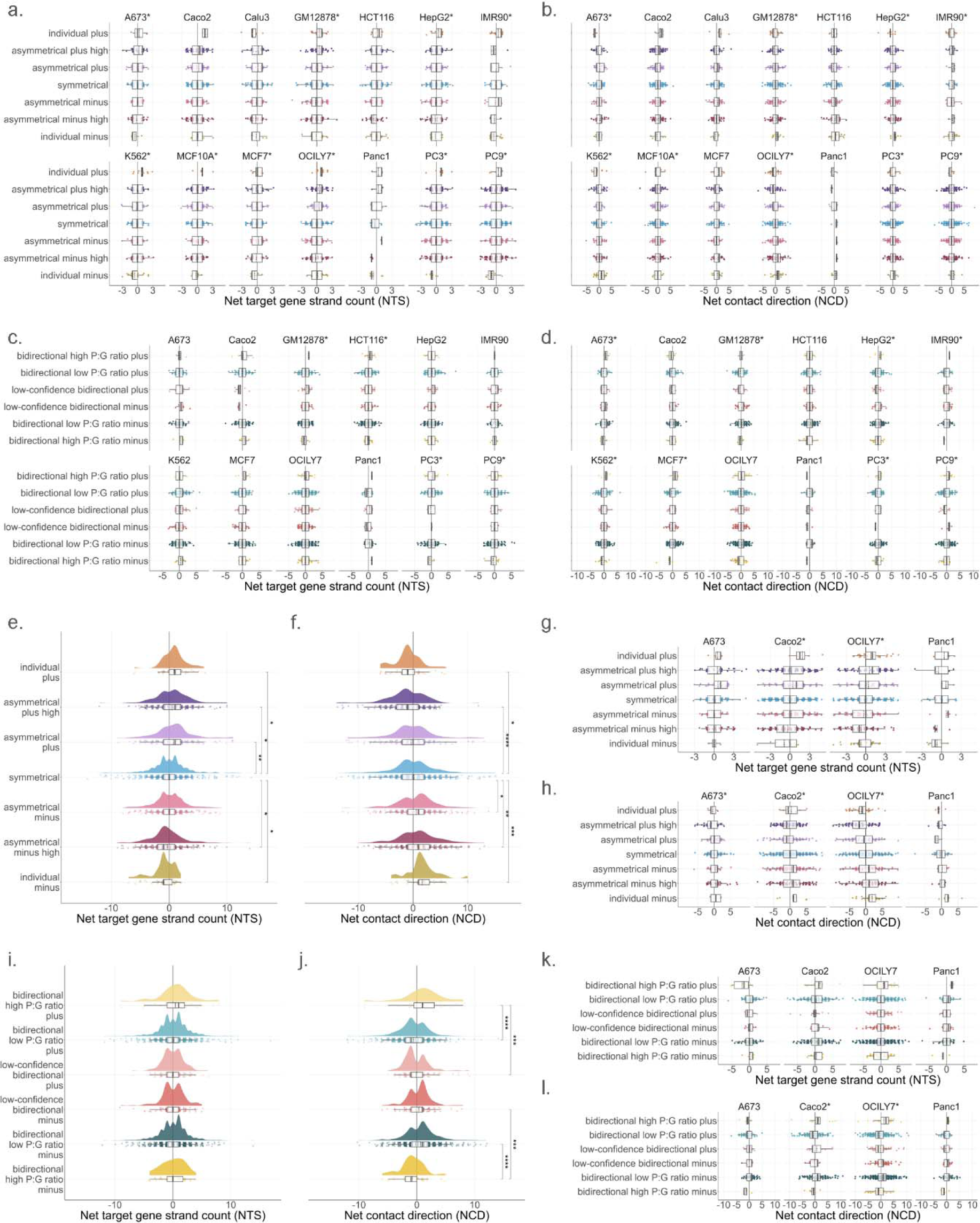
nTS and nCD patterns in individual cell lines. **a-d.** Cell line distributions of nTS (**a,c**) and nCD (**b,c**) for enhancers (**a-b**) and eTSSs (**c-d**) interacting distally with expressed genes via intact Hi-C loops. e-l. Cumulative (**e-f,i-j**) and cell line specific (**g-h,k-l**) distributions of nTS and nCD for enhancers and eTSSs engaging in long-range contacts with expressed genes in POLR2A ChIA-PET data. Asterisks beside cell line names indicate those that generally follow nTS or nCD trends observed cumulatively for all elements in Hi-C (Fig. 3e-f) and ChIA-PET (**e-f,i-j**). FDR adjusted p-values were determined for aggregated ChIA-PET data (**e**) from Wilcoxon signed-rank tests (*p_adj_ < 0.05, **p_adj_ < 0.01, ***p_adj_ < 0.001, ****p_adj_ < 0.0001).

**Extended Data Fig. 7.**
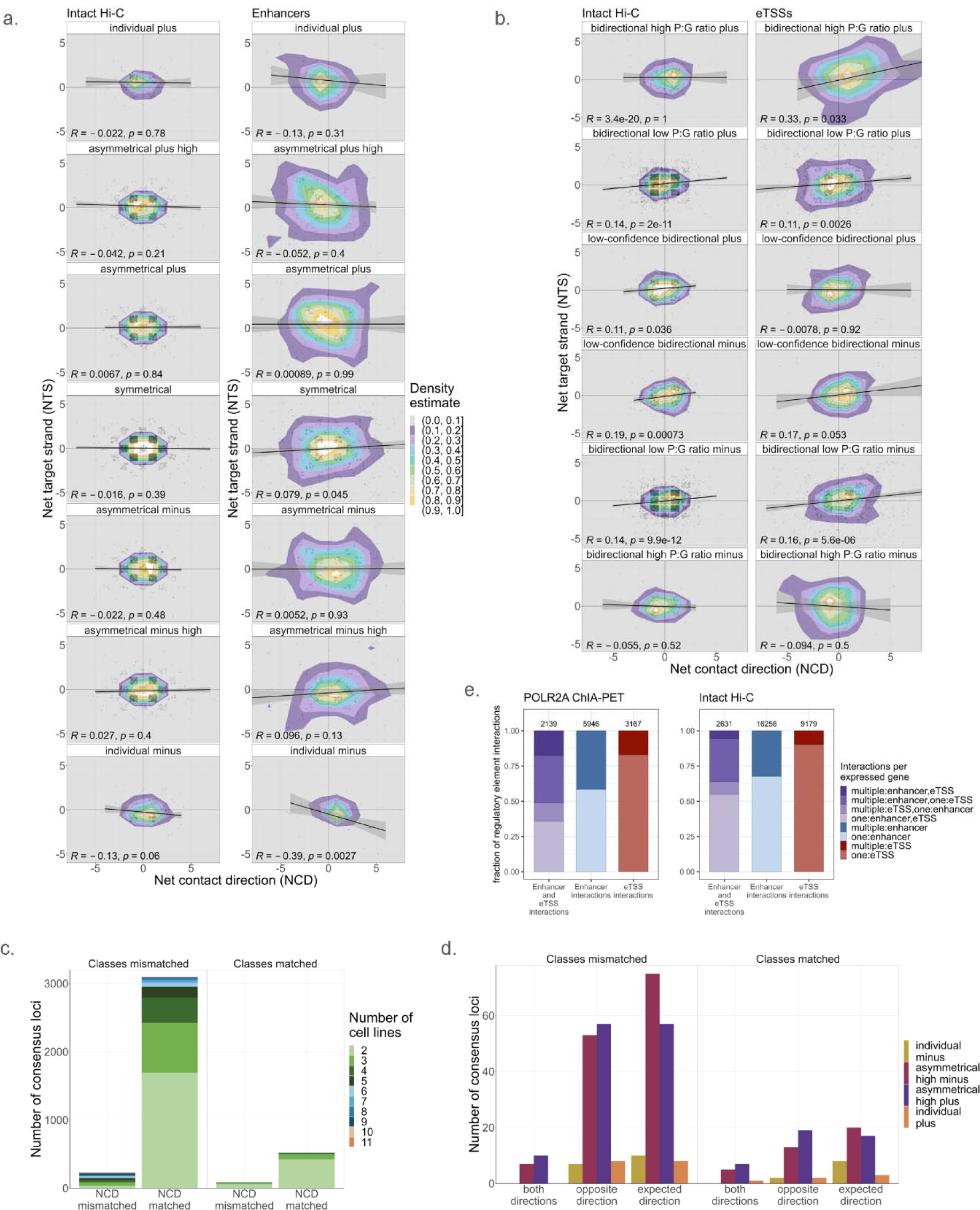
nTS and nCD coincide but do not explain cell line specific signal patterns. **a,b.** Scaled kernel density estimations for nTS and nCD distributions of grouped enhancers (**a**) and eTSSs (**b**). Scaling is performed within groups to account for sample size imbalances. **c.** Quantification of active enhancers that are found in at least 2 cell lines (at consensus loci, see Methods), and are interacting with expressed genes via intact Hi-C loops, with nCD scores that are in the same (matched) or opposite (mismatched) net directions. Consensus loci are distinguished by the classifications of the enhancers in each cell line represented, where either all classes match or at least one cell line has a different signal classification (mismatched). This shows that at most consensus loci a particular enhancer tends to make the same net contacts in all cell lines, even if the signal pattern changes between cell lines. **d.** Quantification of enhancers found in at least 2 cell lines, where multiple cell lines have enhancers that are highly asymmetrical or unidirectional, with nCD scores that follow the expected contact direction pattern (Fig. 3d), show the opposite pattern, or show different patterns between cell lines. Consensus loci are again distinguished by matched or mismatched enhancer classifications between cell lines. Since enhancers at these loci are nearly equally likely to match the expected relationship between eRNA signal and nCD or not, with very few instances of mixed patterns, irrespective of any differences in their signal patterns, we conclude that nCD does not dictate changes in enhancer expression. **e.** Fraction of enhancers or eTSSs engaging in single or multiple interactions with target genes via Hi-C or ChIA-PET loops. Instances of target gene interacting with one or both elements are observed, however, individual interactions are typically more common than multiplex interactions.

**Extended Data Fig. 8.**
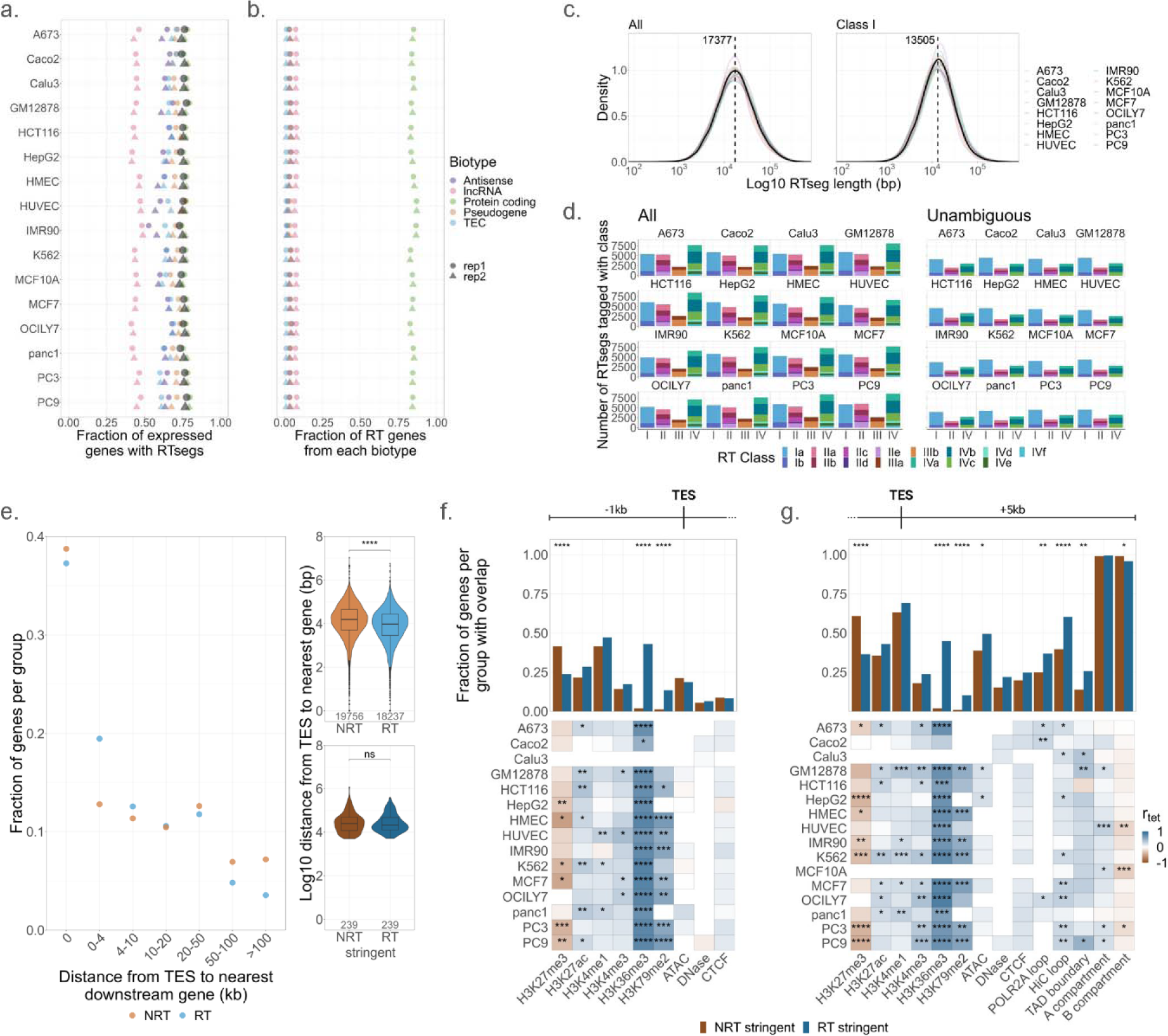
Overview of nascent RTseg and gene features. **a.** Fraction of expressed genes per cell line with an assigned RTseg. For both replicates, the fraction of all expressed genes (black) and the fraction of expressed genes broken down by biotype is shown. **b.** Biotype breakdown of all RT genes identified for each cell line per replicate. **c.** Log10 distribution of lengths for all RTsegs (left) and class I RTsegs (right). Cumulative distributions are shown (black), as in Fig. 4b, along with cell line specific distributions. Dashed line indicates overall median RTseg length. **d.** Classification breakdown for all (left) and unambiguous (right, see Supplementary Methods, section 7.1) RTsegs. Classification descriptions are in Supplementary Table 4. Importantly, RTsegs can be assigned multiple classifications, describing their overlap with multiple genes, thus counts in this plot do not sum to the total number of RTsegs per cell line. **e.** Distances to the nearest downstream gene on either strand for RT or NRT genes. The overall log10 distribution of distances between genes (right) is shown for all RT/NRT genes (top) and stringent RT/NRT genes (bottom). Additionally, the fractions of RT/NRT genes that have a downstream gene within various distance ranges are shown. **f,g.** Overlap of stringent RT/NRT genes with peaks of histone modification or CTCF signal, chromatin accessibility, loop anchors (ChIA-PET and intact Hi-C), chromatin domain (TAD) boundaries, and A/B compartments in the 1 kb upstream and 5 kb downstream of the TES. The fraction of RT/NRT genes that had each of these features in any cell line is shown (top), along with the tetrachoric correlation (r_tet_) between RT/NRT genes and the presence or absence of each feature per cell line (bottom). Fisher’s exact p-values are designated wher significant: *p_adj_ < 0.05, **p_adj_ < 0.01, ***p_adj_ < 0.001, ****p_adj_ < 0.0001. For **e-g** similar patterns to known RT transcripts^35–37^ were found.

**Extended Data Fig. 9.**
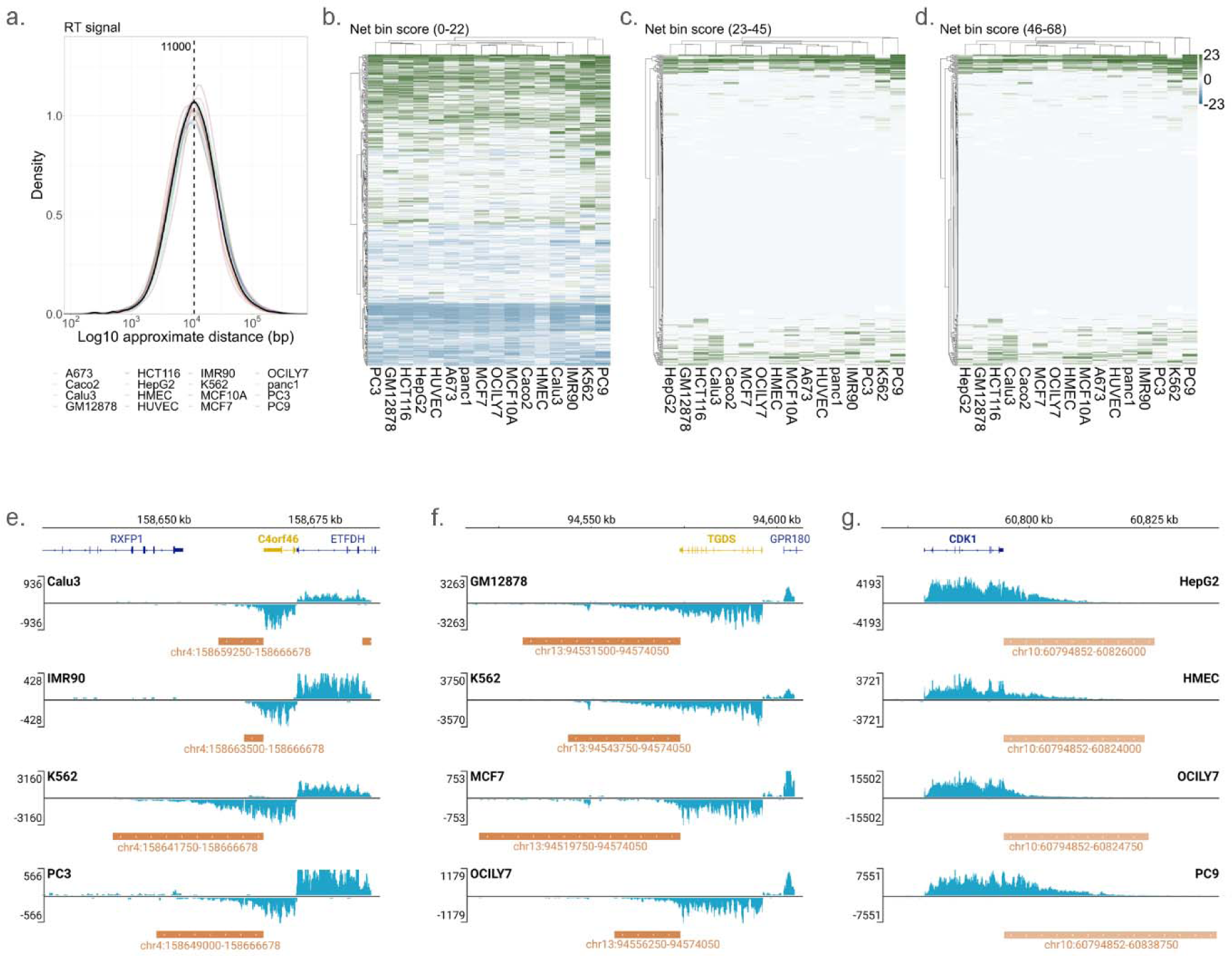
Overview of RT signal patterns. **a.** Log10 approximate distance that each RT signal travels before reaching 0 signal. The cumulative distribution is shown (black) along with cell line specific distributions. Dashed line indicates overall median RT signal length. **b-d.** Net bin scores for each 23 bin (∼5.8 kb) sector of the RT signal regions (17.3 kb) found in all 16 cell lines. Hierarchical clustering was performed for cell lines and genes, and depicts the increased variability observed in the first ∼5 kb downstream of the TES. **e-g.** Bru-seq signal and RTsegs for the three genes highlighted in Fig. 4h-j. Merged replicate Bru-seq signal is shown for the four featured cell lines, along with the merged replicate RTseg labeled with its RT_ID.

**Extended Data Fig. 10.**
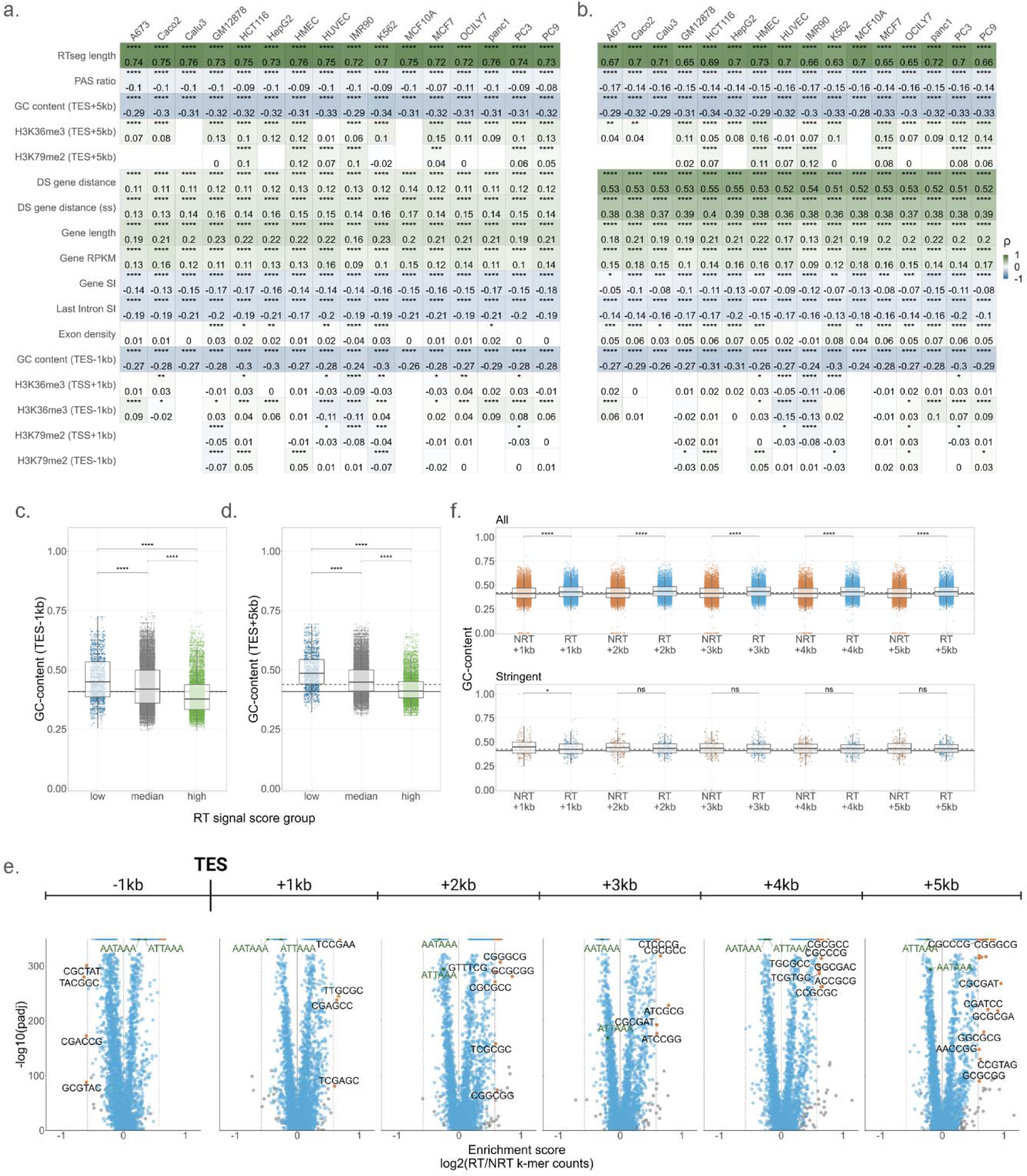
GC-content and G-rich sequences correlate with RT signal length and magnitude. **a,b.** Correlations between RT signal length (approximate distance) and gene sequence or chromatin features for all RT signal regions (**a**) and RT regions that end > 5 kb away from annotated genes (**b**, RTseg class Ia). Spearman’s rank correlation coefficients (ρ) and FDR adjusted p-values are displayed. **c,d.** GC-content for the 1 kb upstream (**c**) an the 5 kb downstream (**d**) of the TES, for RT signals grouped by their first sector net bin scores as in Fig. 4f. Horizontal lines represent the genome-wide GC-content^42^ (solid), and the median GC-content of all 1 kb or 5 kb regions (dashed). FDR adjusted p-values were determined from Wilcoxon tests. **e.** Hexamer enrichment scores for all 1 kb sequences in the 1 kb upstream and 5 kb downstream of the TES for all genes. Gray dots indicate hexamers with fewer than 100 counts per group and labeled hexamers (orange dots) have enrichment scores > 0.56. Labeled in green are the two canonical poly-A signals. **f.** GC-content of 1 kb sequences described in e for all (top) or stringent (bottom) RT/NRT genes. Horizontal lines depict genome and region-specific GC-content as described for **c-d**. FDR adjusted p-values were determined from Wilcoxon tests. Adjusted p-values (**a-d,f**) are denoted as: *p_adj_ < 0.05, **p_adj_ < 0.01, ***p_adj_ < 0.001, ****p_adj_ < 0.0001.

## References

1. Girbig, M., Misiaszek, A.D. & Muller, C.W. Structural insights into nuclear transcription by eukaryotic DNA-dependent RNA polymerases. Nat Rev Mol Cell Biol 23, 603–622 (2022).

2. Nojima, T. & Proudfoot, N.J. Mechanisms of lncRNA biogenesis as revealed by nascent transcriptomics. Nat Rev Mol Cell Biol 23, 389–406 (2022).

3. Jonkers, I. & Lis, J.T. Getting up to speed with transcription elongation by RNA polymerase II. Nat Rev Mol Cell Biol 16, 167–77 (2015).

4. Chen, F.X., Smith, E.R. & Shilatifard, A. Born to run: control of transcription elongation by RNA polymerase II. Nat Rev Mol Cell Biol 19, 464–478 (2018).

5. Grzechnik, P., Tan-Wong, S.M. & Proudfoot, N.J. Terminate and make a loop: regulation of transcriptional directionality. Trends Biochem Sci 39, 319–27 (2014).

6. Agostini, F., Zagalak, J., Attig, J., Ule, J. & Luscombe, N.M. Intergenic RNA mainly derives from nascent transcripts of known genes. Genome Biol 22, 136 (2021).

7. van Bakel, H., Nislow, C., Blencowe, B.J. & Hughes, T.R. Most “dark matter” transcripts are associated with known genes. PLoS Biol 8, e1000371 (2010).

8. Porrua, O. & Libri, D. Transcription termination and the control of the transcriptome: why, where and how to stop. Nat Rev Mol Cell Biol 16, 190–202 (2015).

9. Kapranov, P. et al. RNA maps reveal new RNA classes and a possible function for pervasive transcription. Science 316, 1484–8 (2007).

10. Andersson, R. & Sandelin, A. Determinants of enhancer and promoter activities of regulatory elements. Nat Rev Genet 21, 71–87 (2020).

11. van Arensbergen, J., van Steensel, B. & Bussemaker, H.J. In search of the determinants of enhancer-promoter interaction specificity. Trends Cell Biol 24, 695–702 (2014).

12. Vilborg, A. & Steitz, J.A. Readthrough transcription: How are DoGs made and what do they do? RNA Biol 14, 632–636 (2017).

13. Djebali, S. et al. Landscape of transcription in human cells. Nature 489, 101–8 (2012).

14. Reese, F., et al. The ENCODE4 long-read RNA-seq collection reveals distinct classes of transcript structure diversity. bioRxiv (2023).

15. Blasius, M., Wagner, S.A., Choudhary, C., Bartek, J. & Jackson, S.P. A quantitative 14-3-3 interaction screen connects the nuclear exosome targeting complex to the DNA damage response. Genes Dev 28, 1977–82 (2014).

16. Lubas, M. et al. The human nuclear exosome targeting complex is loaded onto newly synthesized RNA to direct early ribonucleolysis. Cell Rep 10, 178–92 (2015).

17. Preker, P. et al. RNA exosome depletion reveals transcription upstream of active human promoters. Science 322, 1851–4 (2008).

18. Andersson, R. et al. Human Gene Promoters Are Intrinsically Bidirectional. Mol Cell 60, 346–7 (2015).

19. Chen, Y. et al. Principles for RNA metabolism and alternative transcription initiation within closely spaced promoters. Nat Genet 48, 984–94 (2016).

20. Core, L.J., Waterfall, J.J. & Lis, J.T. Nascent RNA sequencing reveals widespread pausing and divergent initiation at human promoters. Science 322, 1845–8 (2008).

21. Duttke, S.H.C. et al. Human promoters are intrinsically directional. Mol Cell 57, 674–684 (2015).

22. Almada, A.E., Wu, X., Kriz, A.J., Burge, C.B. & Sharp, P.A. Promoter directionality is controlled by U1 snRNP and polyadenylation signals. Nature 499, 360–3 (2013).

23. Ntini, E. et al. Polyadenylation site-induced decay of upstream transcripts enforces promoter directionality. Nat Struct Mol Biol 20, 923–8 (2013).

24. Scruggs, B.S. et al. Bidirectional Transcription Arises from Two Distinct Hubs of Transcription Factor Binding and Active Chromatin. Mol Cell 58, 1101–12 (2015).

25. Core, L.J. et al. Analysis of nascent RNA identifies a unified architecture of initiation regions at mammalian promoters and enhancers. Nat Genet 46, 1311–20 (2014).

26. Seila, A.C. et al. Divergent transcription from active promoters. Science 322, 1849–51 (2008).

27. Jin, Y., Eser, U., Struhl, K. & Churchman, L.S. The Ground State and Evolution of Promoter Region Directionality. Cell 170, 889–898 e10 (2017).

28. Lidschreiber, K. et al. Transcriptionally active enhancers in human cancer cells. Mol Syst Biol 17, e9873 (2021).

29. Lewis, M.W., Li, S. & Franco, H.L. Transcriptional control by enhancers and enhancer RNAs. Transcription 10, 171–186 (2019).

30. Andersson, R. et al. An atlas of active enhancers across human cell types and tissues. Nature 507, 455–461 (2014).

31. Mikhaylichenko, O. et al. The degree of enhancer or promoter activity is reflected by the levels and directionality of eRNA transcription. Genes Dev 32, 42–57 (2018).

32. Zhang, Y. et al. Model-based analysis of ChIP-Seq (MACS). Genome Biol 9, R137 (2008).

33. Consortium, E.P. et al. Expanded encyclopaedias of DNA elements in the human and mouse genomes. Nature 583, 699–710 (2020).

34. Koch, F. et al. Transcription initiation platforms and GTF recruitment at tissue-specific enhancers and promoters. Nat Struct Mol Biol 18, 956–63 (2011).

35. Caldas, P. et al. Transcription readthrough is prevalent in healthy human tissues and associated with inherent genomic features. Commun Biol 7, 100 (2024).

36. Vilborg, A. et al. Comparative analysis reveals genomic features of stress-induced transcriptional readthrough. Proc Natl Acad Sci U S A 114, E8362–E8371 (2017).

37. Rosa-Mercado, N.A. & Steitz, J.A. Who let the DoGs out? - biogenesis of stress-induced readthrough transcripts. Trends Biochem Sci 47, 206–217 (2022).

38. Hennig, T. et al. HSV-1-induced disruption of transcription termination resembles a cellular stress response but selectively increases chromatin accessibility downstream of genes. PLoS Pathog 14, e1006954 (2018).

39. Schwalb, B. et al. TT-seq maps the human transient transcriptome. Science 352, 1225–8 (2016).

40. Mo, W. et al. Landscape of transcription termination in Arabidopsis revealed by single-molecule nascent RNA sequencing. Genome Biol 22, 322 (2021).

41. Magnuson, B. et al. CDK12 regulates co-transcriptional splicing and RNA turnover in human cells. iScience 25, 105030 (2022).

42. Piovesan, A. et al. On the length, weight and GC content of the human genome. BMC Res Notes 12, 106 (2019).

43. Bedi, K. et al. Co-transcriptional splicing efficiencies differ within genes and between cell types. RNA 27, 829–40 (2021).

44. Ibrahim, M.M. et al. Determinants of promoter and enhancer transcription directionality in metazoans. Nat Commun 9, 4472 (2018).

45. Bornelov, S., Komorowski, J. & Wadelius, C. Different distribution of histone modifications in genes with unidirectional and bidirectional transcription and a role of CTCF and cohesin in directing transcription. BMC Genomics 16, 300 (2015).

46. Godfrey, L. et al. DOT1L inhibition reveals a distinct subset of enhancers dependent on H3K79 methylation. Nat Commun 10, 2803 (2019).

47. Lim, B. & Levine, M.S. Enhancer-promoter communication: hubs or loops? Curr Opin Genet Dev 67, 5–9 (2021).

48. Di Giammartino, D.C., Polyzos, A. & Apostolou, E. Transcription factors: building hubs in the 3D space. Cell Cycle 19, 2395–2410 (2020).

49. Barshad, G. et al. RNA polymerase II dynamics shape enhancer-promoter interactions. Nat Genet 55, 1370–1380 (2023).

50. Gu, B. et al. Transcription-coupled changes in nuclear mobility of mammalian cis-regulatory elements. Science 359, 1050–1055 (2018).

51. Benedetti, F., Dorier, J. & Stasiak, A. Effects of supercoiling on enhancer-promoter contacts. Nucleic Acids Res 42, 10425–32 (2014).

52. Ljungman, M. & Hanawalt, P.C. Presence of negative torsional tension in the promoter region of the transcriptionally poised dihydrofolate reductase gene in vivo. Nucleic Acids Res 23, 1782–9 (1995).

53. Janissen, R., Barth, R., Polinder, M., van der Torre, J. & Dekker, C. Single-molecule visualization of twin-supercoiled domains generated during transcription. Nucleic Acids Res 52, 1677–1687 (2024).

54. Liang, L. et al. Complementary Alu sequences mediate enhancer-promoter selectivity. Nature 619, 868–875 (2023).

55. Blumberg, A. et al. Characterizing RNA stability genome-wide through combined analysis of PRO-seq and RNA-seq data. BMC Biol 19, 30 (2021).

56. Zheng, M. et al. Multiplex chromatin interactions with single-molecule precision. Nature 566, 558–562 (2019).

57. Kouno, T. et al. C1 CAGE detects transcription start sites and enhancer activity at single-cell resolution. Nat Commun 10, 360 (2019).

58. Cai, Z. et al. RIC-seq for global in situ profiling of RNA-RNA spatial interactions. Nature 582, 432–437 (2020).

59. Skourti-Stathaki, K., Kamieniarz-Gdula, K. & Proudfoot, N.J. R-loops induce repressive chromatin marks over mammalian gene terminators. Nature 516, 436–9 (2014).

60. Geisberg, J.V. et al. Nucleotide-level linkage of transcriptional elongation and polyadenylation. Elife 11(2022).

61. Ginno, P.A., Lim, Y.W., Lott, P.L., Korf, I. & Chedin, F. GC skew at the 5’ and 3’ ends of human genes links R-loop formation to epigenetic regulation and transcription termination. Genome Res 23, 1590–600 (2013).

62. Shah, N. et al. Tyrosine-1 of RNA Polymerase II CTD Controls Global Termination of Gene Transcription in Mammals. Mol Cell 69, 48–61 e6 (2018).

63. Eaton, J.D. et al. Xrn2 accelerates termination by RNA polymerase II, which is underpinned by CPSF73 activity. Genes Dev 32, 127–139 (2018).

64. Arnold, M., Bressin, A., Jasnovidova, O., Meierhofer, D. & Mayer, A. A BRD4-mediated elongation control point primes transcribing RNA polymerase II for 3’-processing and termination. Mol Cell 81, 3589–3603 e13 (2021).

65. Jaeger, M.G. et al. Selective Mediator dependence of cell-type-specifying transcription. Nat Genet 52, 719–727 (2020).

66. Dasilva, L.F. et al. Integrator enforces the fidelity of transcriptional termination at protein-coding genes. Sci Adv 7, eabe3393 (2021).

67. Liu, X. et al. The PAF1 complex promotes 3’ processing of pervasive transcripts. Cell Rep 38, 110519 (2022).

## Method References

68. Kagda, M.S. et al. Data navigation on the ENCODE portal. ArXiv abs/2305.00006(2023).

69. Luo, Y. et al. New developments on the Encyclopedia of DNA Elements (ENCODE) data portal. Nucleic Acids Res 48, D882–D889 (2020).

70. Consortium, E.P. An integrated encyclopedia of DNA elements in the human genome. Nature 489, 57–74 (2012).

71. Hitz, B.C., et al. The ENCODE Uniform Analysis Pipelines. bioRxiv (2023).

72. Frankish, A. et al. GENCODE reference annotation for the human and mouse genomes. Nucleic Acids Res 47, D766–D773 (2019).

73. Ramirez, F. et al. deepTools2: a next generation web server for deep-sequencing data analysis. Nucleic Acids Res 44, W160–5 (2016).

74. Quinlan, A.R. & Hall, I.M. BEDTools: a flexible suite of utilities for comparing genomic features. Bioinformatics 26, 841–2 (2010).

75. Liao, Y., Smyth, G.K. & Shi, W. featureCounts: an efficient general purpose program for assigning sequence reads to genomic features. Bioinformatics 30, 923–30 (2014).

76. Dao, L.T.M. et al. Genome-wide characterization of mammalian promoters with distal enhancer functions. Nat Genet 49, 1073–1081 (2017).

77. Santiago-Algarra, D. et al. Epromoters function as a hub to recruit key transcription factors required for the inflammatory response. Nat Commun 12, 6660 (2021).

78. Tarasov, A., Vilella, A.J., Cuppen, E., Nijman, I.J. & Prins, P. Sambamba: fast processing of NGS alignment formats. Bioinformatics 31, 2032–4 (2015).

79. Bedi, K., Paulsen, M.T., Wilson, T.E. & Ljungman, M. Characterization of novel primary miRNA transcription units in human cells using Bru-seq nascent RNA sequencing. NAR Genom Bioinform 2, lqz014 (2020).

80. Paulsen, M.T. et al. Use of Bru-Seq and BruChase-Seq for genome-wide assessment of the synthesis and stability of RNA. Methods 67, 45–54 (2014).

81. Paulsen, M.T. et al. Coordinated regulation of synthesis and stability of RNA during the acute TNF-induced proinflammatory response. Proc Natl Acad Sci U S A 110, 2240–5 (2013).

