## Supplementary Material for "Characterizing nascent transcription patterns of PROMPTs, eRNAs, and readthrough transcripts in the ENCODE4 deeply profiled cell lines"

### Supplementary Materials Outline

*Supplementary Notes*

1. Factors influencing genome coverage calculations
2. Using Bru-seq and BruUV-seq to study intergenic co-products of transcription
3. Describing cell-to-cell variability in BruUV-seq data
4. Limitations of determining full-length PROMPT, eRNA, and RT transcription units

*Supplementary Methods*

1. Cell Collection and Experimental Protocols
    - 1.1. ATAC-seq and snATAC-seq
    - 1.2. Bru-seq suite of techniques
        - 1.2.1. Bru-seq
        - 1.2.2. BruChase-seq
        - 1.2.3. BruUV-seq
        - 1.2.4. Stranded library preparation (all Bru-seq assays)
    - 1.3. ChIA-PET
    - 1.4. DNase-seq
    - 1.5. Intact Hi-C
    - 1.6. PRO-cap
    - 1.7. RNA-seq
        - 1.7.1. Long-read RNA-seq and miRNA-seq
        - 1.7.2. Total RNA-seq
2. Sequencing Protocols
    - 2.1. ATAC-seq and snATAC-seq
    - 2.2. Bru-seq, BruChase-seq, BruUV-seq
    - 2.3. ChIA-PET
    - 2.4. DNase-seq
    - 2.5. Intact Hi-C
    - 2.6. PRO-cap
    - 2.7. Long-read RNA-seq
    - 2.8. miRNA-seq
    - 2.9. Total RNA-seq
3. Genome Coverage
    - 3.1. Fraction of genome covered by PROMPT, eRNA, and RT regions
    - 3.2. Overlap of PROMPT, eRNA, and RT region with GENCODE v44 annotation
4. Bru-seq and BruUV-seq QC
    - 4.1. Quality control metrics
    - 4.2. 5’ enrichment score
    - 4.3. Principal component analysis (PCA) of regions of interest
5. PROMPTs
    - 5.1. Metagene plots
    - 5.2. TSS generation and signal characterization

5.3. QC analysis of PROMPT regions
5.4. PROMPT:Gene (P:G) ratio correlations (expanded)
5.4.1. PROMPT and genic signal
5.4.2. DNA accessibility signal
5.4.3. GC% ratio in initiating regions
5.4.4. Number of promoter-promoter (p-p) contacts
5.4.5. Histone post-translational modifications
5.5. eTSS considerations
6. eRNAs
6.1. Enhancer/eRNA peak QC
6.2. BruUV-peak validation and feature summary
6.3. nTS/nCD validation with predicted ENCODE4 E2G contacts
6.4. eRNA consensus loci analysis (nCD)
6.5. nTS/nCD kernel density estimation
7. Readthrough
7.1. Readthrough segment (RTseg) classification
7.2. Gene biotype group definitions
7.3. Implications of merging RTsegs for replicate samples
7.4. Downstream gene proximity for RT/NRT genes
7.5. Assessing the chromatin landscape of RT/NRT genes
7.6. Sequence analysis of RT/NRT genes
7.7. Readthrough signal region distinctions
7.8. RT signal versus RTseg lengths
7.9. Co-transcriptional splicing index calculation

*Supplementary Data Files*

1. ENCODE file accessions and metadata
2. DPCL genome coverage data
3. PROMPT regions and data
4. eRNA regions and data
5. RTseg regions and data

*Supplementary Tables*

1. Cell line information
2. Software versions
3. Overview of TSS categories
4. Overview of RTseg classifications

*Supplementary Figures*

1. Genome coverage of DPCL assays and RNA species
2. Metagene plots of Bru-seq and BruUV-seq data for 16 cell lines
3. Comparisons of Bru-seq and BruUV-seq data

4. Principal Component Analysis for regions of interest
5. Signal profiles in 2 kb genome bins
6. Signal profiles of TSS-proximal regions and association of PROMPT regions with supplemental and orthogonal data types
7. Agreement between histone signal and peaks
8. Summary of expression, sequence and chromatin attributes for gene and PROMPT regions
9. Correlations of chromatin features with expression and properties of enhancer-like gene TSSs (eTSS)
10. Overview of BruUV-peak features
11. nTS and nCD validation with E2G prediction data
12. Higher resolution kernel density estimations
13. RT and NRT gene distinctions and RTseg lengths
14. RT and NRT gene features
15. Chromatin features of RT and NRT genes
16. RT signal scaled counts and net bin score distributions

*Supplemental References*

1. ENCODE project portal links
2. References

### Supplementary Notes

#### Note 1: Factors influencing genome coverage calculations

Previous reports from the ENCODE consortium using cumulative data from 15 human cell lines stated that 74.7% of the unstranded genome is covered by primary transcripts[13]. These previous estimates relied on total RNA-seq contigs to infer RNA synthesis from a primarily mature RNA pool as a representation of RNA synthesis. However, the total RNA pool may not be an accurate representation of the transcription potential of the genome, due to the post-transcriptional processing and regulation that an RNA undergoes after being transcribed. Thus, we believe that the use of nascent RNA-seq data (e.g. Bru-seq) is a closer approximation of the transcription potential of the genome, due to the temporal proximity of Bru-labeled RNAs to RNA synthesis, which is measured over 30 minutes (see Supplementary Methods, section 1.2). However, we also note that it is useful to include diverse transcriptional outputs when estimating the transcription potential of the genome where possible, due to the limitations inherent to each sequencing method. This allows more diverse RNA species to be represented in these estimates. For example, techniques that capture highly unstable RNA species (BruUV-seq) or small RNA species (miRNA-seq) add valuable information that other sequencing methods may miss.

In addition to the limitation mentioned above, previous estimates of pervasive transcription were calculated in an unstranded fashion, which does not encompass the unique transcription potential of each strand individually. Thus, we calculated coverage in a strand specific manner, increasing the denominator of our fractions to the total number of bases in the human genome (GRCh38, 6.2 billion bases). Because of this, the proportion of bases covered by sequencing reads that we report is smaller than previously estimated. However, we note that our estimates appear to be consistent with earlier estimates when approximating a stranded coverage value from the total unstranded value reported (e.g. 74.7% / 2 = 37.4% which is similar to Bru-seq and total RNA-seq stranded estimates). This method difference also highlights the importance of our result describing the total cumulative stranded coverage calculated from all DPCL assays (68.8%), as we are approaching these unstranded estimates while accounting for additional information.

Importantly, the method used to calculate the coverages reported is influenced by many factors. Anything from the sequencing read depth or data type to the number of samples or diversity of cell lines/tissues, or even the reference genome used, could influence the values reported. Furthermore, only uniquely mapping and processed alignments are used in our calculation, meaning repetitive regions are excluded. Thus, we emphasize that the coverage fractions reported here may be underestimates of the transcription potential of the genome. It is unclear how high the percent base coverage may actually be with more isogenic replicates, more diverse tissues and cell lines, greater sequencing depth, longer reads to facilitate mapping of repetitive regions, and a more accurate reference (e.g. T2T-CHM13[82]).

### Note 2: Using Bru-seq and BruUV-seq to study intergenic co-products of transcription

Nascent RNA-seq techniques can be used to better study RNA species that are more unstable than mRNA or other RNA species that are well-represented in the total RNA pool. Bru-seq captures the population of nascent RNA that is transcribed during a 30-minute window, where most sequencing reads are largely uniformly distributed throughout the gene due to the lack of splicing and intron degradation during this short labeling period[78] (Fig. 1d, Extended Data Fig. 1c). BruUV-seq[83] is a modification of the Bru-seq method where UVC inhibition of the RNA exosome coupled with UV-induced lesions lead to accumulation of sequencing reads over origins of transcription of both stable and unstable RNA species. This results in an enrichment of 5'-RNA signal at TSSs of coding and non-coding genes, as well as at highly unstable RNA species, like PROMPTs and eRNAs. Each of these techniques allow the exploration of the transcriptome temporally proximally to the polymerase traversing the gene, thus they are very useful for studying the patterns of transcription at RNA species that are expected to be relatively quickly degraded.

### Note 3: Factors influencing cell-to-cell variability in BruUV-seq data

BruUV-seq uses a high dose (100-200J/m$^2$) of UVC light (254 nm) to induce DNA lesions that prevent productive transcription elongation, in addition to inhibiting the activity of the RNA exosome complex. This results in an enrichment of 5'-RNA signal at TSSs of coding and non-coding genes, as well as at highly unstable RNA species, like PROMPTs and eRNAs. BruUV-seq offers the advantage of exploring these patterns of transcription/initiation in live cells by perturbing the mechanisms that turn-over premature transcription termination (PTT) transcripts[2], aberrant, and highly unstable RNA species. However, this also necessitates that this technique is somewhat dependent on the cellular response to being perturbed by UV and impairment of the RNA exosome.

We find that the patterns of 5'-signal enrichment seen across BruUV-seq datasets can be variable between cell lines and replicate samples (Extended Data Fig. 1e, 2a), which could result from either biological or technical mechanisms. However, the source of this variability is difficult to determine and could not be attributed to one mechanism. Cell lines can have different levels of basal transcription, and this can be further influenced by the cell's transcriptional response to UV exposure. Often cells will shut down transcription in response to DNA damaging agents, yet we still observe 5'-signal enrichment that allows the exploration of transcriptional patterns using BruUV-seq. This signal enrichment is thought to arise from two main UV-irradiation consequences, UV-induced elongation defects and RNA exosome stabilization, which lead to the redistribution of sequencing reads near TSSs in our assay. Upon UVC irradiation, DNA lesions, predominantly cyclobutane pyrimidine dimers and 6–4 photoproducts, are introduced which stall RNAPII elongation complexes[84]. These lesions are expected to be randomly distributed throughout the genome, generally occur every ~1 kb at the doses used for BruUV-seq[85] and are expected to be repaired by nucleotide excision repair (NER) on the order of hours to days[86,87]. UVC irradiation is also known to stabilize the RNA exosome, resulting in higher PTTs, and leading to increased 5'-enrichment of signal. This UV-induced stabilization of the RNA exosome is also known to enrich for PROMPT and eRNA species, given the inhibition of the rapid degradation of these species. The disruption of these mechanisms of transcription elongation and RNA degradation, while expected to affect transcription throughout the genome similarly, may be subject to cell line specific variability. For example, RNA exosome abundance and the inhibition of its degradation functionality could differ between cell lines, although the relative differences in RNA exosome biology between all DPCL cell lines are not reported here. Additionally, NER efficiencies can differ between cell lines[86], although these differences may not be as noticeable during the 30-minute labeling period of BruUV-seq. The cellular impacts on transcription and RNA degradation upon UVC irradiation highlight the potential for biological variability to influence BruUV-seq data. In addition, any technical or experimental variability could lead to overall differences in BruUV-seq data. For example, sequencing read depth can lead to less overall enrichment of RNA signal around TSSs, yielding less defined 5'-enrichment of signal that could influence some of these results. Overall, the presentation of this information is meant to increase transparency of the limitations of the BruUV-seq technique and to lend some potential explanations for the variability in BruUV-seq signal enrichment seen between cell lines.

Considering the potential sources of biological or technical variability listed above, we summarized several QC metrics across all cell lines in BruUV-seq and Bru-seq data, including 1.) the number of intergenic reads, 2.) the abundance of spike-in reads (see Supplementary Methods, sections 1.2.4 and 2.2), and 3.) the number of uniquely mapping reads. Although these metrics do not provide a complete explanation for this variation, they aid in our understanding of the data underlying our analyses and their potential contribution to the variable patterns seen in our PROMPT and eRNA results. Qualitative information about the cell lines and replicates can be gleaned from the increase in intergenic read density in BruUV-seq data as compared with Bru-seq, as we expect an accumulation of sequencing reads at unstable, intergenic transcripts after UV-irradiation (Supplementary Fig. 3a). The differences in global transcriptional levels after UV when compared to non-irradiated Bru-seq data, which are expected to be reduced under UV stress, can be extracted from the accumulation of labeled spike-in reads (Supplementary Fig. 3b). Along with these attributes, the uniquely mapping read count influences downstream analyses that are dependent on determining regions of high-confidence signal (Supplementary Fig. 3c). An indication of these possible, but not exhaustive, causes of cell line and replicate differences can be found in the dissimilarities in PROMPT fractions that are above an RPKM threshold of 0.1, or enhancer regions identified using peak calling methods that rely on statistical approaches to identify regions of enriched signal (Supplementary Fig. 3d-f).

Despite the observed variability, we find many similarities between cell lines in the transcription patterns of the PROMPT and eRNA regions assayed in this publication using BruUV-seq data. Although these unstable RNA species, especially enhancers, are expected to be more variable between cell lines, the patterns that we appreciated in the transcription of these RNA species genome-wide can be appreciated in individual cell lines where enough data is available. Thus, we are confident that the data used in this study was appropriate for the analyses performed.

### Note 4: Limitations of determining full-length PROMPT, eRNA, and RT transcription units

The methods used to capture regions of PROMPT, eRNA, and RT transcription from Bru-seq and BruUV-seq data are limited in their ability to capture full-length transcription units of an RNA species. In addition to the inherent limitations of short-read sequencing to provide information on a full-length transcript, our methods generally lack the resolution to accurately identify 5'- and 3'-end sites of a transcript. Furthermore, because a defining feature of the BruUV-seq method is the induction of DNA lesions that inhibit elongation, we do not expect to capture full length transcripts from this technique.

In addition to the limitations of Bru-seq and BruUV-seq described above, each of the methods used to capture our RNA species of interest further restricts our ability to capture full-length transcripts. PROMPTs definitions rely on a fixed region upstream of a genic TSS; thus, no length information is provided in that analysis. eRNA transcription is determined using BruUV-peaks, where peaks will only be called in regions of high signal. This means that, in addition to the exclusion of lowly expressed enhancers, the peak regions captured also do not extend to the 5'- and 3'-ends of the RNA signal appreciable in our data. Finally, RT transcription is identified using an HMM model which is designed to identify transcribed regions of the genome. While this is a powerful tool to define transcription units, this model relies on binned coverage data (250 bp), thus defining absolute start and end sites is not possible. These regions of PROMPT, eRNA, and RT transcription can be used to explore these RNA species and their patterns of expression but are not definitive annotations of their transcription units.

### Supplementary Methods

### 1. Cell Collection and Experimental Protocols

A minimum of 250 million cells were grown (see Supplementary Table 1) in 150mm dishes for each cell line replicate, with the exception of one 60mm dish used for ATAC-seq. For collection, adherent cells were washed once with PBS, then TrypLE Express was added for cell dissociation. Cells were counted and then aliquoted for the various techniques as described below. Suspension cells were counted and aliquoted. For PC-9 cells, which grow as a mixed culture of adherent and suspension cells, both the suspended and adherent cells were collected. All samples were stored at -80°C until distribution to the participating ENCODE consortium labs.

Experimental guidelines and data standards for most assays can be found on the ENCODE portal (https://www.encodeproject.org/data-standards/). These resources can be found linked to each experiment and/or sample, and are also linked in the following sections briefly describing the experimental protocols for each assay in the Deeply Profiled Cell Lines (DPCL) dataset (L2).

#### 1.1. ATAC-seq and snATAC-seq

A separate 60mm dish of cells was treated for 10 minutes with 200U/ml DNase (Worthington) from a stock of 40,000U/ml in Hank's Balanced Salt Solution. Cells were then collected, counted and 75,000 cells were transferred to a microcentrifuge tube. The cells were centrifuged for 5 minutes at 350xg at 4°C. The supernatant was carefully removed, and the cell pellet was resuspended in 100µl cold Bambanker freezing medium (Bulldog Bio), and stored at -80°C.

Cell pellets were used for both ATAC-seq and snATAC-seq, and these samples were processed by Michael Snyder's lab at Stanford University. Bulk ATAC-seq samples were processed using the Omni-ATAC protocol[88] (L3). For snATAC-seq, nuclei were isolated per the Snyder lab's protocol (L4) on the ENCODE portal, DNA was extracted, and scATAC-seq was performed using an adapted 10X genomics, chromium single cell ATAC protocol (L5).

#### 1.2. Bru-seq suite of techniques

One 150mm dish was used for each of the 5 Bru-seq treatments and/or techniques. The final cell counts per sample varied by cell line. All Bru-seq, BruChase-seq, and BruUV-seq samples were processed by Mats Ljungman's lab at the University of Michigan. Nascent RNA is labeled, isolated, and libraries are prepared based on published ENCODE protocols (L6, L7, L8, L9), the steps of which are also detailed below. These methods are updated versions of those previously published[80,81,83].

##### 1.2.1. Bru-seq

For Bru-seq, bromouridine (Aldrich) was added to the conditioned media for a final concentration of 2mM. After incubation at 37°C for 30 minutes, the media was aspirated, cells dissociated with TrypLE Express, counted, then pelleted by centrifugation at 1500rpm for 5 minutes. 3ml of Trizol was added to the cell pellet and the pellet was resuspended. Samples were stored at -80°C until RNA isolation and library preparation.

For the IR samples, the cells were treated with 5Gy of ionizing radiation using an IC-320 Biological Irradiator (Kimtron Inc., Oxford, CT) with a dose rate of ~2 Gy/min of x-rays, then incubated at 37°C for 1 hour. Bromouridine was added to a final concentration of 2mM, and cells incubated at 37°C for an additional 30 minutes before being collected the same as the Bru-seq samples.

#### 1.2.2. BruChase-seq

For BruChase-seq, bromouridine was added to the conditioned media and incubated at 37°C for 30 minutes. The bru-containing media was then aspirated, and cells washed once with PBS. Conditioned media containing 20mM uridine was added and the cells incubated at 37°C for either 2 hours or 6 hours before being collected and stored as above.

#### 1.2.3. BruUV-seq

BruUV-seq samples had their media removed and cells were washed with PBS before being exposed to 100J/m$^2$ (200J/m$^2$ for suspension cell lines) of UV light. Bromouridine was added to the reserved media to a final concentration of 2mM and immediately added back to the cells following UV-irradiation and incubated at 37°C for 30 minutes before being collected and stored as above.

#### 1.2.4. Stranded library preparation (all Bru-seq assays)

Total RNA Isolation:

After thawing Trizol samples, 0.6ml of chloroform was added to each Bru-seq sample and the sample shaken vigorously to mix. The sample was centrifuged at 12,000xg for 15 minutes at 4°C using a Sorvall Lynx centrifuge. The resulting top aqueous layer was carefully removed and transferred to a new centrifuge tube. 3ml of isopropanol was added and the sample centrifuged at 8,000xg for 15 minutes at 4°C. After aspirating the supernatant, the pellet was washed by adding 3ml of 75% ethanol and centrifuged at 8,000xg for 5 minutes at 4°C. The supernatant was again aspirated, and the tube inverted to allow the pellet to dry slightly. The pellet was then resuspended in 100µl of DEPC-treated water and transferred to an RNase-free microcentrifuge tube and incubated at 55°C for 10 minutes to ensure the pellet was completely dissolved. To remove any DNase, the RNA samples were treated with the Turbo DNA-free kit (Invitrogen). The resulting total RNA was then quantitated using a Nanodrop (ThermoScientific).

Bead conjugation:

50µl of goat anti-mouse Dynabeads (Invitrogen,11033) was used for each sample. The beads were washed 3 times with wash buffer (0.1% BSA in DEPC-PBS) and captured on a magnetic stand after each wash. After the final wash, the beads were resuspended in 200µl of wash buffer and 5µl (2.5µg) of anti-BrdU monoclonal antibody (BD Biosciences cat# 555627) was added along with 0.5µl of RNaseOUT (Invitrogen). The bead slurry was then incubated at room temperature for 1 hour with gentle rotation. The beads were again washed 3 times with wash buffer and resuspended in 200µl of wash buffer and 0.5µl of RNaseOUT.

Bru-RNA isolation:

100µg of each total RNA sample was added to a new RNase-free microcentrifuge tube and a spike-in cocktail was added. This cocktail consists of *in vitro* transcribed RNA from *A. thaliana* genes AGP23 (Bru-labeled), OBF-5 (Bru-labeled), PDF1 (unlabeled) and AP2 (unlabeled), total RNA extracted from *D. Melanogaster* (Dm) S3 cells (Bru-labeled), and total RNA extracted from *E.coli* K-12 MG1655 strain (unlabeled), and it was optimized to have final amounts as a percentage by weight of the sample total RNA. 25µl of the spike cocktail was added to Bru-seq samples, 6.25µl was added to BruChase-seq samples and 5µl was added to BruUV-seq samples. DEPC-water was added to bring each sample to a final volume of 200µl. After incubating at 80°C for 10 minutes, the samples were placed on ice to cool briefly before being transferred to a microcentrifuge tube containing the prepared conjugated beads. The RNA-bead mixture was incubated at room temperature for 1 hour with gentle rotation. The samples were then captured on a magnetic stand and the supernatant aspirated. 400µl of wash buffer was added and the samples were gently rotated for 1 minute at room temperature. The beads were captured again, and the supernatant removed. 200µl of wash buffer was added and the tubes were gently flicked to mix, after which the beads were captured, and the supernatant again removed. A final wash was done with 200µl of DEPC-PBS, after which the beads were captured, and the supernatant removed. To elute the Bru-RNA from the beads, 30µl of DEPC-water was added to the beads and the mixture was immediately transferred to a siliconized microcentrifuge tube and incubated in a 95°C heat block for 10 minutes with shaking. Samples were then briefly centrifuged, beads captured and the Bru-RNA containing supernatant was transferred to a new RNase free tube. Quantitation was done using a Nanodrop with DEPC-water as a blank. Bru-RNA was stored at -80°C.

NEB hairpin adapter and barcode preparation:

The sequences of the NEBNext hairpin adapter and primers used can be found in the NEBNext® Multiplex Oligos for Illumina® (Dual Index Primers Set 1) manual (https://www.neb.com/-/media/nebus/files/manuals/manuale7600.pdf?rev=aa1238754b164373a9a213119e2573d3&hash=FE525D5D7D0489B71B3300D3A3A12BF1). The NEBNext hairpin adapter was ordered from IDT and dissolved in IDTE pH 8.0 (IDT) to make a stock solution of 50µM and stored as 400µl aliquots. To the aliquot, 100µl of 10X Standard Taq Reaction Buffer (NEB B9015S) was added to bring the concentration to 40µM. The aliquot was then diluted to 10µM with IDTE 8.0 and further aliquoted into 20µl working stocks. Just prior to use, each working stock was annealed by incubating in a Thermomixer at 55°C for 1 minute, then allowed to slowly cool to room temperature in the Thermomixer. Any unused annealed adapter stock was discarded.

Eight i5 series barcoded primers and 12 i7 series barcoded primers were ordered from IDT with sequences based on the NEBNext Primers. Each primer was dissolved in IDTE 8.0 to a final working concentration of 10µM. 96 unique primer pairs were created, each containing an i5 primer and an i7 primer. 1.5µl of a unique primer pair mix was used per sample in the final PCR reaction, with care taken to avoid duplicating a primer pair mix when working with multiple samples.

Library Preparation:

For each library, 250ng of Bru-RNA was added to a PCR tube and brought to 16µl with UltraPure water. To remove any potential remaining rRNA and fragment the Bru-RNA, a mixture containing:

8µl 5x First Strand Buffer (from Superscript II kit, Invitrogen)
1µl Random Primer (Invitrogen)
0.1µl Fast Select (Qiagen)

was added to each sample. The samples were incubated in a thermal cycler with the following steps:

2 minutes at 75°C
2 minutes at 70°C
2 minutes at 65°C
2 minutes at 60°C
2 minutes at 55°C
5 minutes at 37°C
5 minutes at 25°C
hold at 4°C.

To synthesize the first strand cDNA, the following mixture was added to each sample:

6.9µl UltraPure water
4.0µl 100mM DTT (from Superscript II kit, Invitrogen)
0.8µl 25mM dNTPs
0.8µl ActinomycinD (2.5µg/µl)
0.5µl RNaseOUT, 2.0µl Superscript II

The samples were then incubated in a thermal cycler with the following steps:

10 minutes at 25°C
50 minutes at 42°C
15 minutes at 70°C
hold at 4°C.

The resulting single stranded cDNA was purified using magnetic RNAclean beads (Beckman Coulter). Each sample was transferred to a microcentrifuge tube containing 72µl of RNAclean beads and incubated in a Thermomixer (Eppendorf) for 10 minutes at 24°C with shaking at 550rpm. The beads were then captured on a magnetic stand and the supernatant removed by pipetting. With the tubes still in the stand, 200µl of 80% ethanol was added without disturbing the beads. After 30 seconds, the ethanol was removed, and the wash repeated a second time. After removing the second wash, the tubes were spun briefly, recaptured, and any remaining 80% ethanol removed. The open tubes were then placed in a 37°C Thermomixer to dry until the beads begin to show signs of cracking, after which 32µl of elution buffer (5mM Tris, pH 8.0) was added and the beads resuspended. The samples were then incubated in a Thermomixer for 6 minutes at 28°C with shaking at 550rpm, before the beads were captured and 30µl of the supernatant transferred to a new PCR tube. The second strand synthesis was performed by adding the following mixture to the newly isolated single strand cDNA:

26.48µl UltraPure water
15µl NEBuffer 2 (NEB)
0.9µl dG+dA+dU+dC nucleotide mix
0.75µl RNase H (Invitrogen)
1.875µl DNA Polymerase I (Invitrogen)

and incubating in a thermal cycler for 2 hours at 16°C. Following this step, the double stranded cDNA was purified using 112.5µl magnetic Ampure XP beads (Beckman Coulter), following the same bead purification protocol as above and eluted in 27µl of the elution buffer, with 25µl transferred to a new microcentrifuge tube for the next step. The samples were stored at -20°C unless proceeding immediately to the next step.

Next, End Repair was performed by incubating the cDNA with a mixture containing:

14.7µl UltraPure water
5.0µl T4 DNA Ligase Buffer
0.8µl 25mM dNTPs
2.0µl T4 DNA Polymerase (NEB)
0.5µl DNA Polymerase
Klenow Fragment (NEB)
2.0µl T4 Polynucleotide Kinase (NEB).

The samples were then incubated at 20°C for 30 minutes before bead purification with 75µl Ampure XP beads as described above. The elution was performed with 32µl 5mM Tris, pH 8.0 and 30µl was transferred to a new tube for the subsequent adenylation step. To adenylate, a mixture was made containing 11.5µl UltraPure water, 5.0µl NEBuffer 2, 1.0ul 10mM dATP and 2.5µl Klenow 3'-5' exonuclease (NEB). This mixture was added to the previous elution and the samples were incubated in a thermomixer at 37°C for 30 minutes. For the bead purification, 75µl of Ampure XP beads was added and the protocol followed as before, with the samples eluted using 20µl 5mM Tris, pH 8.0 and 18.5µl used for the adapter ligation.

Prior to the adapter ligation, the pre-aliquoted hairpin adapters (see above) were annealed by incubating in a thermomixer for 1 minute at 55°C, then allowing the aliquot to gradually cool to room temperature in the thermomixer before using. To ligate the adapters to the adenylated cDNA, NEB's Quick Ligation Kit was used. Briefly, a master mix containing 20µl of the Quick Ligase Buffer was mixed with 1µl of the Quick Ligase per sample. 1.5µl of the annealed adapter was added to each sample, along with 20µl of the Quick Ligase mix and the samples incubated at room temperature for at least 30 minutes. The samples were then purified using 40µl of Ampure XP beads following the usual protocol. 32µl of 5mM Tris, pH 8.0 was used in the elution step and 30µl was transferred to a new microcentrifuge tube for size selection.

5µl of 6x gel loading buffer (3% glycerol, 0.25% bromophenol blue, 5mM EDTA) was added to each sample and loaded onto a thick 3% NuSieve agarose gel (Lonza). A 100 bp ladder was loaded between samples. The gel was run for 1 hour, 20 minutes at 75V, in 1X TAE, then rinsed with distilled water before slices were excised at 500bp using Xtracta gel extractors (Promega). A second slice directly above the first excised slice was taken and saved at -20°C for backup. The 500bp gel slice was purified using the Qiaex II kit (Qiagen) and the DNA eluted in 42µl of 5mM Tris, pH 8.0. 20µl was saved at -20°C for backup, and 20µl was used for the subsequent PCR reaction.

1µl of USER enzyme (NEB) was added to each 20µl sample and incubated in a thermal cycler for 15 minutes at 37°C, then cooled to 4°C. A Master Mix was prepared consisting of 25µl of Phusion Master Mix (ThermoFisher) and 0.1µl Thermostable Inorganic Pyrophosphatase (TIPP, NEB) per sample. This mixture was added to the sample along with 1.5µl of a barcoded dual-index primer pair [link to supplemental primer info] and incubated in a thermal cycler using the following program:

98˚C for 30 seconds
11-14 cycles of:
    98˚C for 10 seconds
    60˚C for 30 seconds
    72˚C for 30 seconds
72˚C for 5 minutes
Hold at 4˚C.

The amplified library was then purified with 38µl of AmPure XP beads, following the same protocol as before, and eluted in 36µl of 5mM Tris, pH 8.0.

The final library was quantitated using a Nanodrop (ThermoScientific) and 3µl was run on a thin, 1.5% agarose gel to verify a single band of approximately 500bp. Libraries were submitted to the University of Michigan Advanced Genomics Core for 300 cycles of 150 base, paired-end sequencing (PE150) on an Illumina NovaSeq 6000.

#### 1.3. ChIA-PET

12 million cells were transferred to each of five 15ml centrifuge tubes. Cells were pelleted at 350xg for 5 minutes at 4°C and the supernatant aspirated. To crosslink, each cell pellet was resuspended in 1ml of 2% formaldehyde in Dulbecco's PBS, transferred to separate microcentrifuge tubes and incubated at room temperature for 20 minutes on a tube rotator. The crosslinking was quenched by the addition of glycine to a final concentration of 0.2M and the cells rotated at room temperature for an additional 10 minutes. After centrifugation at 5000rpm for 5 minutes at room temperature, the cell pellets were resuspended in 1ml Dulbecco's PBS, and the cells were counted, and the numbers recorded. The cells were again centrifuged at 5000rpm for 5 minutes at room temperature, after which the supernatant was aspirated and the cell pellets snap frozen in a dry ice/ethanol bath, then stored at -80°C. In the end, each ChIA-PET sample consisted of five cell pellets of approximately 10 million cells each.

ChIA-PET samples were processed by Charles Lee's lab at The Jackson Laboratories. Cell pellets were lysed, DNA was isolated, and libraries were constructed based on published protocols[89] (L10, starting with Part II).

#### 1.4. DNase-seq

One million cells were transferred to a microcentrifuge tube and pelleted by centrifuging for 5 minutes at 350xg at 4°C. The supernatant was aspirated, and the pellet resuspended in 1ml cold Bambanker freezing medium, then stored at -80°C.

DNase-seq samples were processed by John Stamatoyannopoulos' lab at the University of Washington. Nuclei were isolated (L11), DNA was extracted and DNase treated, and libraries were prepared using previously published protocols.

#### 1.5. Intact Hi-C

Six million cells were transferred to each of ten 15ml centrifuge tubes. Cells were pelleted at 350xg for 5 minutes at 4°C, the supernatant aspirated and the cells resuspended in 5ml fresh medium. To crosslink, formaldehyde was added to a final concentration of 1% from a 16% stock solution, and the samples were incubated at room temperature for 10 minutes on a tube rotator. The crosslinking was quenched by the addition of glycine to a final concentration of 0.2M and the cells rotated at room temperature for an additional 5 minutes. After centrifuging at 300xg for 5 minutes at 4°C, the cell pellets were resuspended in 1ml cold PBS, the cells were counted, and the cell counts recorded. The cells were again centrifuged at 300xg for 5 minutes at 4°C, after which the supernatant was aspirated and the cell pellets snap frozen in a dry ice/ethanol bath, then stored at -80°C. In the end, each Hi-C sample consisted of ten cell pellets of approximately 5 million cells each.

Intact Hi-C samples were processed by Erez Lieberman Aiden's lab at Baylor University. Cell pellets were lysed for intact Hi-C based on their published protocol (L12, starting at Module 2).

#### 1.6. PRO-cap

15 million cells were transferred to a 15ml centrifuge tube and pelleted by centrifuging for 5 minutes at 350xg at 4°C. All but 1ml of supernatant was aspirated and the cells were transferred to a microcentrifuge tube. The cells were again centrifuged for 5 minutes at 350xg at 4°C. The supernatant was aspirated, and the pellet was snap frozen in a dry ice/ethanol bath, then stored at -80°C.

PRO-cap samples were processed by Haiyuan Yu's lab at Cornell University. Run-on reactions were carried out, RNA was extracted, and libraries were prepared according to published protocols[90] (L13).

#### 1.7. RNA-seq

3 million cells were transferred to a microcentrifuge tube and pelleted by centrifuging for 5 minutes at 350xg at 4°C. The supernatant was aspirated, and the pellet was snap frozen in a dry ice/ethanol bath, then stored at -80°C. Cell pellets were used for all 3 mature RNA-seq experiments: long-read, miRNA-seq, and total RNA-seq.

##### 1.7.1. Long-read RNA-seq and miRNA-seq

Long-read and miRNA sequencing samples were processed by Ali Mortazavi's lab at University of California, Irvine. Long-read libraries were created for Pacific Biosciences sequencing per published protocols (L14), where cDNA was subject to poly-A selection and exonuclease treatment.

miRNA-seq samples were processed by the Mortazavi lab, with multiplexed miRNA libraries being created based on their published ENCODE protocol (L15) where miRNAs are size-selected from total RNA to be 30 bp or smaller.

#### 1.7.2. Total RNA-seq

Total RNA-seq samples were processed by Barbara Wold's lab at California Institute of Technology. Total RNA was extracted and libraries were prepared based on published protocols[91] (L16).

### 2. Sequencing and Data Processing

The ENCODE Data Coordinating Center (DCC) has developed Uniform Processing Pipelines for most major assay types generated by the project[71]. Details about the DCC Uniform Processing Pipelines can be found on the ENCODE portal (https://www.encodeproject.org/pipelines/) and code is available on the ENCODE DCC GitHub (https://github.com/ENCODE-DCC). These resources can be found linked to each experiment and/or sample and are also linked in the following sections briefly describing the experimental protocols and data processing for each assay in the Deeply Profiled Cell Lines (DPCL) dataset. Not all data processing pipelines are currently among the Uniform Processing Pipelines developed by the DCC, but these are described below in more detail. All data discussed in this section is available on the ENCODE portal or is hosted on AWS, see Supplementary Data File 1 for details on data availability.

#### 2.1. ATAC-seq and snATAC-seq

ATAC-seq libraries were sequenced on the Illumina HiSeq 4000 platform (100 bp paired-end reads) to a depth of ~50 million reads (https://www.encodeproject.org/data-standards/atac-seq/atac-encode4/). ATAC-seq data was processed using the DCC uniform processing pipeline[92] (https://github.com/ENCODE-DCC/atac-seq-pipeline). For our analyses, we have primarily used the conservative IDR peaks derived from biological replicate samples.

snATAC-seq libraries were sequenced on the Illumina NovaSeq 6000 platform (50 bp paired-end reads). ATAC-seq data was processed using a lab custom pipeline which is available on GitHub (https://github.com/kundajelab/ENCODE_scatac). This data was not incorporated into any analysis for this paper but could be integrated with other data if processed into pseudo-bulk data or via other integrations of single-cell and bulk datasets.

#### 2.2. Bru-seq, BruChase-seq, BruUV-seq

Strand-specific Bru-seq libraries (including BruChase-seq and BruUV-seq) were sequenced on the Illumina NovaSeq 6000 platform (150 bp paired-end reads) to a depth of ~50 million reads. Samples were sequenced to a depth of approximately 50 million read pairs. These data were mapped using a single custom sequencing pipeline that contains additional steps for BruUV-seq data only (L18). First, raw sequencing reads were trimmed (bbTools/bbduk) to remove adapter sequences. Reads not rejected during trimming were then pre-aligned to the premap reference set, which includes 1.) ribosomal RNA (rRNA) repeating unit (ENCSR612DCK; GenBank U13369.1), 2.) the human mitochondrial genome (chrM; ENCSR884DHJ), 3.) the EBV genome (ENCSR992NKC; RefSeq NC_007605.1), and 4.) select spike-in references for fruit fly (dm6) rRNA and mitochondrial genome (ENCSR817UPT). Pre-alignment was carried out using Bowtie2 and samtools, and any reads aligning to these sequences were removed. Reads not aligned to the premap references were then aligned to the main reference set, which includes GRCh38/GENCODE v29 (ENCSR884DHJ), and the following spike-in references (ENCSR817UPT): 1.) the fruit fly genome (dm6), 2.) the *E. coli* genome (K-12 MG1655), and 3.) *in vitro* transcribed *A. Thaliana* RNA sequences. Alignment was performed using STAR (v2.7.0f) and samtools producing unfiltered alignment files (BAM).

Alignments from genome BAM files were used to generate signal (bigWig) files separately for plus and minus strand data for all canonical chromosomes using deeptools. BigWigs were generated for all reads (including multimapping reads) and uniquely mapping reads only (MAPQ ≥ 255) and were signal normalized by RPKM using the total number of uniquely mapped reads as the total read depth. Duplicated reads were not filtered out, nor were any regions of the genome removed or filtered (e.g. problematic regions[93]).

Strand-specific coverage values were calculated per base and in 1 kb bins from uniquely mapping reads (MAPQ ≥ 255) using bedtools. Strand-specific read coverage per base was determined by intersecting the alignment with the reference sequence and was reported as 1/read length, where genomic bases with no read overlap are not reported. Binned coverages (1 kb) were calculated by intersecting binned regions with the base coverages and recalculating the coverage for each bin (sum of per-base coverages / bin size), where similarly bins with no read overlap are not reported. These coverage values were then used to generate various downstream files, including transcription segmentation files (see Methods: RT segment identification) and BruUV-peak files (BruUV-seq data only, see Methods: BruUV-seq peak calling…). In addition, these coverages were used to obtain counts over genes or other regions of interest (e.g. PROMPT, eRNA, and RT regions) that are used as input to many downstream analyses.

Counts for a region of interest were obtained by calculating a fractional coverage of sequenced reads over each base in a strand-specific manner and then summing the fractional coverages along the entire region[80]. The fractional coverage is calculated as the number of reads overlapping a base on each strand divided by the sequenced read length (SRL). The SRL is determined by Read1 and Read2 overlaps. If there is no read overlap, the SRL is 300, which is the sum of the length of the two reads. If Read1 and Read2 overlap, the SRL is the sum of the reads' overlap length and the length of the read regions that do not overlap. Fractional coverages are then typically rounded for use in count-based analyses.

### 2.3. ChIA-PET

ChIA-PET libraries were sequenced on the Illumina Novaseq 6000 platform (150 bp paired-end reads) to a depth that results in at least 20 million usable fragments (https://www.encodeproject.org/chia-pet/). ChIA-PET data was processed using the Ruan lab ChIA-PIPE pipeline[94] (http://github.com/TheJacksonLaboratory/ChIA-PIPE). These samples were run through ChIA-PIPE individually without a ChIP-seq input control, thus using MACS2 as a peak caller (L17).

Due to this method of processing, we performed the following additional steps to merge replicates and obtain relevant ChIA-PET loops. Replicate consensus peaks were obtained from intersecting individual replicate peaks using bedtools intersect (-f 0.5 -F 0.5). Each replicate consensus peak was overlapped with both loop anchors separately using bedtools intersect with default options. The loop anchor and replicate consensus peak information was then compiled per loop to obtain a comprehensive set of ChIA-PET interactions. Loops, where both anchors overlapped each other, were removed from the analysis. A more optimal method of processing these data would be to run ChIA-PIPE on merged replicate fastq files with an input control, and then performing downstream steps to obtain loops overlapping ChIP-seq peaks.

### 2.4. DNase-seq

DNase-seq libraries were sequenced on the Illumina Novaseq 6000 platform (100-150 bp paired-end reads) to a depth of ~50 million reads (https://www.encodeproject.org/data-standards/dnase-seq-encode4/). DNase-seq data was processed using the DCC uniform processing pipeline (https://github.com/ENCODE-DCC/dnase-seq-pipeline). For our analyses, we used 0.1% FDR narrow peak files and merged replicates by obtaining consensus peaks from both samples that overlapped by at least 50% using bedtools intersect (-f 0.5).

### 2.5. Intact Hi-C

Intact Hi-C libraries were sequenced on the Ultima Genomics platform (single-end reads) to a depth of ~2 billion total reads (https://www.encodeproject.org/hic/). Intact Hi-C data was processed using the DCC uniform processing pipeline (https://github.com/ENCODE-DCC/hic-pipeline) based on Juicer (https://github.com/aidenlab/juicer). For our analysis, we have used both genome compartments (5 kb bin size, mapq30 annotated), contact domains (mapq30), and loops (mapq30) files available on the encode portal.

Contact domain (TAD) boundaries (used in section 7.5) were derived by obtaining the coordinates from contact domain bedpe files, which represented the start and end of the entire domain, and constructing a bed file where the “boundary” coordinates for each domain are defined as the 1 kb region flanking each start/end point (e.g. start coordinate +500 bp and start coordinate -500 bp yields the upstream boundary). This was done to provide a boundary window at each domain end, similar in concept to a Hi-C loop anchor, to allow additional flexibility for downstream overlap analyses.

### 2.6. PRO-cap

Strand-specific PRO-cap libraries were sequenced on the Illumina HiSeq X Ten platform (150 bp paired-end reads) to a depth of ~50 million reads. PRO-cap data was processed using the Yu lab custom pipeline and PINTS peak calling tool[95] (https://github.com/hyulab/PINTS). For our analyses, we obtained all PRO-cap peaks by merging unidirectional and bidirectional peak outputs from PINTS. To do this, bidirectional peak pairs identified by PINTS were separated into distinct peaks and were merged into one file.

#### 2.7. Long-read RNA-seq

Strand-specific long-read RNA-seq libraries were sequenced on the Pacific Biosciences Sequel II platform to a depth of ~600,000 full-length nonchimeric reads (https://www.encodeproject.org/rna-seq/long-read-rna-seq/). Long-read sequencing was performed using the DCC uniform processing pipeline (https://github.com/ENCODE-DCC/long-read-rna-pipeline) based on TALON[96] (https://github.com/mortazavilab/TALON). For our genome coverage analysis, we used filtered alignment files (.bam).

#### 2.8. microRNA-seq

Unstranded microRNA-seq (miRNA-seq) libraries were sequenced on the Illumina NextSeq 2000 platform (75 bp single-end reads) to a depth of ~5 million reads (https://www.encodeproject.org/microrna/microrna-seq-encode4/), where mapped read lengths should be a minimum of 16 bp. miRNA-seq data was processed using the DCC uniform processing pipeline (https://github.com/ENCODE-DCC/mirna-seq-pipeline). For our genome coverage analysis, we used unfiltered alignment files (bam).

#### 2.9. Total RNA-seq

Strand-specific total RNA-seq libraries were sequenced on the Illumina HiSeq 2500 platform (100 bp paired-end reads) to a depth of ~30 million reads (https://www.encodeproject.org/data-standards/encode4-bulk-rna/). Total RNA-seq data was processed using the bulk RNA-seq pipeline that was developed as a part of the DCC uniform processing pipeline (https://github.com/ENCODE-DCC/rna-seq-pipeline). For our genome coverage analysis, we used unfiltered alignment files (.bam).

### 3. Genome coverage

#### 3.1. Fraction of genome covered by PROMPT, eRNA, and RT regions

To assess the fraction of intergenic space that was covered by the regions of PROMPT, eRNA, and RT signal that were identified in this study, we first removed any regions from the bed files for each RNA species that overlapped an annotated gene (GENCODE v29 or v44) per cell line (bedtools subtract -s). This was only applicable to the eRNA and RT regions which could overlap annotated genes. We then concatenated and merged the remaining regions for each RNA from all cell lines (bedtools merge -s). Finally, regions for all three RNA species were merged for all cell lines ("All") as well as for each cell line individually. The lengths (bp) of the resulting regions from each bed file were calculated and summed to obtain the number of intergenic bases that were covered by all regions identified. We then divided each value by the total number of intergenic bases (4418874935, see Methods: Genome coverage calculations) to obtain the fraction of bases covered by the regions of intergenic transcription identified in our analyses (Supplementary Fig. 1e).

#### 3.2. Overlap of PROMPT, eRNA, and RT region with GENCODE v44 annotation

Regions of intergenic PROMPT, eRNA and RT signal identified in our analysis were overlapped with annotated genes from the GENCODE v44 annotation using bedtools intersect using the following parameters: -s, -f 0.9 -F 0.9 -e. Based on these parameters, we found 472/12939 PROMPT regions, 1263/32144 eRNA regions (denominator is only intergenic regions, see Supplementary Methods, section 3.1), and 1280/18244 RT regions that overlapped an annotated gene in the most recent GENCODE version available.

### 4. Bru-seq and BruUV-seq QC

#### 4.1. Quality control metrics

All uniquely mapping reads (MAPQ ≥ 255), as well as those mapping to intergenic regions and labeled spike-ins were counted using custom scripts. For the intergenic and spike-in unique reads, their relative proportion to all uniquely mapping reads in the standard chromosomes or the complete set of chromosomes (standard + spike-in), respectively, were calculated. The values for these metrics were determined for all cell lines and their replicates in Bru-seq and BruUV-seq assays (Supplementary Fig.3a-c)

#### 4.2. 5'-enrichment score

For genes 30 kb and above, Bru-seq and BruUV-seq expression values (RPKM) were obtained for sectioned parts of the gene i.e., for TSS to TSS+15 kb and TSS+15 kb to TSS+30kb. All genes were required to be expressed above an RPKM threshold of 0.1 in all the cell lines. Per gene in each cell line replicate, the 5'-enrichment score was calculated as a ratio between the contiguous sections of the gene: 5'-enrichment score = (TSS to TSS+15 kb RPKM) / (TSS+15 kb to TSS+30 kb RPKM). A median score was computed per cell line replicate and assay (Extended Data Fig. 1e).

#### 4.3. Principal component analysis (PCA) of regions of interest

To assess signal variability in cell line replicates, we performed PCA for gene, PROMPT, enhancer, and RT regions using Bru-seq or BruUV-seq counts (Supplementary Fig. 4). Counts were obtained for bed regions as described in Supplementary Methods section 2.2. Counts were normalized using the variance stabilizing transformation (vst) function in R/DESeq2 (v1.36.0), and the variance was determined from all regions. The principal components with the highest variance (PC1 and PC2) were plotted using the plotPCA function in R/DESeq2 for all the cell lines and replicates.

For genes, Bru-seq counts were obtained for the full length of the gene, however, only TSS-proximal counts in the first 2 kb of the gene (TSS+2kb) were used to compare BruUV-seq data, due to the lack of gene body signal expected in this data type (Supplementary Fig. 4a-b). BruUV-seq counts were obtained for all PROMPT regions (see Methods: TSS curation…), as well as enhancer consensus loci (see Supplementary Methods, section 6.3) that do not overlap any annotated genes (GENCODE v29), to prevent gene body signal from influencing this analysis (Supplementary Fig. 4c-d). Bru-seq counts were obtained in the 5 kb region downstream of the TES (TES+5kb) for a subset of genes that were assigned an RTseg in our analysis and do not have another annotated gene within 5 kb downstream (Supplementary Fig. 4e). We observed a high replicate concordance, and the variability between samples was generally low (PC1 = 18-24% variance, PC2 = 11-13% variance).

### 5. PROMPTs

#### 5.1. Metagene plots

Strand-specific RPKM-normalized individual or merged replicate bigwigs generated from uniquely mapping reads (MAPQ ≥ 255) per cell line from BruUV-seq and Bru-seq data were used to create the metagene plots for all 16 cell lines. Out of the 12939 TSSs, genes that were at or above 5 kb in length, as well as those that did not have any annotated genes on either strand (any biotype, GRCh38 GENCODE v29) in a 5 kb region upstream of the TSS were selected for this analysis (5615). For each stranded bigwig, the score per genomic regions of interest was calculated for plus (2876/5615) and minus strand genes (2739/5615) separately using deeptools computeMatrix reference-point -S --referencePoint TSS --upstream 5000 --downstream 5000 --binSize 50 --sortRegions descend --sortUsing median --averageTypeBins median. Per cell line, using deeptools computeMatrixOperations rbind -m, sense signal score matrices were produced by combining the score matrices generated from the following: plus strand genes + plus strand bigwigs, minus strand genes + minus strand bigwigs, and for antisense signal score matrices: plus strand genes + minus strand bigwigs, minus strand genes + plus strand bigwigs. Profile plot genomic scores were calculated for sense and antisense matrices per cell line using deeptools plotProfile -m --averageType mean --refPointLabel TSS. These scores were then used to produce the final metagene plot in R/ggplot2 (Merged replicates, BruUV-seq data used in Extended Data Fig. 2a, and individual cell line replicates, BruUV-seq and Bru-seq data used in Supplementary Fig. 2a).

Histone metagene plots were generated from signal p-value bigwigs for the cell lines that had data for all modifications (10/16) (Supplementary Data File 1), for a 1 kb region up-and downstream of bidirectional TSSs in all cell lines (1041). Per cell line, the score per genomic regions of interest was calculated using deeptools computeMatrix reference-point -S --referencePoint center --upstream 1000 --downstream 1000 --binSize 10 --sortRegions descend --sortUsing median --averageTypeBins median. Profile plot genomic scores were calculated for the matrices per cell line using deeptools plotProfile -m --averageType mean --refPointLabel TSS. These scores were then used to produce the final metagene plot in R/ggplot2 (Extended Data Fig. 3b, top panel). Additionally, the signal profile for H3K79me2 was plotted for bidirectional and unidirectional enhancers (including the bidir_low_3 enhancer class, see Supplementary Data File 4) for selected cell lines (Extended Data Fig. 5g-i).

#### 5.2. TSS generation and signal characterization

The current definition of PROMPTs is based on their proximity to a gene, with these transcripts produced from upstream, divergent promoters in a nucleosome-depleted region shared with genes. In this paper, a fixed 2 kb divergent region upstream of the annotated gene

TSS, that represented the majority of the PROMPT signal, was used to characterize the PROMPTs. The region's length is in line with previous observations regarding where we tend to see signal[17], however, we do observe PROMPT transcription extending past this region, up to 5 kb and potentially beyond (Extended Data Fig. 2a). For this analysis, we focused on the transcripts generated upstream of RNAPII-directed genes, although PROMPTs originate upstream of all three RNAP products[97]. Due to the 5'-end signal enrichment we observe in BruUV-seq, we restricted the region where we investigated the genic signal to a TSS-proximal 2kb region. Annotated protein-coding genes and transcripts in the GRCh38 GENCODE v29 basic annotation were used for this analysis, resulting in multiple TSSs per gene which, if within 2 kb of each other, were consolidated into a single TSS unit. This approach to creating a merged unit of adjacent annotated start sites and designating the leftmost coordinate as the primary TSS was largely adopted due to the limited base-pair resolution information from BruUV-seq, thereby making it challenging to confidently determine the exact origin of transcription. Employing long-read technologies in conjunction with targeted enrichment of these unstable species such as auxin-mediated degradation of the components of the RNA exosome, particularly DIS3, will aid in estimating accurate transcription start sites as well as provide a better understanding of TSS-proximal signal.

To determine the transcriptional signal, PROMPT expression was obtained from BruUV-seq data largely due to UV-associated stabilization of transcripts that are typically degraded by the RNA exosome. Due to the potential consequences of using UV irradiation (Supplementary Note 3), for higher genic signal confidence, we used both BruUV-seq and Bru-seq data to get gene expression values. Nonetheless, BruUV-seq demonstrated a very similar measure of TSS-proximal gene expression to that of Bru-seq, as shown by the high degree of agreement between the signals in the two assays (Supplementary Fig. 3g). For our expression metrics, our method of determining high-confidence regions was based on a threshold of 0.1 RPKM, differing from other statistical methods such as peak calling to identify significantly enriched regions. By binning the standard chromosomes (1-22, X and Y) into 2 kb regions using bedtools makewindows -g -w 2000 for plus and minus strands, and calculating their RPKM values, we found that the 0.1 RPKM cut-off was considerably above the median stranded genomic 2 kb bin signal across all cell lines (Supplementary Fig. 5a), as well as above very lowly expressed regions for BruUV-seq and Bru-seq data (Supplementary Fig. 5b-c). For quality control (QC) purposes, a parallel classification of PROMPTs was performed based on their expression where they were designated as high-confidence (BruUV-seq RPKM > 0.1) or low-confidence (BruUV-seq RPKM ≤0.1). To validate this classification against an annotation-independent method of identifying regions of signal, we overlapped the PROMPT regions with peaks called in the BruUV-seq dataset (see Methods: BruUV-seq peak calling and eRNA identification) and observed that our classifications highly agree with the peak overlap patterns (Supplementary Fig. 6a). Although there are PROMPTs in the high-confidence class that do not overlap a peak, having shown that our RPKM threshold is above the background genomic signal, we proceeded to use all the regions categorized using the RPKM thresholds for a more inclusive set of regions to analyze.

Using the full set of classifications, we plotted the PROMPT and TSS-proximal gene RPKM distributions (Supplementary Fig. 6e). The presence of signal at or below our RPKM cut-off is potentially ambiguous, which may indicate technical artifacts or real signal that could be resolved upon deeper sequencing. Due to this reason, any classification that had a low-confidence PROMPT signal was called low-confidence bidirectional. Interestingly, for both bidirectional and low-confidence bidirectional categories, we find a smaller proportion of TSSs that have genic signal above our criteria cut-offs in either BruUV-seq or Bru-seq (Extended Data Fig. 2c-d). While exonic signal could be contributing to the presence of intragenic TSSs in these categories, many of the examples we observed were of TSSs with enhancer-like profiles (data not shown). We also observed that for the High PROMPT, low gene signal category, most of the PROMPT signal was due to continuing upstream transcription from longer PROMPTs or eRNA. There could be instances of intentional expression of only PROMPTs, potentially, in the pursuit of acquiring more gene-like functional properties. Although beyond the scope of this paper, a future systematic analysis investigating diverging and emergent characteristics of all PROMPTs would be of relevance, shedding light on the evolution of genes overall.

#### 5.3. QC analysis of PROMPT regions

The upstream divergent PROMPT region selected for our analysis includes the nucleosome-depleted region that is critical for transcriptional activation. To determine the prevalence of accessible chromatin regions as well as their relationship to PROMPT expression, the 2kb PROMPT regions were overlapped with DPCL ATAC-seq peaks (cell line-specific unstranded IDR conservative peaks, Supplementary Data File 1) or DNase-seq peaks (FDR thresholded peaks < 0.1) using bedtools intersect with default overlap options. To determine the agreement with an orthogonal nascent RNA-seq technique, PRO-cap, the closest upstream peak within 1 kb of the PROMPT regions was determined with bedtools closest -s -id -D -t first. The peak overlaps were largely in agreement with the PROMPT expression classifications (Supplementary Fig. 6b-d). We find that a smaller fraction of PROMPTs (below 25%) shows results that do not match our expectations. The presence of peaks overlapping, or in the proximity of, such regions in either PROMPT class could be a result of low-level transcriptional or chromatin accessibility signal in the PROMPT region, or could be a reflection of the transcriptional profile of the associated gene being different from the PROMPT's profile.

#### 5.4. PROMPT:Gene (P:G) ratio correlations (expanded)

For bidirectional TSSs across all cell lines (1041), Spearman correlation coefficients were calculated between log10 transformed P:G expression ratios and the various expression, chromatin and sequence parameters listed below. This was done in R using the cor.test function (method = "Spearman", exact = FALSE), and FDR adjusted p-values were calculated from p-values using the p.adjust function (method="BH") represented by $^{ns}p_{adj.} > 0.05$, $^{*}p_{adj.} < 0.05$, $^{**}p_{adj.} < 0.01$, $^{***}p_{adj.} < 0.001$, $^{****}p_{adj.} < 0.0001$. No TSSs were excluded from this analysis as a result of infinite or NaN ratio values. To determine if our method of merging TSSs that were within 2 kb of each other was influencing the results, we performed this analysis with a smaller set of single-isoform genes (234/1041) and found a high degree of agreement between the two correlation results (Supplementary Fig. 8d).

##### 5.4.1. PROMPT and genic signal

Log10 transformed TSS- proximal PROMPT or gene expression (RPKM) was used to determine the correlation with log10 P:G expression values. The moderate correlations that we observe indicate varying levels of expression values associated with these ratios.

##### 5.4.2. DNA accessibility signal

To determine open chromatin signal, counts were determined from ATAC-seq (GM12878, HCT116, HepG2, IMR90, K562, MCF-7, panc1, PC3) or DNase-seq (A673, Caco2, Calu3, HMEC and PC9) signal p-value bigwigs using deeptools multiBigwigSummary for a 500 bp region up-and downstream of the gene TSS. Log10 normalized signal counts (CPM) were used to determine the correlation with log10 P:G expression values.

##### 5.4.3. GC% ratio in initiating regions

The underlying nucleotide content (GC% and AT%) for three PROMPT and gene regions was obtained using bedtools nuc -s -seq, using the gene TSS as the reference. These values were obtained for the gene in: Region 1 (TSS+50 bp), Region 2 (TSS+500 bp) and Region 3 (TSS+500 bp to TSS+2000 bp). For the PROMPT, the sequence composition was obtained for comparable regions while excluding the NDR (100 bp is an average estimate of the NDR for the TSSs in this analysis): Region 1 (TSS-100 to TSS-150 bp), Region 2 (TSS-100 to TSS-600 bp) and Region 3 (TSS-600 bp to TSS-2100 bp). In Extended Data Fig. 2h, for uniformity, the regions are represented by the gene region designations for both features, with schematics to indicate the region per feature. Wilcoxon-signed rank tests were performed between the three regions in the gene and PROMPT regions using the compare_means function (method = “wilcox.test”, p.adjust.method = "BH", R/rstatix). False Discovery Rate (FDR) adjusted p-values were calculated from p-values and represented by $^{ns}$padj. > 0.05, *padj. < 0.05, **padj. < 0.01, ***padj. < 0.001, ****padj.< 0.0001. Additionally, we calculated the P:G ratio of the nucleotide content of these three regions. The P:G region 2 GC% log10 ratio was used to determine the correlation with log10 P:G expression values.

##### 5.4.4. Number of promoter-promoter (p-p) contacts

Promoter-promoter (p-p) contacts between the 12939 TSSs and an expressed protein-coding or lincRNA gene (TSS-proximal 2 kb BruUV-seq RPKM > 0.1) were determined using bedtools pairToBed to overlap loop anchors obtained from Intact Hi-C or POLR2A ChIA-PET data with regions 1kb up- and downstream of the two sets of TSSs. Self-interacting promoters were removed, and reciprocal promoter-promoter interactions were retained. The number of p-p contacts per TSS were used to determine the correlation with log10 P:G expression values.

##### 5.4.5. Histone post-translational modifications

Selected histone post-translational modifications namely H3K9ac, H3K27ac, H3K4me1, H3K4me2, H3K4me3, H3K27me3, H3K36me3, H3K79me2 for the cell lines that had data for all the modifications (10/16) were used for this analysis. TSS-proximal histone signal (counts) was obtained from signal p-value bigwigs for the various histone post-translational modifications (Supplementary Table 1) using deeptools multiBigwigSummary for gene (TSS+1kb) and PROMPT regions (TSS-1.1kb, 100 bp is an average estimate of the NDR for the TSSs in this analysis). These regions were determined from the histone aggregate plots (Extended Data Fig. 3b). Normalized counts (CPM) were generated for these regions and log10 transformed normalized counts were used to determine the correlations of each histone modification's signal to transcriptional levels, as well as to the signal of the different histone modifications for the gene and PROMPT region individually. We observe moderate correlations with expression, and moderate to high correlations between several histone modifications (Supplementary Fig. 9a-b). Additionally, normalized CTCF signal counts (CPM) were obtained using deeptools multiBigwigSummary for a region 350 bp upstream of the gene TSS to correlate with PROMPT expression (Supplementary Fig. 9a-b).

The histone P:G ratios were calculated as follows: Histone P:G ratio = TSS-1.1kb PROMPT CPM / TSS + 1 kb gene CPM. Log10 transformed ratio values were used to determine the correlation with log10 P:G expression values. Along with H3K9ac, H3K27ac, H3K4me3 and H3K79me2, H3K4me2 showed a mild correlation with PROMPT expression (Supplementary Fig. 9a). This is not entirely reflected in the dynamic correlations with the PG ratios potentially due to differences in the deposition of this mark on +1 nucleosomes in the PROMPT regions compared to genes (Supplementary Fig. 9a).

#### 5.5. eTSS considerations

Our method of identifying gene promoters with potential enhancer-like properties (eTSSs) did not require any intrinsic regulatory element expression. eTSSs were selected based on the presence of histone modifications typically associated with transcriptionally active enhancers, as well as interactions with distally expressed protein-coding or lincRNA genes. However, we found confirmation of activity based on the expression profiles at eTSSs, where they were primarily bidirectional or showed high-confidence signal in the gene body proximal to the TSS (Extended Data Fig. 5f). This can be contrasted with the TSS category distribution for all promoter-promoter contacts, where we see a higher fraction of TSSs with low-confidence signal (Supplementary Fig. 9c), most notably the negligible signal TSSs. Although not the majority, this suggests that genomic regions engaging in distal contacts with expressed genes do not necessarily have to be active, either reflecting similarities with inactive/poised enhancers or indicating differential roles for gene TSSs engaged in distal interactions.

For all the eTSSs that made distal contacts in POLR2A ChIA-PET or Intact Hi-C data, we found that the linear interaction distances (measured as the distance between loop anchor midpoints of the interacting regions) range from ~8-1000 kb, with the median distance across cell lines at 100 kb (Supplementary Fig. 9d). Since co-regulation of expression is a feature of enhancer-target gene interactions, we investigated whether such a relationship could be observed between an eTSS and its distally interacting gene. For target genes that interacted with only one eTSS identified in either POLR2A ChIA-PET or Intact Hi-C data (Extended data Fig. 7e, one:eTSS group) and out of this group, after extracting eTSSs that interacted with only one target gene, we found low correlations between the expression of the eTSS gene or

PROMPT and the target gene (Supplementary Fig. 9e). This result aligns with previous findings[76], and further studies exploring a concordant transcriptional response to exogenous stress or high-throughput candidate screening assays, such as massively parallel reporter assays (MPRAs) or STARR-seq[98], will be essential in determining whether the eTSSs we identified truly have distal regulatory potential.

### 6. Enhancers/eRNA

#### 6.1. BruUV-peak and enhancer peak validation and feature summary

Our method of calling narrow peaks using default MACS2[32] parameters was determined to be appropriate due to the tendency of distinct peaks of RNA signal to form at TSSs (e.g. 5'-ends of genes). However, given that gene body RNA signal can often still be detected in BruUV-seq, refinement of this method could be undertaken to further reduce the prevalence of peaks called over noisy gene body regions. While further exploration into potential modifications to the parameters used for peak calling was not performed during this study, we validated the relevance of the BruUV-peaks that were called using the data that was generated by the ENCODE4 consortium that could confirm these are regions of broadly relevant transcription. For these analyses, we considered all BruUV-peaks discovered in any cell line or all BruUV-peaks found per cell line and found that the features of these peaks were very similar between cell lines. We also considered only the subset of intergenic BruUV-peaks, which were defined as not overlapping an annotated gene in the GENCODE v29 annotation, to explore a subset of peaks that does not include peaks of gene body signal that may not indicate regions of 5'-signal enrichment (transcription initiation) as these could be sources of noise in the data.

To initially validate the performance of our method of peak calling from BruUV-seq data, we summarized some of the features of the BruUV-peaks and enhancer peaks that we called with our method. We found that BruUV-peaks range in size from ~500 bp to ~8 kb and have a median length of ~1.5 kb (Supplementary Fig. 10a). By exploring peaks overlapping dELS cCREs (eRNA signal) and PLS cCREs (gene signal), we confirmed that peaks over gene promoters are longer on average than peaks at enhancer elements (Supplementary Fig. 10a). This matched our expectations of BruUV-signal at each of these genomic regions; more and broader signal is expected at gene promoters where transcription initiation occurs more frequently and where productive elongation occurs under unperturbed conditions. Because eRNAs are known to be much shorter than protein-coding genes, it was expected that these BruUV-seq enhancer-peaks would likely be shorter overall. This was true for all peaks as well as intergenic peaks only (Supplementary Fig. 10a), and for all individual cell lines with little variability observed between them (Supplementary Fig. 10d-e). While most BruUV-peaks overlapping dELS cCREs were short, we did note the presence of longer peaks in this data. These longer peaks have not been exhaustively examined, however, many of these may be the result of the technical limitations of our peak calling algorithm to accurately resolve regions of nearby enhancers that are all transcribed (e.g. super-enhancers). We also note that some of the longest examples of BruUV-peaks overlapping dELS cCREs are also unique to gene body regions, thus contaminating gene signal may also result in longer peaks.

We further established the relevance of all and intergenic BruUV-peaks by calculating the fraction of peaks that overlap each of the ENCODE4 cCREs[33], regions of open chromatin (ATAC-seq, DNase-seq), and nascent RNA-seq peaks from an orthologous method, PRO-cap (Supplementary Fig. 10b-c). cCRE, ATAC-seq peak (Supplementary Methods, section 2.1), and DNase-seq peak (Supplementary Methods, section 2.4) overlaps were obtained using bedtools intersect. For PRO-cap peaks (Supplementary Methods, section 2.1), we used bedtools closest (-id -s -D a) to obtain PRO-cap peaks that were within 300 bp upstream of a BruUV-seq peak. Due to the very narrow footprint of PRO-cap peaks that occur directly at a TSS, and the tendency of BruUV-seq peaks to start slightly downstream of where the detectable signal at the 5'-end of the transcribed region begins, a direct overlap may not exist between the two. Thus, we employed this bedtools closest method for our comparisons to provide a 300 bp buffer region where we can detect peaks from both methods that we expect are arising from the same enhancer.

Using the data available for each cell line in the DPCL collection, we found that the highest proportions of BruUV-peaks overlapped PLS, dELS, or pELS cCREs (Supplementary Fig. 10b), supporting the expectation that BruUV-seq signal peaks are found at the TSSs of genomic elements, like gene promoters and enhancers. Furthermore, ~70% of BruUV-peaks have a corresponding PRO-cap peak, and ~40% of peaks overlap an open chromatin region on average (Supplementary Fig. 10c). This lower proportion of overlapping regions can be partially explained by the stringent FDR thresholds imposed on the ATAC/DNase-seq data used in these comparisons, however an exhaustive exploration of BruUV-peaks without evidence of open chromatin was not performed. When we focused on BruUV-seq enhancer peaks, we found a higher degree of overlap between our peaks and chromatin accessibility regions, with ~80% of our peaks overlapping at least one ATAC-seq peak and ~50-60% overlapping at least one DNase-seq peak (Supplementary Fig. 10c). We also observed a high degree of overlap with PRO-cap peaks at around 75%, and these proportions were slightly increased in the intergenic subset of peaks where we had a higher confidence that the peak signal was the result of enhancer activity and not annotated gene activity (Supplementary Fig. 10c).

Together these data suggested that calling peaks of BruUV-seq signal allowed us to identify relevant regions of high RNA expression and that, using an enhancer annotation (e.g. ENCODE4 dELS cCREs), we can extract information about eRNA transcription from this data that has expected markers of active transcription and correlate with an analogous method. It is important to note that at the current sequencing depth BruUV-seq is not as sensitive as other nascent RNA-seq techniques, such as PRO-cap, which yield more precise 5'-end information and by extension a larger number of discrete peaks as they enrich for capped RNA. An advantage of BruUV-seq over a cap-based method, however, is that we can obtain a broader region of signal at these elements which can provide further insight into the patterns of transcription occurring at these loci.

### 6.2. nTS/nCD validation with predicted ENCODE4 E2G contacts

The results presented in the main text described a pattern in the RNA signal at an enhancer element and the enhancer-promoter contacts that were made in both intact Hi-C and POLR2A ChIA-PET data. To further support the patterns in net target strand (nTS) and net contact direction (nCD) described, we performed these analyses (for full description see Methods: Enhancer-promoter and eTSS-promoter looping analysis) using predicted enhancer-gene contacts (E2G predictions) published by the ENCODE4 consortium[99]. These E2G predictions come from a new predictive model, ENCODE-rE2G (https://github.com/karbalayghareh/ENCODE-E2G), and were available for all cell lines that were included in our enhancer-promoter interaction analysis (ENCODE accessions for these data are found in Supplementary Data File 1). For this analysis we used thresholded E2G predictions that were selected based on a score cutoff that results in 70% recall in the CRISPR benchmarks used to validate these predictions[99].

These analyses were carried out as described for both intact Hi-C and POLR2A ChIA-PET, however the enhancer-gene pairs were already defined, meaning our identified enhancer peaks were simply intersected with E2G predicted enhancers to obtain the final list of enhancer-promoter contacts assessed. Importantly, the same results were obtained using this third method of obtaining relevant enhancer-gene pairs (Supplementary Fig. 11), further supporting our observations that asymmetrical transcription at an enhancer element often arises from looping-interactions with a target gene promoter with a preferred orientation with respect to its strand and linear position. These data are available in Supplementary Data File 4.

#### 6.3. Enhancer consensus locus analysis (nCD)

To compare eRNA between cell lines we obtained consensus regions (bedtools intersect -f 0.90 -F 0.90 -e) and reported these coordinates as the most 5' and 3' coordinates of all overlapping peaks (Supplementary Data File 4). For consensus loci that contained expressed enhancers in 2 or more cell lines, we counted the number of loci where all classifications were the same and where at least one classification was different (excluding low-confidence bidirectional peaks) and found that if we consider all signal classes, we had a roughly even split between loci with the same or different classifications (1434 and 1572, respectively). However, due to the potential for signals to be very similar near the thresholds used for each classification (Extended Data Fig. 5a, Supplementary Data File 4), we focused on loci where multiple cell lines displayed symmetrical or highly asymmetrical classifications (asymmetrical high plus/minus, individual plus/minus). In this subset, most loci had similar eRNA signal (912/1387), agreeing with previously published results[28]. We assessed the nCD indices between cell lines in this subset of consensus loci with symmetrical or highly asymmetrical signal classifications. Because our data for ChIA-PET was shallower (compiled from only 4 cell lines), we focused on intact Hi-C data only for this analysis. We counted the number of consensus loci where all enhancers displayed nCD indices that were in the same net direction (i.e. positive or negative scores) and grouped these according to if their signal classifications matched or at least one class was different (Extended Data Fig. 7c) and found that most nCD scores were similar at consensus loci and this was true for both matched and mismatched signal classifications. We then assessed only consensus loci that had 2 or more cell lines with highly asymmetrical or unidirectional enhancer signal and counted the number of loci where the signal classification corresponded to the expected direction bias in nCD (asymmetrical/individual plus enhancers with a negative nCD, and vice versa) in all or a subset of cell lines (Extended Data Fig. 7d). This analysis revealed that very few consensus loci displayed enhancers with different nCD orientations between cell lines, indicating that at a consensus locus, most enhancers seem to make similar contacts and suggesting that nCD is not predictive of signal but is merely correlated. These results were validated using E2G enhancer-promoter loops (see Supplementary Methods section 6.2, Supplementary Fig. 11g-h).

#### 6.4. nTS/nCD correlation and kernel density estimation

Co-occurrence of nTS and nCD were assessed using linear regression and density estimation in R. Scaled kernel density estimates performed by the ggplot2 plotting function stat_density_2d_filled(n = 10) where the contour_var = 'ndensity' and the fill was implemented by after_stat(level). The lower n parameter (grid points) was used in the main plots to smooth density projection; however, these data were also plotted under the default setting of n = 100 (Supplementary Fig. 12). Linear regression was performed using the geom_smooth(method = 'lm', formula = y~x) function, and R and p-values were displayed on the plots using the stat_cor() function of ggpubr (v0.4.0).

### 7. Readthrough

#### 7.1. Readthrough segment (RTseg) classification

To easily identify RTsegs with particular features or that may reflect technical ambiguity, our RTsegs are tagged with a classification string that describes their overlap with downstream annotated genes (GENCODE v29). RTsegs were classified into 4 main categories: Class I RTsegs do not overlap any annotated genes on the same strand, Class II RTsegs overlap annotated genes that are either small (<1 kb) or not expressed (BruChase-seq 6h RPKM < 0.5), Class III RTsegs overlap downstream genes that are either expressed (BruChase-seq 6h RPKM > 0.5) or had complex, ambiguous annotations, and Class IV RTsegs overlap genes on the opposite strand. Importantly, some RTsegs have patterns of gene overlap that make them more or less suitable for a given analysis. These classifications can help to distinguish the potentially unique properties of each type of RT event and identify segments that may have ambiguity associated with them for easier removal from downstream analyses.

The full breakdown of these classifications with their definitions can be found in Supplementary Table 4. All Class I and II RTsegs were considered to be unambiguous and usable in most analyses. Class I RTsegs in particular could be useful for understanding patterns of RT transcription without the possibility of downstream gene sequence features or protein-binding influencing the patterns of transcription observed. However, it is important to note that class I RTsegs can still overlap genes on the opposite strand (class IV) which could influence RT signal patterns. Class IV RTsegs were generally included in downstream analyses, but RTsegs that overlap TSSs of genes on the opposite strand (Class IVd, IVe, IVf) were considered ambiguous due to the possibility that PROMPT signal may influence the RT signal profile. However, in analyses that were not focused on RT signal patterns, class IV segments were typically retained.

All Class III RTsegs were considered ambiguous and were excluded from downstream analyses involving segment lengths or signal. This was due to the difficulty determining if the downstream RT signal captured is a result of exclusively RT transcription or is coming from new initiation at a downstream gene (Class IIIa) or expression of an overlapping gene (Class IIIb). Class IIIa RTsegs may result from the technical limitations of our method, where RTsegs that overlap expressed genes could result from the inability of the HMM to detect an increase in transcription (new initiation) at the downstream TSS (i.e. no increase in HMM index). Expression in this case was determined by mature mRNA presence in 6h BruChase-seq data due to the possibility of RT transcription from upstream genes to influence the RPKM of the overlapped gene in the nascent data. Class IIIb are likely to represent real examples of RT transcription, but due to overlapping annotations there is uncertainty surrounding the accuracy of their boundary definitions.

An additional step was added to manually evaluate Class III segments for discernable initiation in Bru-seq samples at downstream expressed genes, in an effort to redefine RTseg boundaries and remove ambiguity where possible. To do this, counts and RPKMs of the 1 kb regions flanking the TSS of the overlapping gene were obtained, where the 1 kb upstream of the TSS was considered the possible end of the readthrough for the upstream gene (E), and the 1 kb downstream of the TES represented the gene body signal (G). Similar to our method of selecting TESs, we used the GENCODE v29 annotation to obtain one TSS per gene by selecting the 5'-most start site from all annotated gene isoforms. Signal ratios between the intergenic readthrough region, E, and the gene body, G, were calculated (E/G) and RTseg boundaries were redefined if the E/G ratio was less than 0.5 or if the 1 kb intergenic region, E, had zero counts. In these cases, the 3' coordinate of the original RTseg was set to the TSS of the evaluated gene. A new RTseg was defined for the downstream gene in cases where the previous segment traversed the entirety of the gene body and ended in the intergenic space downstream, where the new 5' coordinate of this segment was defined as the TES of the associated gene. Redefined or newly defined segments from this step are denoted as Class IId (Supplementary Table 4).

It is important to note that one RTseg may overlap multiple genes on the sense or antisense strand, thus RTsegs can have multiple classes listed in their classification string, each describing a different gene(s) that it overlaps (note: a string contains each class only once, thus may not directly indicate the number of genes overlapped). If a particular RTseg had any ambiguous classifications that could influence downstream results, it was removed from our analyses. However, it was assumed in analyses that were not based on the length or signal of an RTseg, that all segments reflected real RT signal downstream of genes, thus RTsegs were included unless other requirements of the analysis were not met, e.g. requiring no downstream genes within 7 kb when assessing sequence and chromatin features (Supplementary Methods, section 7.5-6). Although ambiguous RTsegs (Class IIIa, IIIb, IVd, IVe, IVf) were excluded from analyses involving RT length, it is worth noting that the overall length distribution is shifted only ~2.5 kb shorter (median = 14.9 kb, Supplementary Fig. 13c) than the distribution reported for all segments, thus these segments may not shift our understanding of the typical RTseg length in a way that greatly changes our interpretation of the data overall.

### 7.2. RT gene biotype group definitions

GENCODE biotypes (https://www.gencodegenes.org/pages/biotypes.html) were reorganized into 8 summary groups based on their definitions as follows (group name – biotype list):

| | |
|---|---|
| Protein-coding..... | protein_coding |
| lncRNA………….. | lincRNA, bidirectional_promoter_lncRNA, macro_lncRNA, 3prime_overlapping_ncRNA, non_coding, processed_transcript, sense_intronic, sense_overlapping |
| Antisense……….. | antisense |
| Pseudogene…..... | processed_pseudogene, unprocessed_pseudogene, transcribed_unprocessed_pseudogene, transcribed_processed_pseudogene, transcribed_unitary_pseudogene, unitary_pseudogene, polymorphic_pseudogene, pseudogene, translated_processed_pseudogene |
| IG………………... | IG_C_gene, IG_D_gene, IG_J_gene, IG_LV_gene, IG_V_gene, TR_C_gene, TR_J_gene, TR_V_gene, TR_D_gene, IG_pseudogene, IG_C_pseudogene, IG_J_pseudogene, IG_V_pseudogene, TR_V_pseudogene, TR_J_pseudogene |
| TEC……………... | TEC |
| nc_RNA………… | snRNA, miRNA, snoRNA, rRNA, scaRNA, sRNA, Mt_rRNA, Mt_tRNA, misc_RNA, scRNA, ribozyme, vaultRNA |
| nc_pseudogene... | Mt_tRNA_pseudogene, tRNA_pseudogene, snoRNA_pseudogene, snRNA_pseudogene, scRNA_pseudogene, rRNA_pseudogene, misc_RNA_pseudogene, miRNA_pseudogene |

These groups were used for the purposes of summarizing the identities of all RT genes only (Extended Data Fig. 8a-b, Supplementary Fig. 13a). Biotypes were grouped based on our interpretation of the transcripts that fall under each GENCODE biotype based on the table linked above. Due to the exclusion of genes that are less than 1 kb in length, small non-coding RNA biotypes (nc_RNA and nc_pseudogene) are not present in our analysis. In addition, less than 5 genes in the IG group (Immunoglobulin variable chain and T-cell receptor genes) were found for any sample in our analysis, thus this group is excluded from the plots in Extended Data Fig. 8a-b.

### 7.3. Implications of merging RTsegs for replicate samples

RTsegs were identified for each sample individually but were merged to create a cell line specific annotation of RT regions for most analyses. To merge replicates per cell line, the longest segment from either replicate sample was retained as the RTseg for each gene. This method of merging replicate information relied on the assumption that each replicate would have identical RT potential downstream of a particular gene, which we think is appropriate for the current study. However, it is important to note that the RTseg length distributions per sample were slightly shorter than those found for merged segments (median = 12.1-15.2 kb), with similar length distributions observed between replicates and cell lines (Supplementary Fig. 13e). When we compared the RTseg lengths between replicates, we found the median segment length difference was 3 kb (Supplementary Fig. 13f). However, the range of segment length differences was just as wide as the lengths of the segments themselves, from 0 bp to more than 500 kb. The larger differences observed between replicate RTsegs may be due to the technical limitations of our method, which are described further in Supplementary Methods section 7.8.

Overall, our data for both replicate specific and merged RTseg lengths agree with previously published distributions of RT transcript/DoG lengths[6,100], which have reported medians on the order of 10-20 kb and maximum lengths of hundreds of kilobases downstream of the TES. An important difference between our method, and those used previously to assay RT transcripts/DoGs, is that we do not take a threshold for a minimum RT length, however we observe few genes below the typical DoG length requirement of 5-10 kb.

### 7.4. Downstream gene proximity for RT/NRT genes

We assessed the relative positions of nascent RT genes to their neighboring genes to address findings that stress-induced RT transcripts/DoGs tended to be more proximal to downstream genes[36]. The distance in bp of the nearest downstream gene on either strand to each RT (18244) and NRT (19762) gene TES was obtained with bedtools closest (-iu -d). Downstream genes on both strands were included due to the potential for RT to influence, or be influenced by, genes in either orientation[37]. The overall distribution of distances (log10 bp) was reported, in addition to grouping all genes by distance and reporting the overall fraction of genes within each group (Extended Data Fig. 8e). The distance distribution was also reported for the stringent RT/NRT subset established (239 genes in both subsets, Extended Data Fig. 8e), as well as a protein-coding gene subset (Supplementary Fig. 14b) that helps to remove the potential effects of the varying genomic locations of non-coding RNA species. We note that in the stringent and protein-coding subsets, the preference of RT genes for having more proximal downstream neighbors is not quite as clear. Importantly, this analysis was also performed to assess the proximity of the nearest up- or downstream gene (bedtools closest -iu), where a similar trend in gene proximity was observed (Supplementary Fig. 14a).

A recent study suggested that there is no correlation between the distance of RT or NRT genes to their nearest expressed gene downstream[35]. While this is difficult to assess in our data given the relative infrequency of expressed genes that have no RT signal, our RTseg classifications indicate that the majority of RTsegs identified by our algorithm overlap a downstream gene in some capacity (class II, class III, class IV), with less than 20% that overlap no annotated genes on either strand (class Ia only, Supplementary Fig. 14c). The frequency of RTsegs that are found to overlap at least one downstream gene may support the hypothesis that RT transcription can influence the expression of downstream genes either positively or negatively (e.g. by opening chromatin at downstream genes or repressing gene transcription, respectively[37]).

### 7.5. Assessing the chromatin landscape of RT/NRT genes

To compare the chromatin landscape at the ends and downstream of nascent RT genes to those of homeostatic and stress-induced RT transcripts/DoGs[36], we utilized several ENCODE4 datasets describing histone modifications, CTCF binding, chromatin accessibility, and units of 3D chromatin organization. Peaks of ENCODE4 ChIP-seq signal were obtained for the chromatin binding protein CTCF and the following histone modifications: H3K27me3, H3K27ac, H3K4me1, H3K4me3, H3K36me3, and H3K79me2 (bed). In addition, we used peaks called from DPCL ATAC-seq and DNase-seq data, and loop anchors from DPCL POLR2A ChIA-PET (section 2.3) and intact Hi-C loops (bedpe). Finally, chromatin domain boundaries (TAD boundaries) and A/B compartments (bed) were obtained from DPCL intact Hi-C data (section 2.5). File accession numbers for ChIP-seq data and DPCL data can be found in Supplementary Data File 1. All these data describing various features of the chromatin landscape will be referred to as our chromatin regions-of-interest when discussed together throughout the rest of this section.

For each cell line for which we had each chromatin data type, we counted the frequency that the regions 1 kb upstream (TES-1kb) and 5 kb downstream (TES+5kb) of each gene TES were overlapped by a chromatin region-of-interest using bedtools (intersect for .bed files and pairToBed for bedpe files). As with all RT analyses, genes were required to be greater than 1 kb in length. Genes were grouped based on the presence (ROI) or absence (noROI) of an overlapping chromatin region-of-interest, either in each cell line individually or in any cell line. This left us with a binary grouping describing the patterns of chromatin region-of-interest on a per cell line basis or for all cell lines combined. As before, we split our overall gene list into RT and NRT genes, but removed any gene that had an annotated gene on either strand (sense or antisense) within 7 kb of its TES. This resulted in total RT and NRT subsets of 6187 and 8313, respectively. For cell line specific RT/NRT subsets, we obtained all RT genes found in each cell line and generated a random sample of NRT genes of an equal size using the sample() function in base R.

We then performed a Fisher's exact test to compare our binary groups (RT/NRT and ROI/noROI) for each cell line individually and all cell lines combined. Bar graphs were used to summarize the fraction of all RT and NRT genes that had an overlapping chromatin region-of-interest in any cell line. To summarize cell line specific trends between RT/NRT genes and chromatin regions-of-interest, we performed a tetrachoric correlation[101] (https://john-uebersax.com/stat/tetra.htm) using the tetrachoric() function from the psych package in R. between our binary groups (RT/NRT and ROI/noROI) and reported the correlation coefficient ($r_{tet}$) in a heatmap along with Fisher's exact adjusted p-values as calculated above. This analysis was performed for all RT/NRT genes as described (Supplementary Fig. 15a), as well as for stringent RT/NRT genes (Extended Data Fig. 8f-g). For the stringent RT/NRT subset, a slight modification was applied to the cell line specific data, where instead of examining cell line specific RT/NRT genes, the full list of stringent RT/NRT genes was used in each cell line and compared with cell line specific chromatin regions-of-interest. This was done due to the small number of stringent RT/NRT genes obtained from our data, which were not required to be expressed in all cell lines (see Methods: RT versus NRT gene definitions). In addition, we performed these analyses for protein-coding gene subsets of all and stringent RT/NRT genes to verify our results were not influenced by the imbalance of non-protein-coding biotypes found in the NRT gene set (Supplementary Fig. 15b-c).

In addition to the regions described above (TES-1kb, TES+5kb), we explored the chromatin landscape at the start of the gene, from the TSS to 1 kb into the gene body (TSS+1kb). Similarly to previous studies[36], we found RT genes were enriched for markers of actively transcribing genes (Supplementary Fig. 14d, 15). These markers of active transcription are not unique to RT genes, however, and are expected given that most expressed genes have evidence of RT transcription.

### 7.6. Sequence analysis of RT/NRT genes

To understand if there are sequence motifs that demarcate RT genes, such as higher GC-content[35] or depletion of poly-A sites[36,100] (PASs), we obtained the sequences (GRCh38) in the 1 kb upstream (TES-1kb) and 5 kb downstream (TES+5kb) of gene TESs using bedtools getfasta (-s -bedOut). For sequence analyses, we required that genes had no annotated gene on either strand within 7 kb downstream of its TES, as described in Supplementary Methods section 7.5. Using custom python scripts to parse sequence strings, we counted various motifs in each sequence.

First, we quantified the occurrences of each nucleotide (A, T, C, G) and calculated the GC-content of each region flanking the TES: (G+C)/(G+C+A+T). The distribution of GC-content was then reported for RT versus NRT genes both upstream and downstream of the TES (TES-1kb, TES+5kb). We then counted all possible hexamers in each sequence, using similar methods to those used to study RT transcripts/DoGs previously[35,36]. Sequences 5 kb downstream of the TES were split into 1000-character strings (e.g. 1 kb regions), allowing easier comparison with previous studies (which focus on regions more proximal to the TES), as well as allowing easier comparison to the upstream region (TES-1kb). Each six-string k-mer was counted per region for each gene, and then the frequency of each k-mer was summed for same size subsets of RT and NRT genes. Gene subsets were generated by randomly sampling NRT genes down to the number of RT genes (6187) using pandas DataFrame.sample() weighted by the distribution of biotype counts in the gene list to avoid an imbalance of gene types in the NRT subset which has higher numbers of non-protein-coding genes. Once the total counts of each hexamer were obtained per region for RT and NRT genes, we calculated an enrichment score to compare RT versus NRT k-mer counts as previously described[35]: log2(RT/NRT k-mer counts). To assess the significance of each hexamer that was found to be differentially enriched between RT and NRT genes, we established 1000 equal size subsamples (weighted by biotype as above) of RT/NRT genes (N = 500) and generated a sampling distribution of counts (n = 1000) for each hexamer from which we could calculate a p-value by comparing RT and NRT distributions via t-test using scipy.stats.ttest_ind(). K-mers that had at least 100 counts in both the RT and NRT groups, which had an adjusted p-value < 0.01 and had a log2 enrichment score > 0.56 or < -0.56, were considered interestingly enriched or depleted in RT genes relative to NRT genes (Extended Data Fig. 10e).

Based on this hexamer enrichment analysis, we were also able to distinguish the enrichment/depletion of the two most common canonical PASs, AATAAA and ATTAAA, in RT genes. According to this analysis, a slight but significant depletion of these PAS motifs is found downstream of RT genes. In addition, we calculated the sense versus antisense ratio of the PAS motif AATAAA in the 5 kb region downstream of RT versus NRT genes as described previously[100]. To do this, we simply calculated the occurrence of the string AATAAA on the sense and antisense strands downstream of all genes and calculated a ratio of sense/antisense counts. Similarly to previous reports[100], we found a slight depletion in RT genes (Supplementary

Fig. 14e), although this ratio was not found to correlate strongly with RT length (Extended Data Fig. 10a-b).

### 7.7. Readthrough signal region distinctions

Scaled RT signal was calculated from binned counts from either full length RTsegs or from a fixed 17.3 kb region downstream of RT genes (see Methods: Scaled RT signal calculation). These two methods were used to better align with the two primary analyses performed using RT signal regions: 1.) assessment of RT signal length and 2.) comparisons of RT signal patterns between cell lines. RT signal length can of course not be accurately assessed using a fixed length region; thus, it was necessary to obtain binned counts for the full length RTsegs for the purposes of this analysis. This also served as an additional confirmation that our RTseg definitions were representative of relevant RT signal (see Supplementary Methods, section 7.8). However, due to the likelihood that RTsegs of different lengths were observed for the same gene between cell lines, it would have been challenging to accurately compare RT signal patterns in RTsegs between cell lines. Thus, for these analyses we assayed the scaled counts in a fixed length region that was equal to the median RTseg length (17377 bp).

### 7.8. RT signal versus RTseg lengths

Because we measured RT length in two different ways, one based on segmentation data and the other based on binned counts, we assessed the similarity of the two. First, it is important to note the features and limitations of each method. The method of genome segmentation used to identify RTsegs (see Methods: RT segment identification) was a powerful tool that allowed us to find relevant regions of RT signal in an annotation independent manner and without the need for establishing strict coverage cutoffs. However, the assumptions made within this algorithm, such as the requirement for adjacent segments to display equal or decreasing scores, do introduce some technical variability when compiling RTsegs. It is important to note that any instance of signal variability where coverage over a bin (250 bp) spikes or dips, due to read distribution variability not uncommon to Bru-seq data, could result in the definition of a unique segment with higher or lower HMM scores than those around it. Because of this potential for variability, there are instances of RTsegs that are much shorter than the actual RT signal would suggest, due to a segment whose score did not follow the assumed decreasing trend. Additionally, some segments can extend much longer than one might expect based on the appreciable reads at the tail of the segment, however these segments were determined to have relevant signal in our algorithm (HMM index ≥ 3). For these reasons we thought it was important to validate the lengths of our RTsegs using the signal itself, however that method has its own assumptions and limitations. To determine the length of RT signal, we divided each RTseg into 250 bp bins and calculated the scaled binned counts in the region (see Methods: Scaled RT signal calculation) and then determined the overall length of the RT signal to be where the scaled counts reached zero and remained at or below that threshold for four consecutive bins, e.g. 1 kb (see Methods: Approximate distances traveled by RT signals). The first limitation of this method is that the definition of RT regions is still dependent on the RTsegs called, thus segments that are shorter than the true signal profile will not have their RT signal length better resolved by this analysis. However, this method of calculating an approximate distance (to a resolution of 250 bp due to bin size) based on the counts gives us an alternative perspective on the distance that the most relevant signal may travel. The limitation surrounding this method of assessing distance is due to the assumption that no Bru-seq counts for 1 kb indicates that there is no additional relevant signal. Lack of signal can also arise from mapping artifacts (e.g. regions of low mappability), thus these signal-based length measurements are also subject to technical variability. Nevertheless, we think the use of these two orthogonal methods of measuring the length that RT signal extends downstream of the TES helps support our findings.

We found that the length distributions of RTsegs (Extended Data Fig. 8c, Supplementary Fig. 13c) and RT signals (Extended Data Fig. 9a) were quite similar, although RT signals tended to be shorter, as might be expected given the methods used. For an individual gene per cell line, the median difference between the RTseg and RT signal lengths was ~3.6 kb (Supplementary Fig. 13d). Upon correlating RT signal length to RTseg length (log10), we found the two were highly correlated for all RT signals and signals from class Ia RTsegs only (Spearman's ρ = 0.65-0.76, Extended Data Fig. 10a-b). Additionally, all correlations between RT signal lengths and gene features were also performed using RTsegs and were found to give very similar results (Supplementary Fig. 13g-h). It is worth noting that the correlations seen between RT signal length and the proximity of downstream (DS) genes on either strand or the same strand only (ss) are higher in the class Ia subset of RT genes (Extended Data Fig. 10a-b). This higher correlation is difficult to interpret, as it could be an artifact of our method of defining RT regions. The class Ia group is defined as having no overlap with downstream genes and ending more than 5 kb from the nearest downstream gene, thus inherent to this group is a requirement for a particular distance from a gene, making this correlation to distance difficult to interpret in that subset.

### 7.9. Co-transcriptional splicing index calculation

The co-transcriptional splicing index (SI) was calculated per intron as described previously using the following equation[43]:

$$SI = \frac{exon - exon\,reads}{exon - exon\,reads + \frac{(5'\,exon - intron\,reads\ +\ 3'\,exon - intron\,reads)}{2}}$$

and the median SI for all introns per gene was calculated. Both median SI values and the SI of the last intron only were correlated with RT signal length (see Methods: RT signal length correlations) and RTseg length (see Supplementary Methods, section 7.8). In addition, the distributions of both RT gene SI values were plotted showing that RT genes are not exclusively unspliced (Supplementary Fig. 14f).

### Supplementary Tables

| Cell line | Cell type | Status | Source | Culture Medium Composition |
|---|---|---|---|---|
| IMR90 | Lung fibroblast | Normal | ATCC | MEM + 10% FBS |
| HCT116 | Colorectal carcinoma | Cancer | ATCC | McCoy's 5A + 10% FBS |
| K562 | Chronic myeloid leukemia (CML) lymphoblast | Cancer | ATCC | IMDM + 10% FBS |
| PC-3 | Prostate adenocarcinoma | Cancer | ATCC | F-12K + 10% FBS |
| GM12878 | Lymphoblastoid from B-lymphocyte (EBV transformed) | Normal | Coriell | RPMI + 15% FBS |
| Panc1 | Pancreatic carcinoma | Cancer | ATCC | DMEM + 10% FBS |
| HepG2 | Hepatocellular carcinoma | Cancer | ATCC | MEM + 10% FBS |
| MCF7 | Breast adenocarcinoma | Cancer | ATCC | MEM + 10% FBS |
| A673 | Ewing sarcoma | Cancer | ATCC | DMEM + 10% FBS |
| Caco-2 | Colorectal adenocarcinoma | Cancer | ATCC | MEM + 20% FBS |
| MCF10A | Breast epithelial (fibrocystic disease) | Normal | ATCC | MEGM™ BulletKit™ Lonza (MEBM + supplements, excluding GA-1000***) + 100 ng/ml cholera toxin |
| Calu-3 | Lung adenocarcinoma | Cancer | ATCC | MEM + 10% FBS |
| PC-9 | Non-small cell lung adenocarcinoma | Cancer | Sigma | RPMI + 10% FBS |
| OCI-LY7 | Diffuse large B-cell non-Hodgkin lymphoma | Cancer | DSMZ | IMDM + 10% FBS |
| HUVEC | Primary human umbilical vein endothelial | Normal | Lonza | EGM™ BulletKit™ Lonza (EBM + supplements) |
| HMEC | Primary human mammary epithelial | Normal | Lonza | MEGM™ BulletKit™ Lonza (MEBM + supplements) |

**Supplementary Table 1 | Cell line information.** All cell lines were grown under standard conditions in culture medium with the described compositions in a humidified incubator at 37°C with 5% $CO_2$. Cells were treated and/or collected at approximately 80% confluency and were between 6 and 12 passages at the time of collection. ***Please note that MCF10A were cultured in MEBM with all supplements included in the MEGM™ Mammary Epithelial Cell Growth Medium BulletKit™ except for the gentamicin-amphotericin B mix (GA-1000), and with the addition of 100ng/ml cholera toxin.

| Software/package | Version | Subcommands/functions | Links |
|---|---|---|---|
| Bedtools[74] | v2.28.0 | merge, intersect, closest, subtract, getFasta, pairToBed, makewindows, nuc | https://github.com/arq5x/bedtools2 |
| Deeptools[73] | v3.5.1 | bamCoverage, multiBigwigSummary, computeMatrix, plotProfile, computeMatrixOperations | https://github.com/deeptools/deepTools https://github.com/deeptools/pyBigWig |
| bbTools | v38.46 | bbduk | https://sourceforge.net/projects/bbmap/ |
| Bowtie2[102-104] | v2.3.3 | | https://github.com/BenLangmead/bowtie2 |
| samtools[105] | v1.9 | | https://github.com/samtools/samtools |
| STAR[106] | v2.7.0f | | https://github.com/alexdobin/STAR |
| UCSC kentUtils | v1.04.00 | bedGraphToBigWig, bedSort, bigWigMerge | https://github.com/ucscGenomeBrowser/kent/tree/master |
| Subread[75] | v2.0.6 | featureCounts | https://github.com/ShiLab-Bioinformatics/subread |
| MACS2[32] | v2.2.7.1 | callpeak | https://github.com/macs3-project/MACS |
| sambamba[78] | v0.8.0 | | https://github.com/biod/sambamba |
| R | v4.0.4/ v4.2.1 | | |
| DESeq2[107] | v1.36.0 | plotPCA() | |
| ggplot2 | v3.4.3/ v3.4.4 | | |
| ggpubr | v0.4.0 | stat_compare_means() | |
| pheatmap | v1.0.12 | | |
| psych | v2.3.3 | tetrachoric() | |
| rstatix | v0.7.2 | compare_means() | |
| stats | | cor.test(), p.adjust() | |
| tidyverse | v2.0.0 | | |
| Python | v3.9.16 | | |
| pandas | v1.5.3 | DataFrame.sample() | |
| pyBigWig | v0.3.18 | bw.stats(type="coverage") | |
| scipy | v1.8.1 | scipy.stats.ttest_ind() | |

**Supplementary Table 2 | Software versions.** Versions for software and packages used for the primary analyses performed in this publication (including Bru-seq mapping: Supplementary Methods, section 2.2) are listed along with their citations and code source. Most subcommands or functions used from larger tool suites or packages are noted. R and python packages are note under each software (gray).

| Category number | Category Name | BruUV-seq PROMPT RPKM | BruUV-seq gene RPKM | BruUV-seq TSS ratio enrichment | Bru-seq gene RPKM | Bru-seq TSS ratio enrichment | Combined number of TSSs for all 16 cell lines |
|---|---|---|---|---|---|---|---|
| 1 | bidirectional | + | + | + | + | + | 49459 |
| 2 | bidirectional, BruUV-seq gene signal | + | + | + | - | - | 290 |
| 3 | bidirectional, Bru-seq gene signal | + | - | - | + | + | 1 |
| 4 | High PROMPT, low gene signal | + | - | - | - | - | 536 |
| 5 | negligible signal | - | - | - | - | - | 62789 |
| 6 | low-confidence bidirectional, BruUV-seq gene signal | - | + | + | - | - | 976 |
| 7 | low-confidence bidirectional, Bru-seq gene signal | - | - | - | + | + | 150 |
| 8 | low-confidence bidirectional | - | + | + | + | + | 18598 |
| 9 | ambiguous | This includes all the other combinations, primarily consisting of TSSs with inconsistent genic assay parameter tags, i.e., expression and enrichment, per assay | | | | | 74225 |

**Supplementary Table 3 | Overview of TSS categories.** Complete set of parameters used to classify TSSs based on PROMPT and pre-mRNA signal. For a TSS, the criteria received a "+" tag if it was above the set criteria threshold or a "-" tag if it was equal to or below the threshold. We required the two genic parameters, i.e. the TSS-proximal gene RPKM and the TSS ratio enrichment tags, to be the same per assay.

| Class | Status | Gene overlap | Overlap pattern with downstream genes | Class description | Mean N | RTseg length distribution |
|---|---|---|---|---|---|---|
| Ia | Unambiguous | Same strand | Gene ├>5kb┤ RTseg | RTsegs that do not overlap a gene on the same strand and whose 5'-end is more than 5 kb from the nearest gene | 4475 | |
| Ib | Unambiguous | Same strand | Gene ├<5kb┤ RTseg | RTsegs that do not overlap a gene on the same strand and whose 5' end is less than 5 kb from the nearest gene | 1162 | |
| IIa | Unambiguous | Same strand | ├<1kb┤ Gene RTseg | RTsegs that overlap a gene on the same strand that is less than 1 kb in length | 1396 | |
| IIb | Unambiguous | Same strand | Gene Not Exp RTseg | RTsegs that partially overlap an unexpressed gene on the same strand | 1220 | |
| IIc | Unambiguous | Same strand | Gene Not Exp RTseg | RTsegs that completely overlap an unexpressed gene on the same strand | 1287 | |
| IId | Unambiguous | Same strand | Gene ├<1kb┤ Exp 6h Re-evaluated RTseg RTseg1 RTseg2 | RTsegs that overlap an expressed gene on the same strand, but was reevaluated (see Supplementary Methods, section 7.1) | 220 | |
| IIe | Unambiguous | Same strand | ├<2kb┤ Gene Exp 6h RTseg | RTsegs that overlap an expressed gene on the same strand by less than 2 kb, and are presumed to be a result of annotation artifacts or delayed signal increase | 959 | |
| IIIa | Ambiguous | Same strand | Gene Exp 6h RTseg | RTsegs that overlap an expressed gene on the same strand | 830 | |
| IIIb | Ambiguous | Same strand | Gene Exp 6h + overlapping RTseg1 | RTsegs that overlap an expressed gene on the same strand whose annotation overlaps the assigned gene | 1451 | |
| IVa | Unambiguous | Opposite strand | Gene ├<1kb┤ RTseg | RTsegs that overlap a gene on the opposite strand that is less than 1kb in length | 1588 | |
| IVb | Unambiguous | Opposite strand | Gene Not Exp RTseg | RTsegs that completely overlap an unexpressed gene on the opposite strand | 2884 | |
| IVc | Unambiguous | Opposite strand | Gene ├>2kb┤ Exp 0h RTseg | RTsegs that partially overlap an expressed gene on the opposite strand, and ends more than 2 kb away from the TSS of the reference gene | 1744 | |
| IVd | Ambiguous | Opposite strand | Gene ├<2kb┤ Exp 0h RTseg | RTsegs that partially overlap an expressed gene on the opposite strand, and ends less than 2 kb away from the TSS of the reference gene | 419 | |
| IVe | Ambiguous | Opposite strand | Gene Exp 0h RTseg | RTsegs that completely overlap an expressed gene on the opposite strand | 593 | |
| IVf | Ambiguous | Opposite strand | Gene Exp 0h RTseg | RTsegs that partially overlap the TSS of an expressed gene on the opposite strand | 399 | |
| V | Excluded | Either strand | Gene Exp??? RTseg | RTsegs that overlap a gene on either strand that is missing expression information. This is a rare occurence and these RTsegs were excluded if encountered. | 2 | |

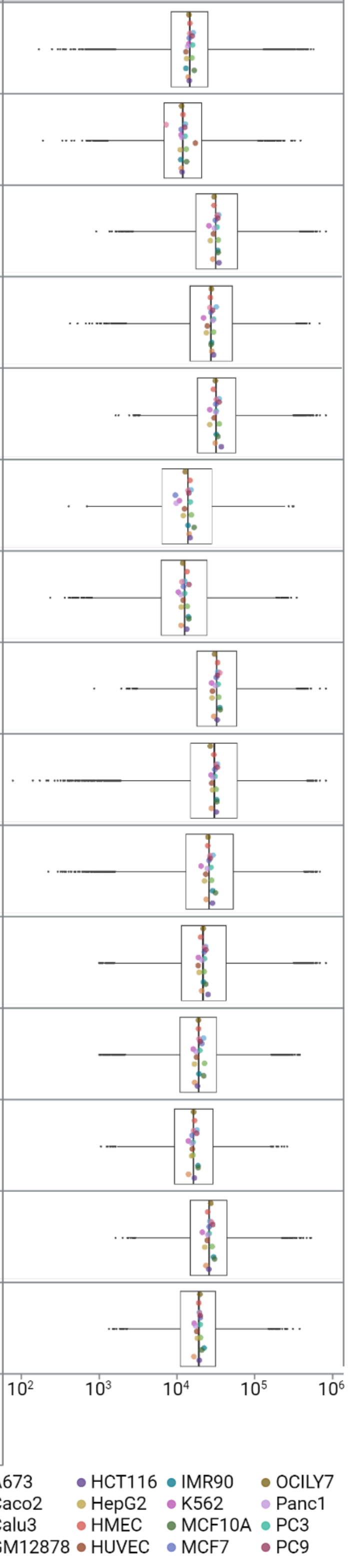

**Supplementary Table 4 | Overview of RT classes.** Each classification string describing possible patterns of downstream gene overlap for RTsegs is defined, where each column provides the following information: Status indicates if the class would be considered ambiguous for measures of RTseg length, Gene overlap denotes if the class describes an overlap with a gene that is on the same strand as the RTseg or the opposite strand, Overlap pattern with downstream genes shows a schematic representation of the overlap pattern for the class, Class description provides a written description of the overlap parameters that distinguish each class that are depicted in the schematic, Mean N gives the average number of RTsegs that have each class across all cell lines, and RTseg length distribution shows the length distribution (log10 bp) of all RTsegs that have each classification (boxplot), where the median length per cell line is denoted by the points on each plot. For more information, see Supplementary Methods section 7.1. “Gene” = RT parent gene, “Exp 0h” = Expressed in Bru-seq data (0h, RPKM > 0.25, used for class IV), “Exp 6h” = Expressed in BruChase-seq data (6h, RPKM > 0.5, used for classes II and III), “Not Exp” = Not expressed (based on expression cutoffs specific to assay used).

### Supplementary Figures

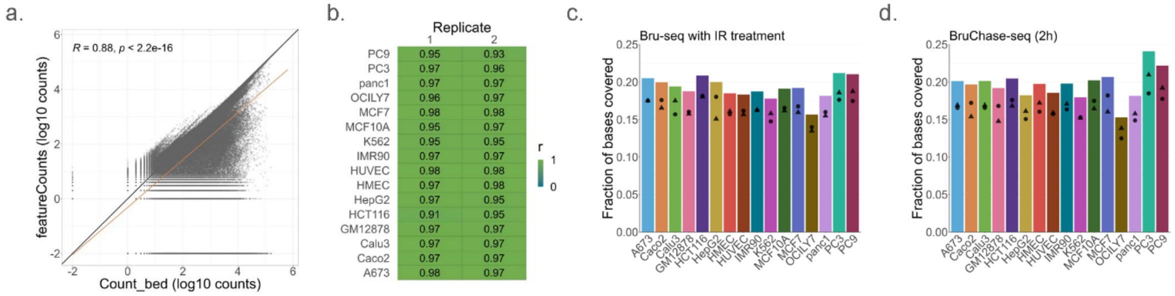

e.

| Cell line | Fraction of intergenic space: Bru-seq | BruUV-seq | PROMPT, eRNA, RT | Fraction of genome: Bru-seq | BruUV-seq | PROMPT, eRNA, RT |
|---|---|---|---|---|---|---|
| All | 0.30095 | 0.30183 | 0.12452 | 0.21531 | 0.21594 | 0.08909 |
| A673 | 0.07901 | 0.07571 | 0.04720 | 0.05653 | 0.05417 | 0.03377 |
| Caco2 | 0.08082 | 0.07916 | 0.04319 | 0.05782 | 0.05664 | 0.03090 |
| Calu3 | 0.07856 | 0.06807 | 0.04553 | 0.05620 | 0.04870 | 0.03257 |
| GM12878 | 0.08806 | 0.09173 | 0.04685 | 0.06300 | 0.06563 | 0.03352 |
| HCT116 | 0.08600 | 0.08027 | 0.04979 | 0.06153 | 0.05743 | 0.03562 |
| HepG2 | 0.06451 | 0.07096 | 0.03980 | 0.04615 | 0.05077 | 0.02847 |
| HMEC | 0.06898 | 0.06860 | 0.04014 | 0.04935 | 0.04908 | 0.02872 |
| HUVEC | 0.06036 | 0.05506 | 0.03625 | 0.04318 | 0.03939 | 0.02593 |
| IMR90 | 0.06339 | 0.05265 | 0.03565 | 0.04535 | 0.03767 | 0.02551 |
| K562 | 0.07399 | 0.07077 | 0.03842 | 0.05294 | 0.05063 | 0.02749 |
| MCF10A | 0.07126 | 0.06716 | 0.04481 | 0.05098 | 0.04805 | 0.03206 |
| MCF7 | 0.08293 | 0.08104 | 0.04240 | 0.05933 | 0.05798 | 0.03034 |
| OCILY7 | 0.06272 | 0.06172 | 0.03848 | 0.04487 | 0.04415 | 0.02753 |
| panc1 | 0.06293 | 0.07564 | 0.04016 | 0.04502 | 0.05411 | 0.02873 |
| PC3 | 0.09049 | 0.08640 | 0.04912 | 0.06474 | 0.06181 | 0.03514 |
| PC9 | 0.09757 | 0.07989 | 0.05039 | 0.06981 | 0.05715 | 0.03605 |

**Supplementary Fig. 1 | Genome coverage of DPCL assays and RNA species. a-b.** Gene counts correlation between the two methods of counting used in this study: Count_bed (see Supplementary Methods, section 2.2) and featureCounts (Methods: Genome coverage calculations). Scatterplot (**a**) shows all data for all cell lines and heatmap (**b**) shows the Pearson's correlation (r) for each cell line. **c-d.** Fraction of bases covered per cell line for DPCL Bru-seq (with ionizing radiation treatment) and BruChase-seq (2h). Bars indicate the cumulative values for each cell line (N = 2), and dots indicate the individual replicate coverages. **e.** Regions of intergenic PROMPT, eRNA, and RT transcription were merged per cell line and for all cell lines to calculate the fraction of bases covered by all identified regions in this study. The fractions of both the intergenic space only (4418874935, based on gencode v29) and the whole genome (6176539664, GRCh38) that are covered by the 3 transcription-associate lincRNAs identified are shown, along with the proportion of intergenic bases covered by all Bru-seq and BruUV-seq reads (as calculated in Fig. 1).

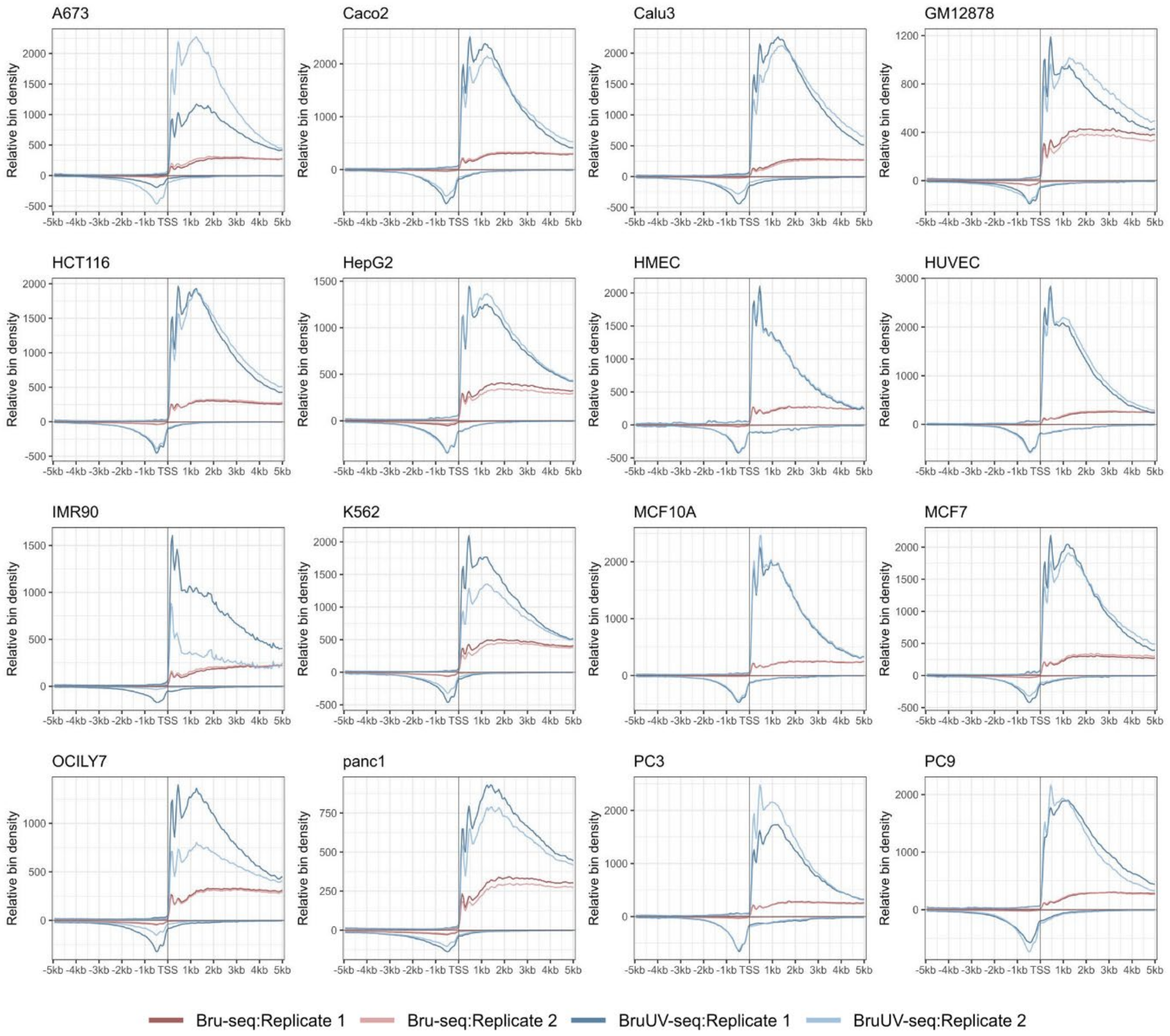

**Supplementary Fig. 2 | Metagene plots of Bru-seq and BruUV-seq data for 16 cell lines.** Metagene plots for 16 cell lines and their replicates comparing Bru-seq and BruUV-seq data profiles for annotated protein-coding gene TSSs (GENCODE v29 annotation, $N_{TSS}$ = 5615).

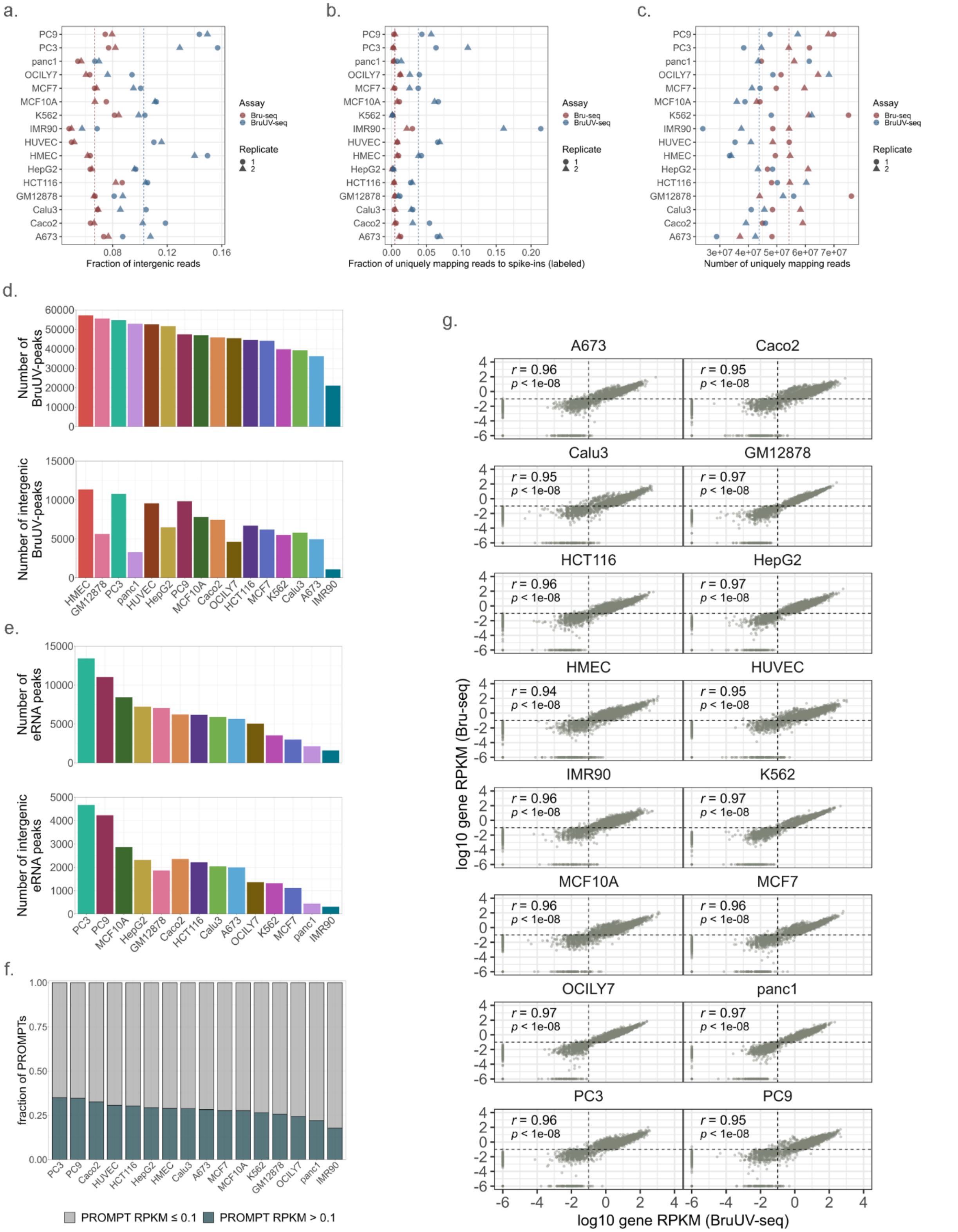

**Supplementary Fig. 3 | Comparisons of Bru-seq and BruUV-seq data. a.** Proportion of uniquely mapping reads mapping to intergenic regions per cell line replicate in Bru-seq and BruUV-seq data. **b.** Proportion of uniquely mapping reads mapping to labeled spike-ins per cell line replicate in Bru-seq and BruUV-seq data. **c.** Proportion of uniquely mapping reads per cell line replicate in Bru-seq and BruUV-seq data. **d-e.** Numbers of BruUV-peaks (**d**) and enhancer peaks (**e**) called in our analysis for each cell line. Peak numbers are shown for all peaks (top) and intergenic peaks only (bottom). **f.** Fraction of PROMPTs (12939) per RPKM criteria shown for each cell line. **g.** Scatterplots showing the relationship between TSS-proximal BruUV-seq and Bru-seq gene expression (log10 RPKM, 2 kb regions) for all cell lines. Pearson's correlations (r) and  associated p-values are displayed for each cell line.

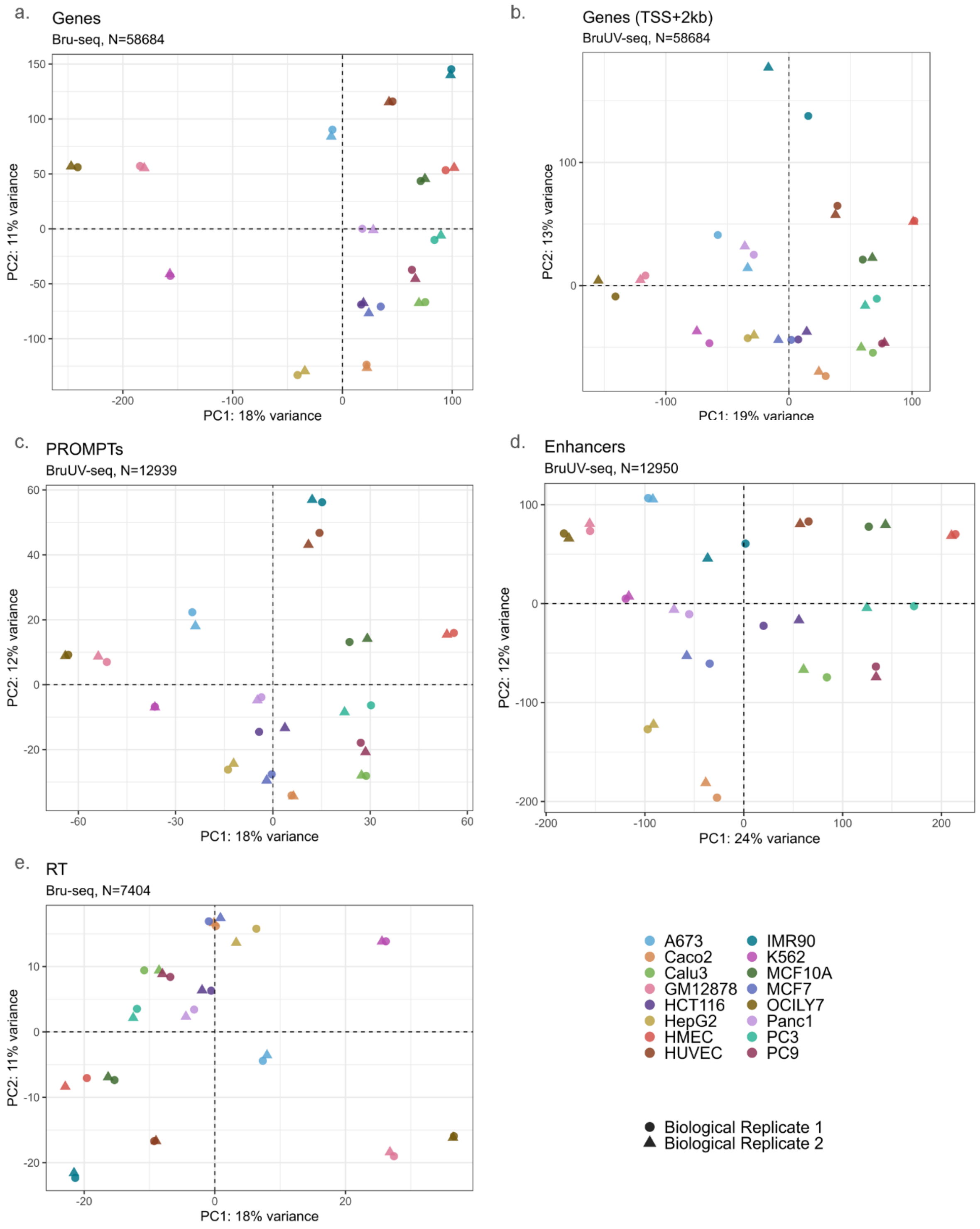

**Supplementary Fig. 4 | Principal component analysis for regions of interest.** Scatterplots showing the principal components with the highest signal variance, PC1 and PC2, between all cell line replicates for the following regions of interest: gene body (**a**, Bru-seq counts), TSS-proximal gene signal (**b**, BruUV-seq counts, TSS+2kb), PROMPTs (**c**), intergenic enhancer consensus loci (**d**), readthrough (RT) transcription (**e**, TES+5kb).

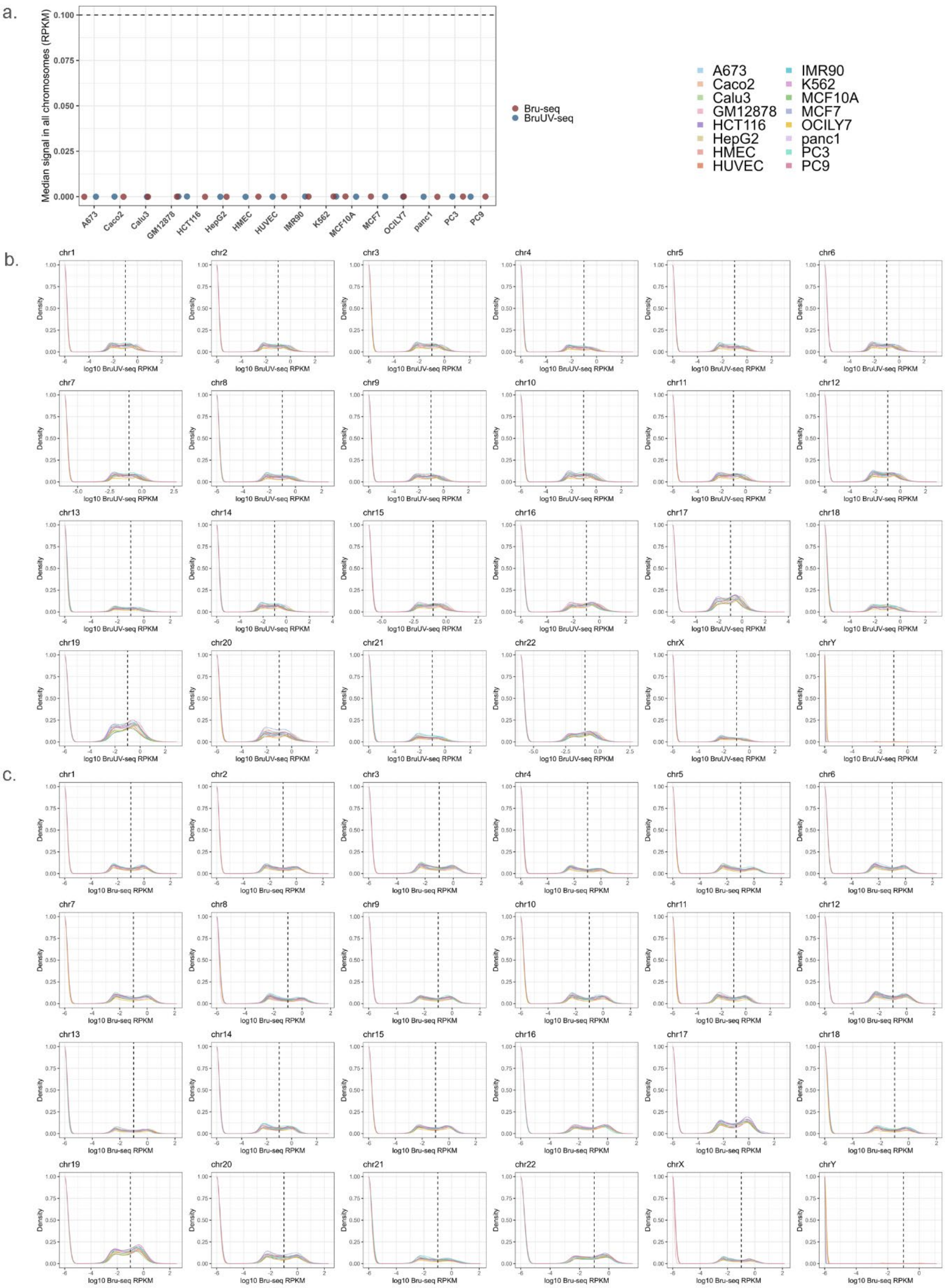
a.
Median signal in all chromosomes (RPKM)
0.100
0.075
0.050
0.025
0.000
A673
Caco2
Calu3
GM12878
HCT116
HepG2
HMEC
HUVEC
IMR90
K562
MCF10A
MCF7
OCILY7
panc1
PC3
PC9
Bru-seq
BruUV-seq
A673
Caco2
Calu3
GM12878
HCT116
HepG2
HMEC
HUVEC
IMR90
K562
MCF10A
MCF7
OCILY7
panc1
PC3
PC9
b.
chr1
chr2
chr3
chr4
chr5
chr6
chr7
chr8
chr9
chr10
chr11
chr12
chr13
chr14
chr15
chr16
chr17
chr18
chr19
chr20
chr21
chr22
chrX
chrY
Density
log10 BruUV-seq RPKM
c.
chr1
chr2
chr3
chr4
chr5
chr6
chr7
chr8
chr9
chr10
chr11
chr12
chr13
chr14
chr15
chr16
chr17
chr18
chr19
chr20
chr21
chr22
chrX
chrY
Density
log10 Bru-seq RPKM

**Supplementary Fig. 5 | Signal profiles in 2 kb genome bins. a.** Median expression value (RPKM) for 2 kb bins across all the standard chromosomes (chr1-22, X and Y) calculated per strand for all cell lines. **b-c.** Density plots of RPKM values for 2 kb bins per chromosome for BruUV-seq data (**b**) and Bru-seq data (**c**).

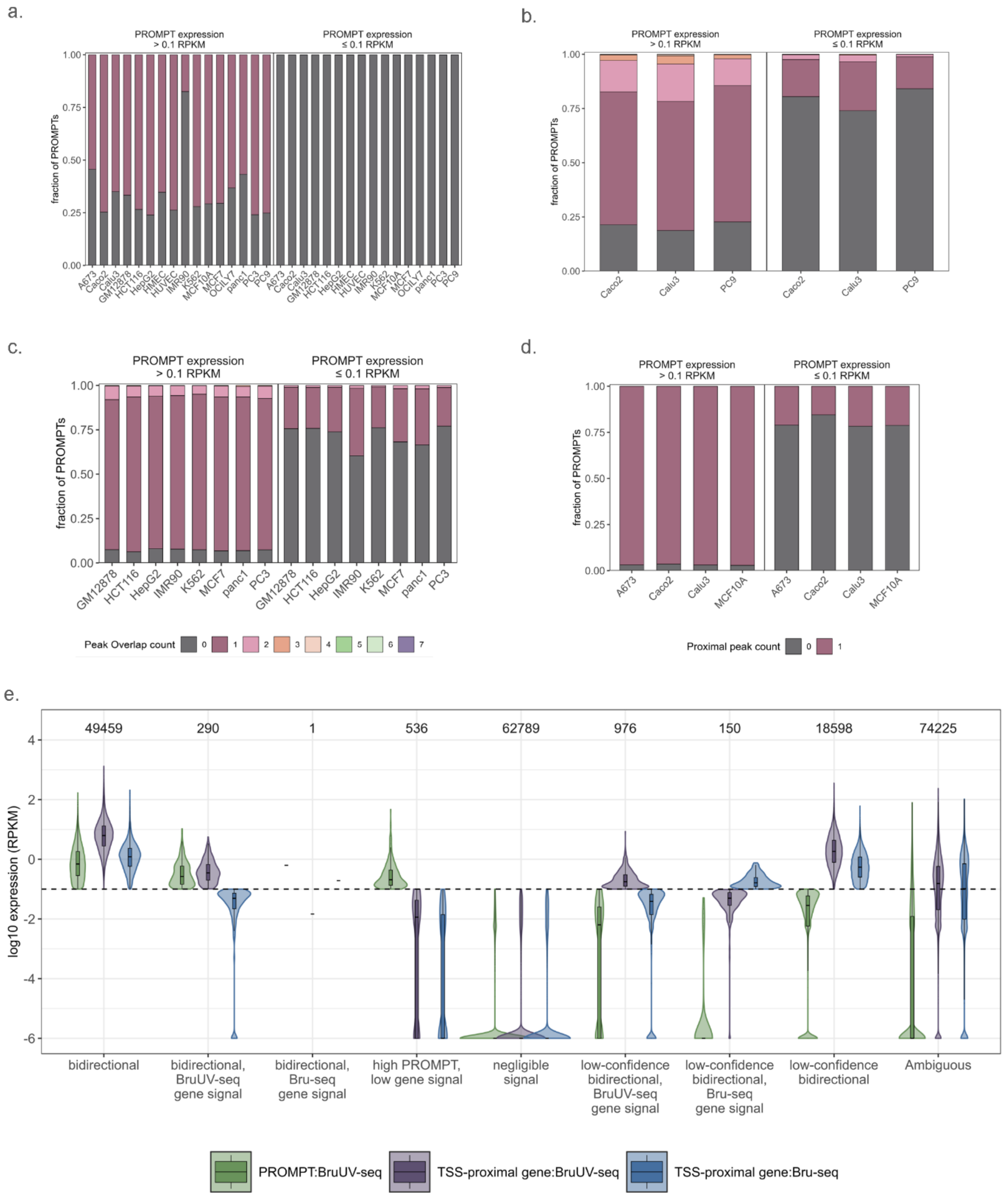

**Supplementary Fig. 6 | Signal profiles of TSS-proximal regions and association of PROMPT regions with supplemental and orthogonal data types. a-d.** Fraction of PROMPT regions with BruUV-seq peak overlaps (**a**), DNase-seq peak overlaps (**b**), ATAC-seq peak overlaps (**c**), or with PRO-cap peaks within 1 kb upstream (**d**). Each plot is faceted based on PROMPT expression representing high-confidence (BruUV-seq RPKM > 0.1) or low-confidence signal (BruUV-seq RPKM ≤0.1). **e.** Distributions of BruUV-seq and Bru-seq RPKMs for the PROMPT and TSS-proximal 2 kb regions for TSS categories. Zero RPKM values are represented by a log10 value of -6. The total number of observations per category is indicated above each set of distributions.

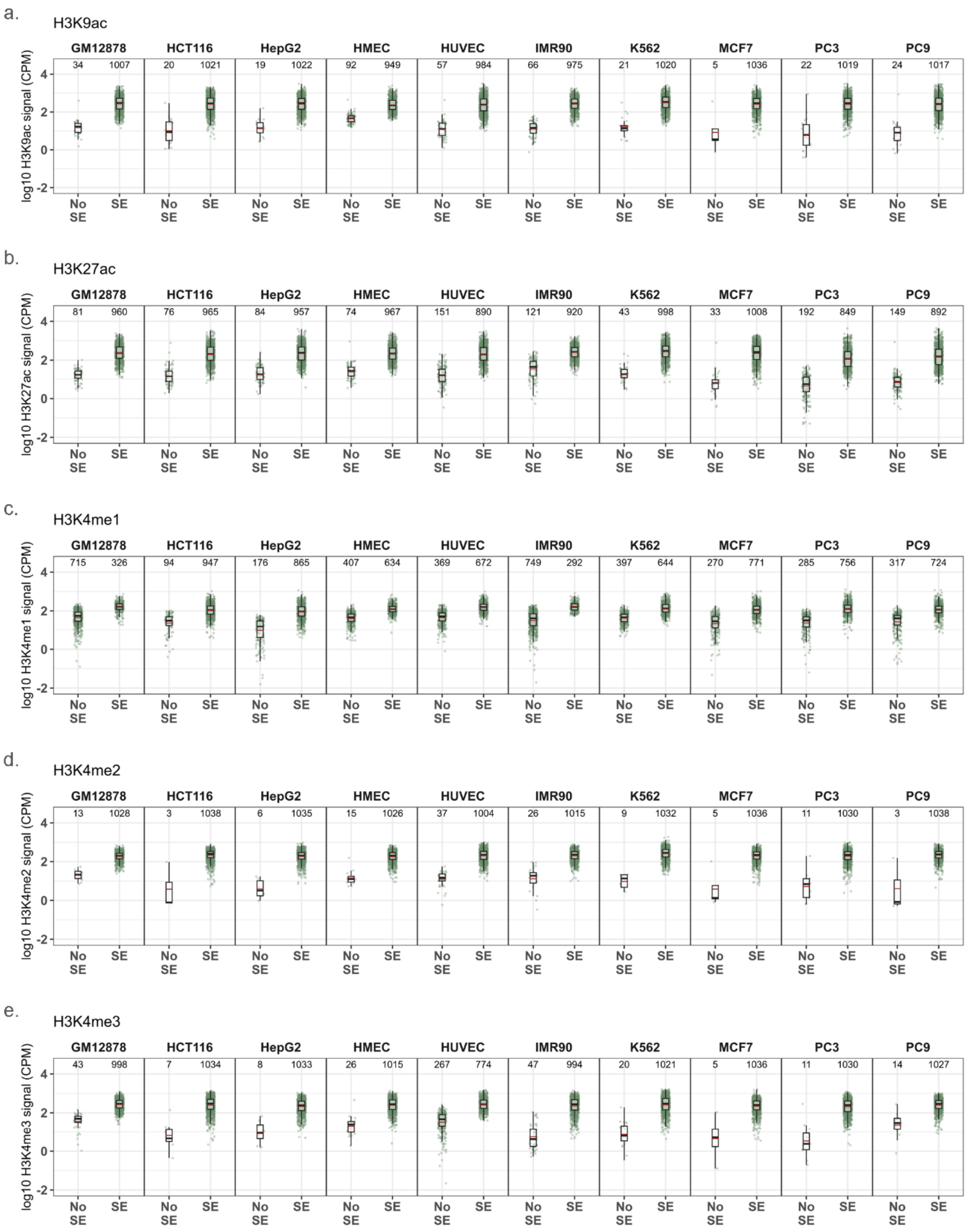

**Supplementary Fig. 7 | Agreement between histone signal and peaks.** Distributions of histone modification signal (log10 CPM) and regions of detected signal enrichment (determined by peak overlaps) for H3K9ac (**a**), H3K27ac (**b**), H3K4me1 (**c**), H3K4me2 (**d**), and H3K4me3 (**e**) for bidirectional TSSs across all cell lines (1041/12939). SE = Signal Enrichment.

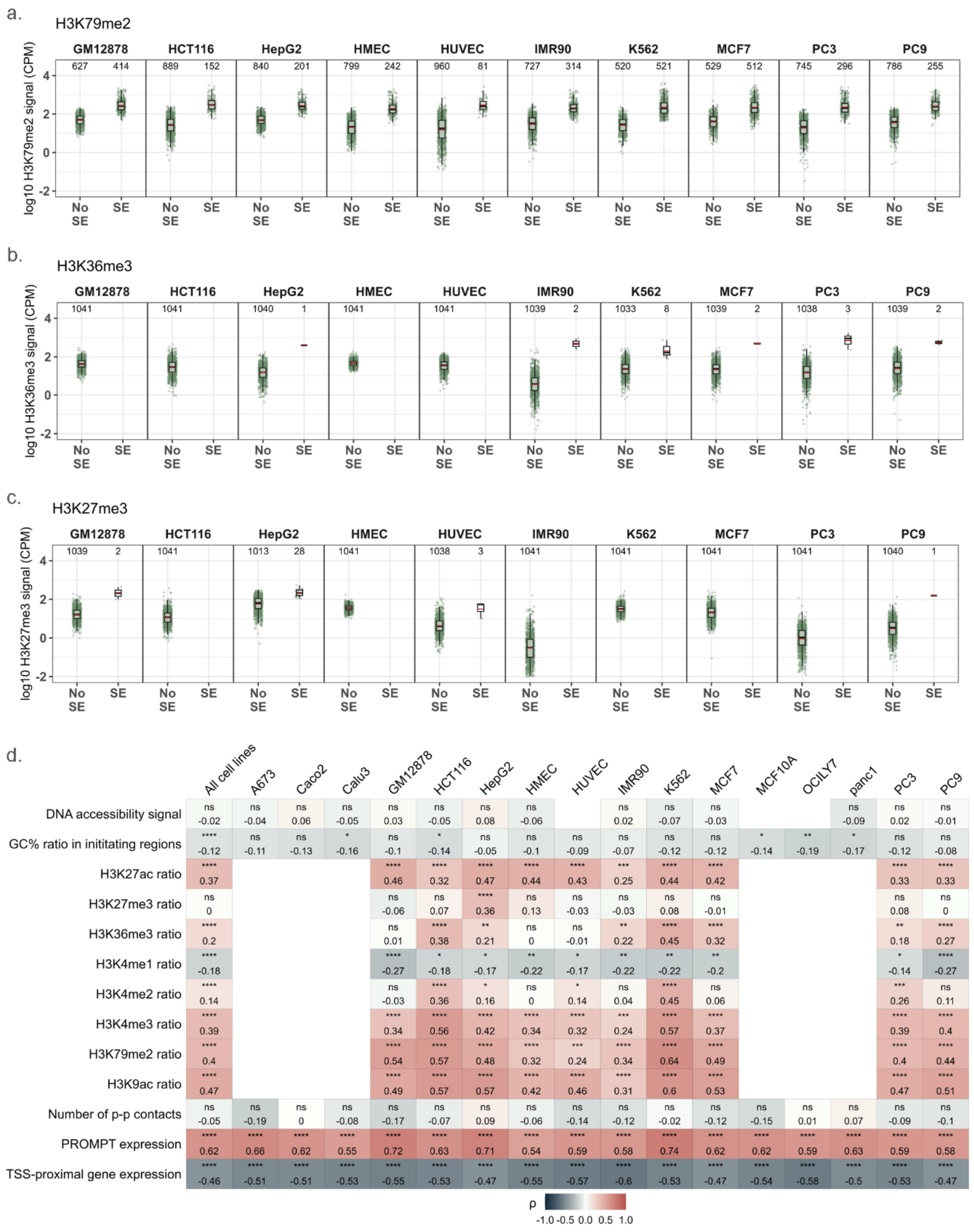

**Supplementary Fig. 8 | Summary of expression, sequence and chromatin attributes for gene and PROMPT regions. a-c.** Distribution of histone modification signal (log10 CPM) and regions of detected signal enrichment (determined by peak overlaps) for H3K79me2 (**a**), H3K36me3 (**b**), and H3K27me3 (**c**) for bidirectional TSSs across all cell lines (1041/12939). SE = Signal Enrichment. **d.** Heatmap of Spearman correlation coefficients (ρ) between various genomic and chromatin features and expression (RPKM) ratios over PROMPT and gene regions flanking the TSS for a subset of single-isoform bidirectional TSSs (234/1041). False discovery rate (FDR) adjusted p-values are indicated as: $^{ns}p_{adj} > 0.05$, $^{*}p_{adj} < 0.05$, $^{**}p_{adj} < 0.01$, $^{***}p_{adj} < 0.001$, $^{****}p_{adj} < 0.0001$.

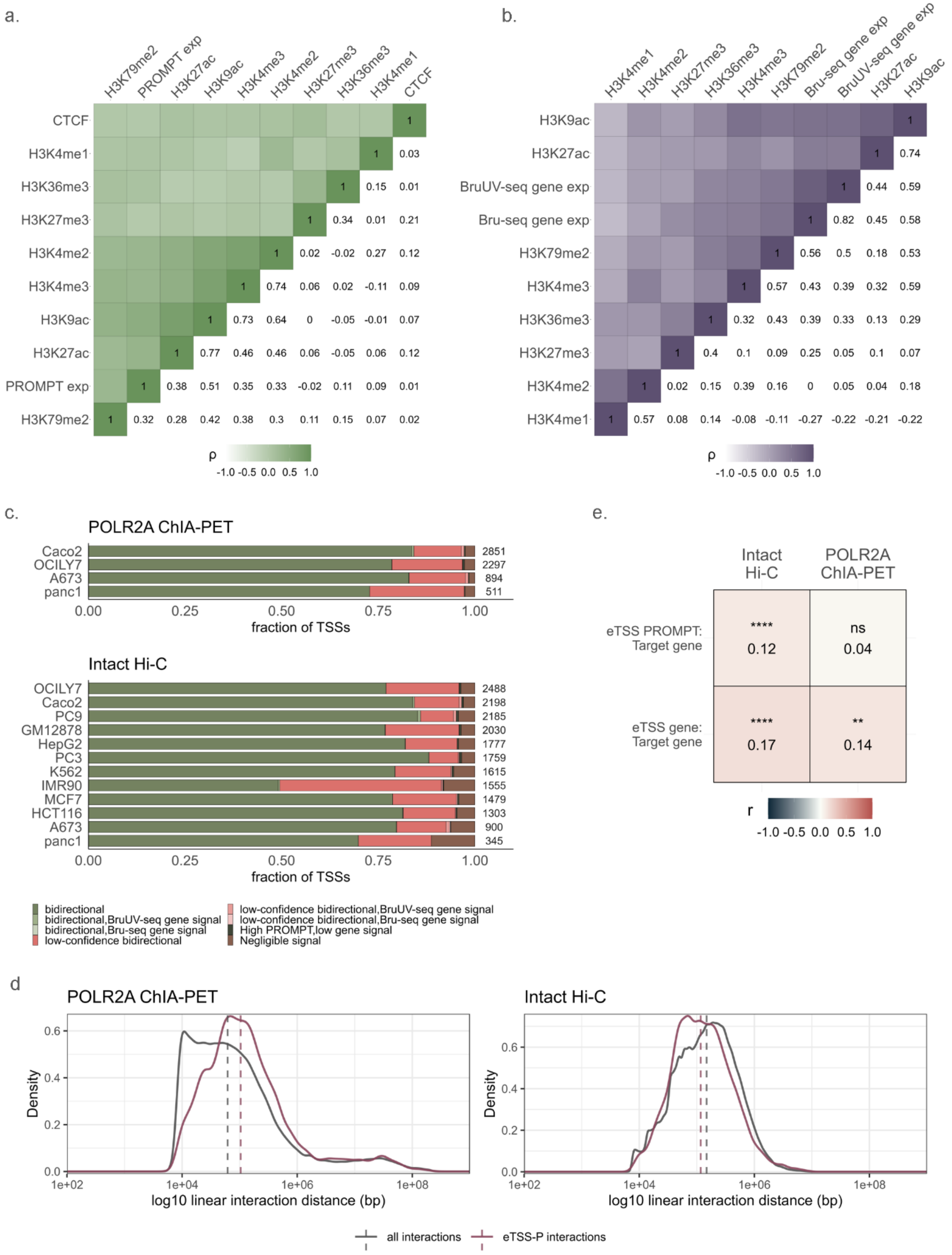

**Supplementary Fig. 9 | Correlations of chromatin features with expression and properties of enhancer-like gene TSSs (eTSS). a-b.** Heatmap of Spearman correlation coefficients (ρ) between various histone modifications, CTCF signal (log10 CPM) and expression (log10 RPKM) values over PROMPT (**a**) and TSS-proximal gene regions (**b**) for bidirectional TSSs (1041/12939). **c.** Distribution of categories per cell line for those TSSs engaging in promoter-promoter distal interactions identified in POLR2A ChIA-PET or intact Hi-C data. **d.** Density plot of interaction distances for eTSSs compared to all interactions. **e.** Heatmap of Pearson's correlation coefficients (r) between an eTSS's PROMPT or gene expression with its target gene's expression. Only eTSSs that were found to make contacts with a single target gene, as well as those single target genes that do not contact any other eTSSs were considered for this analysis (one eTSS-one target gene interactions).

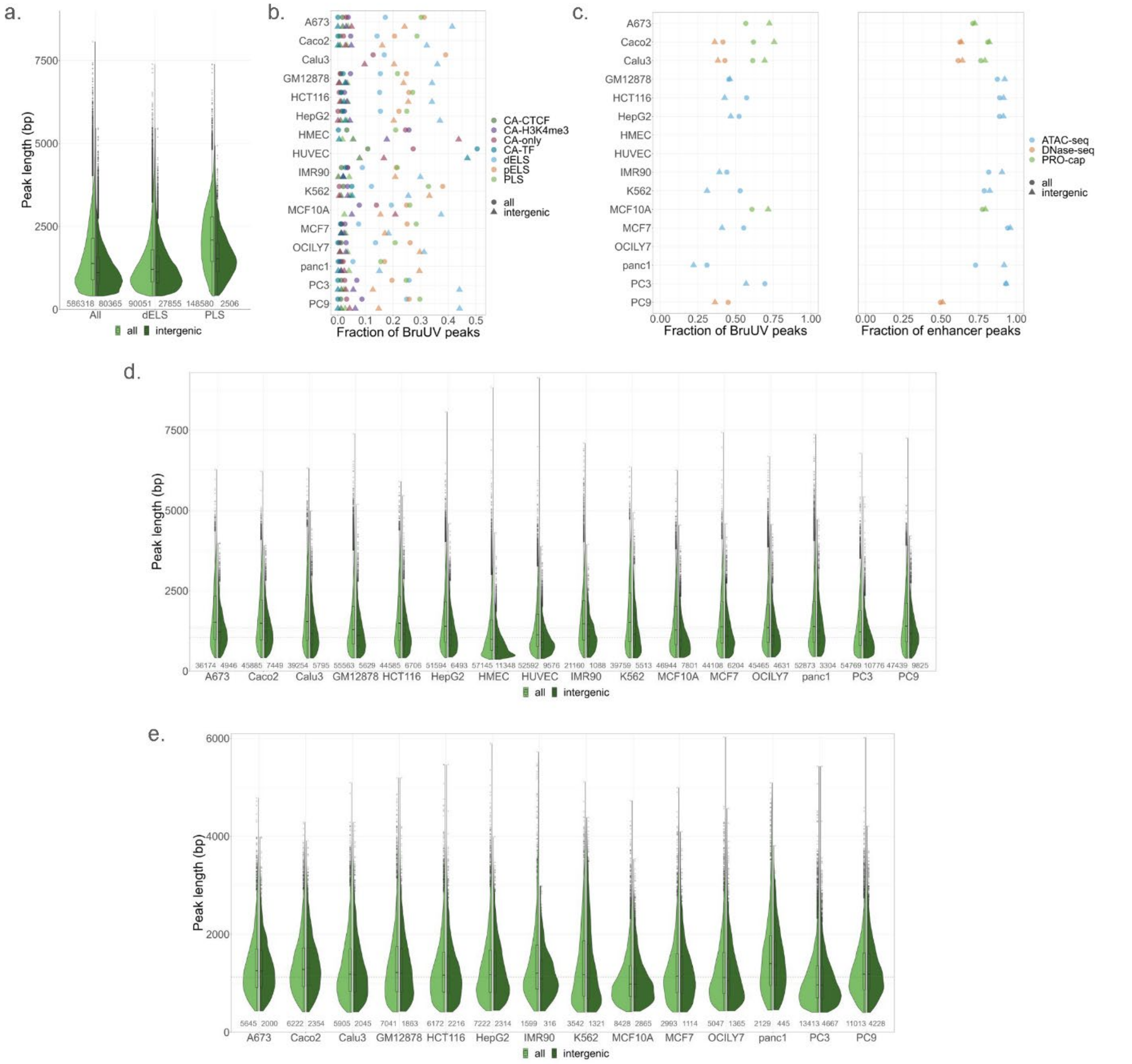

**Supplementary Fig. 10 | Overview of BruUV-peak features. a.** Distributions of BruUV-seq peak lengths for all peaks, peaks overlapping distal enhancer-like cCREs (dELS), and promoter-like cCREs (PLS) in 13 cell lines for which we have these cCRE data. Split violin shows distributions for all peaks (left) and intergenic peaks only (right), and the number below each half indicates the number of peaks in each group. **b.** Fractions of BruUV-peaks that overlap each of the ENCODE4 cCRE regions. Any missing cCRE type per cell line indicates that no data was available for that cCRE type in that cell line. Proportions are shown for all peaks, and intergenic peaks only. **c.** Fractions of BruUV-peaks (left) and enhancer peaks (right) that overlap peaks called in ATAC-seq, DNase-seq, and PRO-cap per cell line. Proportions are shown for all peaks, and intergenic peaks only. **d-e.** Cell line distributions of BruUV-peak (**d**) and enhancer (**e**) peak lengths. Split violin shows distributions for all peaks (left) and intergenic peaks only (right), and the number below each half indicates the number of peaks in each subset.

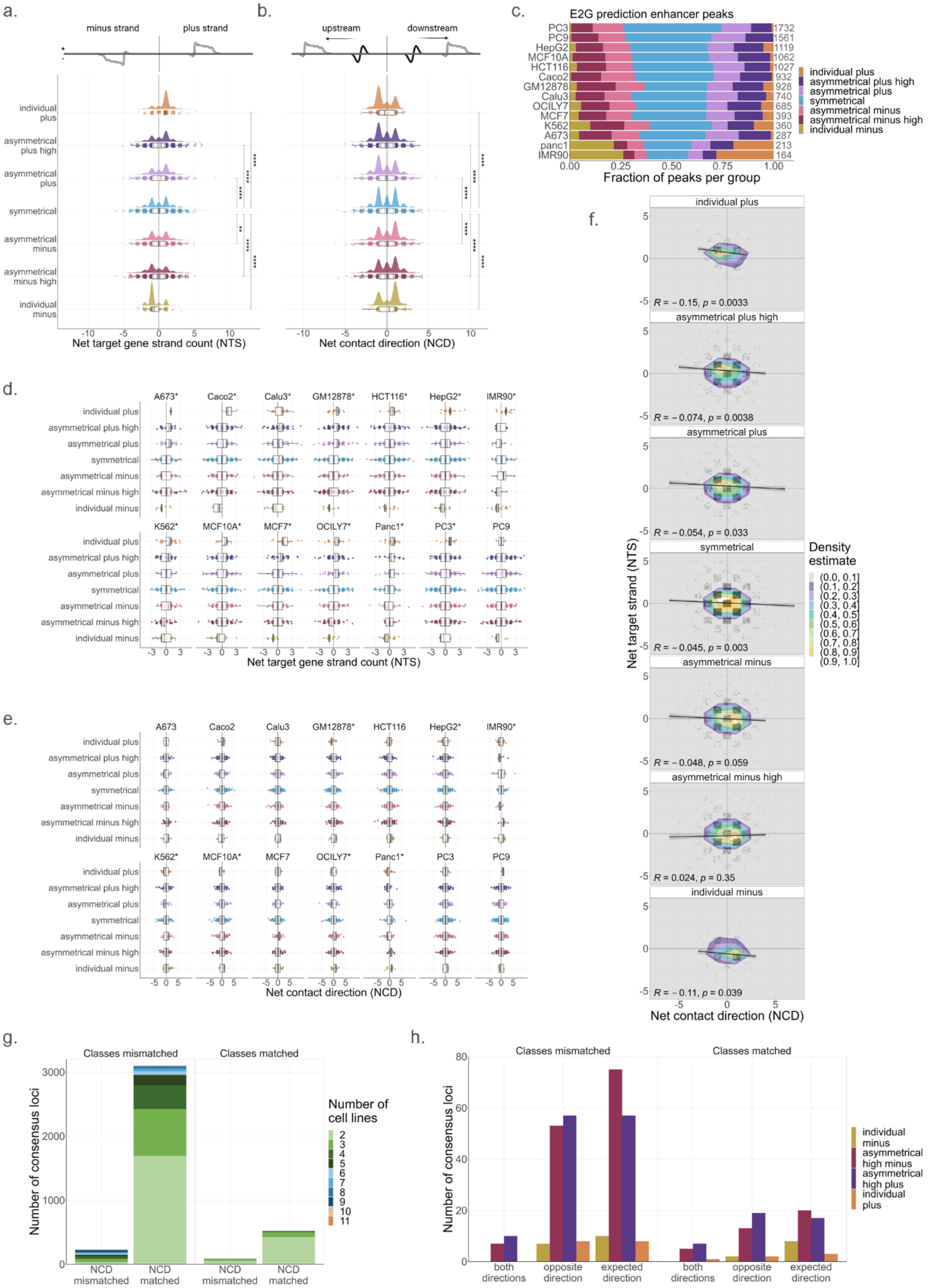

a.
minus strand
plus strand
individual plus
asymmetrical plus high
asymmetrical plus
symmetrical
asymmetrical minus
asymmetrical minus high
individual minus
Net target gene strand count (NTS)
b.
upstream
downstream
Net contact direction (NCD)
c.
E2G prediction enhancer peaks
PC3 1732
PC9 1561
HepG2 1119
MCF10A 1062
HCT116 1027
Caco2 932
GM12878 928
Calu3 740
OCILY7 685
MCF7 393
K562 360
A673 287
panc1 213
IMR90 164
Fraction of peaks per group
individual plus
asymmetrical plus high
asymmetrical plus
symmetrical
asymmetrical minus
asymmetrical minus high
individual minus
d.
A673*
Caco2*
Calu3*
GM12878*
HCT116*
HepG2*
IMR90*
K562*
MCF10A*
MCF7*
OCILY7*
Panc1*
PC3*
PC9
Net target gene strand count (NTS)
e.
A673
Caco2
Calu3
GM12878*
HCT116
HepG2*
IMR90*
K562*
MCF10A*
MCF7
OCILY7*
Panc1*
PC3
PC9
Net contact direction (NCD)
f.
individual plus
R = −0.15, p = 0.0033
asymmetrical plus high
R = −0.074, p = 0.0038
asymmetrical plus
R = −0.054, p = 0.033
symmetrical
R = −0.045, p = 0.003
asymmetrical minus
R = −0.048, p = 0.059
asymmetrical minus high
R = 0.024, p = 0.35
individual minus
R = −0.11, p = 0.039
Net target strand (NTS)
Net contact direction (NCD)
Density estimate
(0.0, 0.1]
(0.1, 0.2]
(0.2, 0.3]
(0.3, 0.4]
(0.4, 0.5]
(0.5, 0.6]
(0.6, 0.7]
(0.7, 0.8]
(0.8, 0.9]
(0.9, 1.0]
g.
Classes mismatched
Classes matched
Number of consensus loci
NCD mismatched
NCD matched
Number of cell lines
h.
Classes mismatched
Classes matched
Number of consensus loci
both directions
opposite direction
expected direction
individual minus
asymmetrical high minus
asymmetrical high plus
individual plus

**Supplementary Fig. 11 | nTS and nCD validation with E2G prediction data. a-b.** nTS and nCD distributions calculated for enhancers interacting with expressed genes via E2G predicted loops. Schematics represent the interpretation of positive and negative nTS and nCD scores (top), where the target gene (gray) is oriented relative to the DNA strand or about the enhancer (black). For gene schematics, the sharp peak represents the PROMPT, and the broad peak represents the gene. FDR adjusted p-values were determined from Wilcoxon signed-rank tests (*$p_{adj}$ < 0.05, **$p_{adj}$ < 0.01, ***$p_{adj}$ < 0.001, ****$p_{adj}$ < 0.0001). **c.** Peak classifications per cell line for enhancers interacting with expressed genes via E2G predicted loops. **d-e.** Cell line distributions of nTS (**d**) and nCD (**e**) for enhancers interacting distally with expressed genes via E2G predicted loops. **f.** Scaled kernel density estimations for nTS and nCD distributions of grouped enhancers. Scaling is performed within groups to account for sample size imbalances. **g.** Quantification of active enhancers that are found in at least 2 cell lines (at consensus loci, see Supplementary Methods, section 6.3), and are interacting with expressed genes via intact Hi-C loops, with nCD scores that are in the same (matched) or opposite (mismatched) net directions. Consensus loci are distinguished by the classifications of the enhancers in each cell line represented, where either all classes match or at least one cell line has a different signal classification (mismatched). This shows that at most consensus loci a particular enhancer tends to make the same net contacts in all cell lines, even if the signal pattern changes between cell lines. **h.** Quantification of enhancers found in at least 2 cell lines, where multiple cell lines have enhancers that are highly asymmetrical or unidirectional, with nCD scores that follow the expected contact direction pattern (Fig. 3d), show the opposite pattern, or show different patterns between cell lines. Consensus loci are again distinguished by matched or mismatched enhancer classifications between cell lines. Since enhancers at these loci are nearly equally likely to match the expected relationship between eRNA signal and nCD or not, with very few instances of mixed patterns, irrespective of any differences in their signal patterns, we conclude that nCD does not dictate changes in enhancer expression.

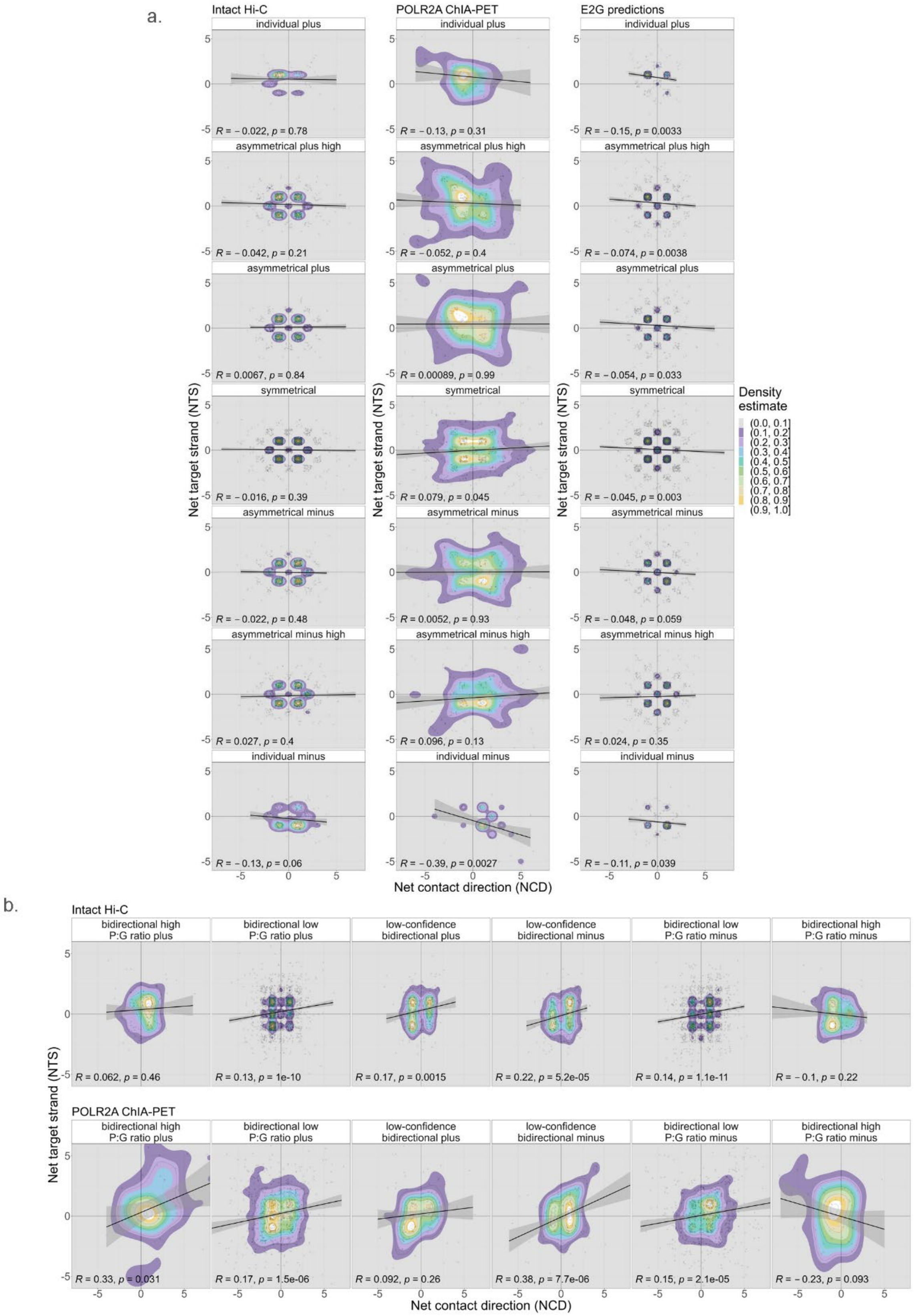
a.
Intact Hi-C
POLR2A ChIA-PET
E2G predictions
individual plus
asymmetrical plus high
asymmetrical plus
symmetrical
asymmetrical minus
asymmetrical minus high
individual minus
R = − 0.022, p = 0.78
R = − 0.13, p = 0.31
R = − 0.15, p = 0.0033
R = − 0.042, p = 0.21
R = − 0.052, p = 0.4
R = − 0.074, p = 0.0038
R = 0.0067, p = 0.84
R = 0.00089, p = 0.99
R = − 0.054, p = 0.033
R = − 0.016, p = 0.39
R = 0.079, p = 0.045
R = − 0.045, p = 0.003
R = − 0.022, p = 0.48
R = 0.0052, p = 0.93
R = − 0.048, p = 0.059
R = 0.027, p = 0.4
R = 0.096, p = 0.13
R = 0.024, p = 0.35
R = − 0.13, p = 0.06
R = − 0.39, p = 0.0027
R = − 0.11, p = 0.039
Net target strand (NTS)
Net contact direction (NCD)
Density estimate
(0.0, 0.1]
(0.1, 0.2]
(0.2, 0.3]
(0.3, 0.4]
(0.4, 0.5]
(0.5, 0.6]
(0.6, 0.7]
(0.7, 0.8]
(0.8, 0.9]
(0.9, 1.0]
b.
Intact Hi-C
bidirectional high P:G ratio plus
bidirectional low P:G ratio plus
low-confidence bidirectional plus
low-confidence bidirectional minus
bidirectional low P:G ratio minus
bidirectional high P:G ratio minus
R = 0.062, p = 0.46
R = 0.13, p = 1e-10
R = 0.17, p = 0.0015
R = 0.22, p = 5.2e-05
R = 0.14, p = 1.1e-11
R = − 0.1, p = 0.22
POLR2A ChIA-PET
R = 0.33, p = 0.031
R = 0.17, p = 1.5e-06
R = 0.092, p = 0.26
R = 0.38, p = 7.7e-06
R = 0.15, p = 2.1e-05
R = − 0.23, p = 0.093
Net target strand (NTS)
Net contact direction (NCD)

**Supplementary Fig. 12 | Higher resolution kernel density estimations.** Scaled kernel density estimations with the default number of grid points (bins) in each direction: stat_density_2d(n=100). This is shown because n was reduced to 10 in main plots to smooth density projection. Data is shown for nTS and nCD distributions of grouped enhancers (**a**) and eTSSs (**b**) for all looping assays. Scaling is performed within groups to account for sample size imbalances.

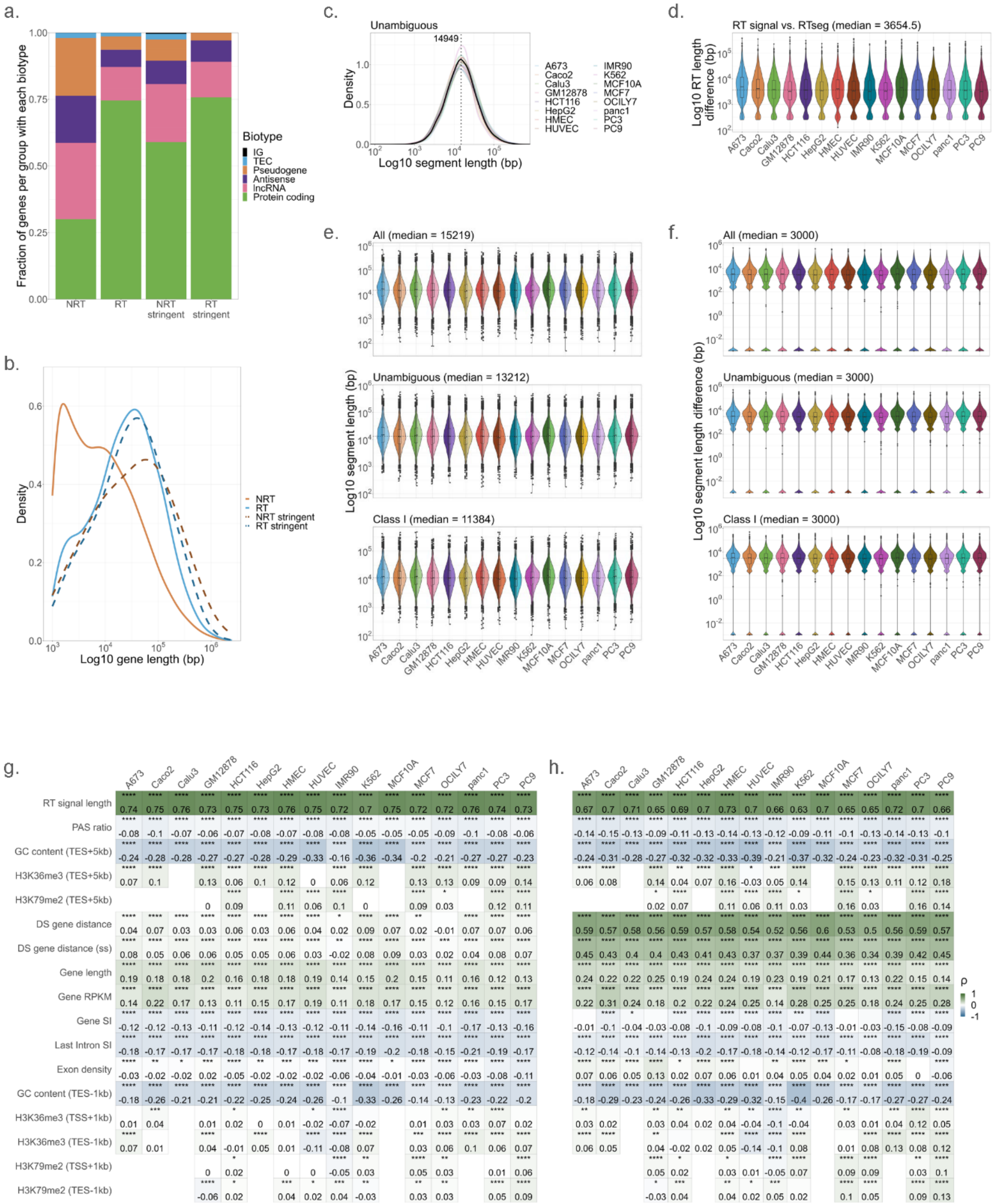

**Supplementary Fig. 13 | RT and NRT gene distinctions and RTseg lengths. a.** The fraction of all and stringent RT/NRT genes that fall into each biotype group. **b.** Log10 distribution of gene lengths (bp) in each of our RT/NRT gene subsets. **c.** Log10 RTseg length distributions for unambiguous RTsegs. Median length is shown (black), along with the cell line specific distributions. **d.** Log10 distribution of RT length differences between RTsegs and RT signal per gene. The median length difference across all cell lines is noted. **e.** Split violin plots depicting the distributions of log10 RTseg length (bp) for replicates 1 and 2 (left and right, respectively) of each cell line. Length distributions are shown for all, unambiguous, and class I RTsegs and the median length for all samples is noted. **f.** Log10 distribution of RTseg length differences (bp) between replicate samples. The median length difference across all cell lines is noted. **g-h.** Correlations between RTseg length and gene sequence or chromatin features for all RT signal regions (**g**) and RT regions that end

> 5 kb away from annotated genes (**h**, RTseg class Ia). Spearman's rank correlation coefficients (ρ) and FDR adjusted p-values (*$p_{adj}$ < 0.05, **$p_{adj}$ < 0.01, ***$p_{adj}$ < 0.001, ****$p_{adj}$ < 0.0001) are displayed.

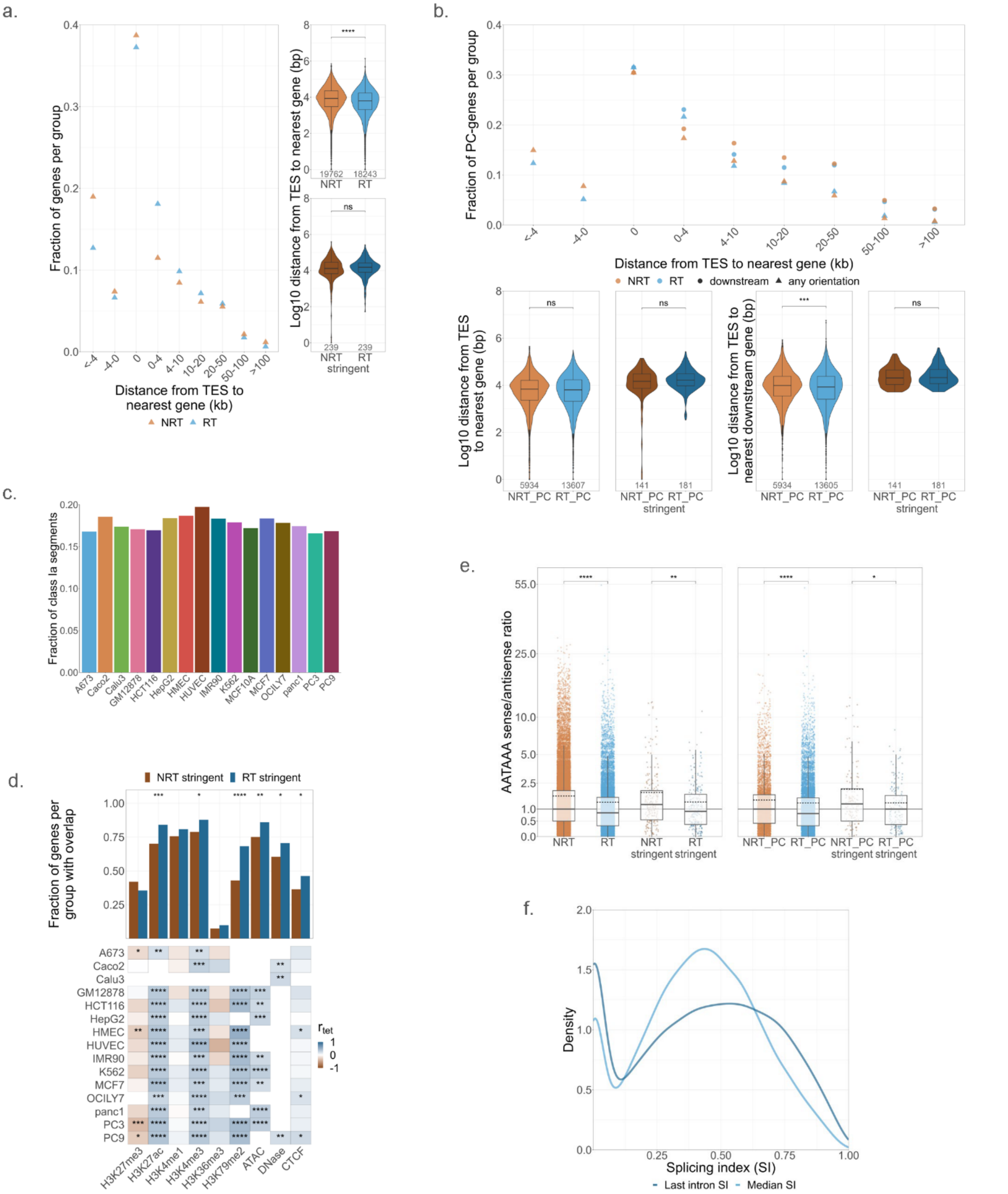

**Supplementary Fig. 14 | RT and NRT gene features. a-b.** Distances to the nearest downstream gene (circle) or the nearest gene in either direction (triangle) for RT or NRT genes. These data are shown for all and stringent RT/NRT genes (**a**) and for protein-coding gene subsets (**b**). The overall log10 distribution of distances between genes (violin) is shown for all RT/NRT genes and stringent RT/NRT genes, with FDR adjusted p-values determined from Wilcoxon tests. The fractions of RT/NRT genes that have a downstream gene within various distance ranges are shown (dot plot). **c.** The fraction of RTsegs that are class Ia and have no other class designations (i.e. overlap no genes on either strand). **d.** Overlap of stringent RT/NRT genes with various chromatin features in the 1 kb downstream of the TSS. The fraction of RT/NRT genes that had each of these features in any cell line is shown (top), along with the tetrachoric correlation

($r_{tet}$) between RT/NRT genes and the presence or absence of each feature per cell line (bottom). Fisher's exact p-values are designated where significant: *$p < 0.05$, **$p < 0.01$, ***$p < 0.001$, ****$p < 0.0001$. **e.** The distributions of PAS sense/antisense ratios for RT/NRT genes. Stringent and protein-coding gene sets are also shown. FDR adjusted p-values were determined from Wilcoxon tests. f. The distribution of gene-based median and last intron SI values for RT genes, showing RT genes can be both poorly- and well-spliced. Adjusted p-values (**a-b,e**) are denoted as: ns = not significant, *$p_{adj} < 0.05$, **$p_{adj} < 0.01$, ***$p_{adj} < 0.001$, ****$p_{adj} < 0.0001$.

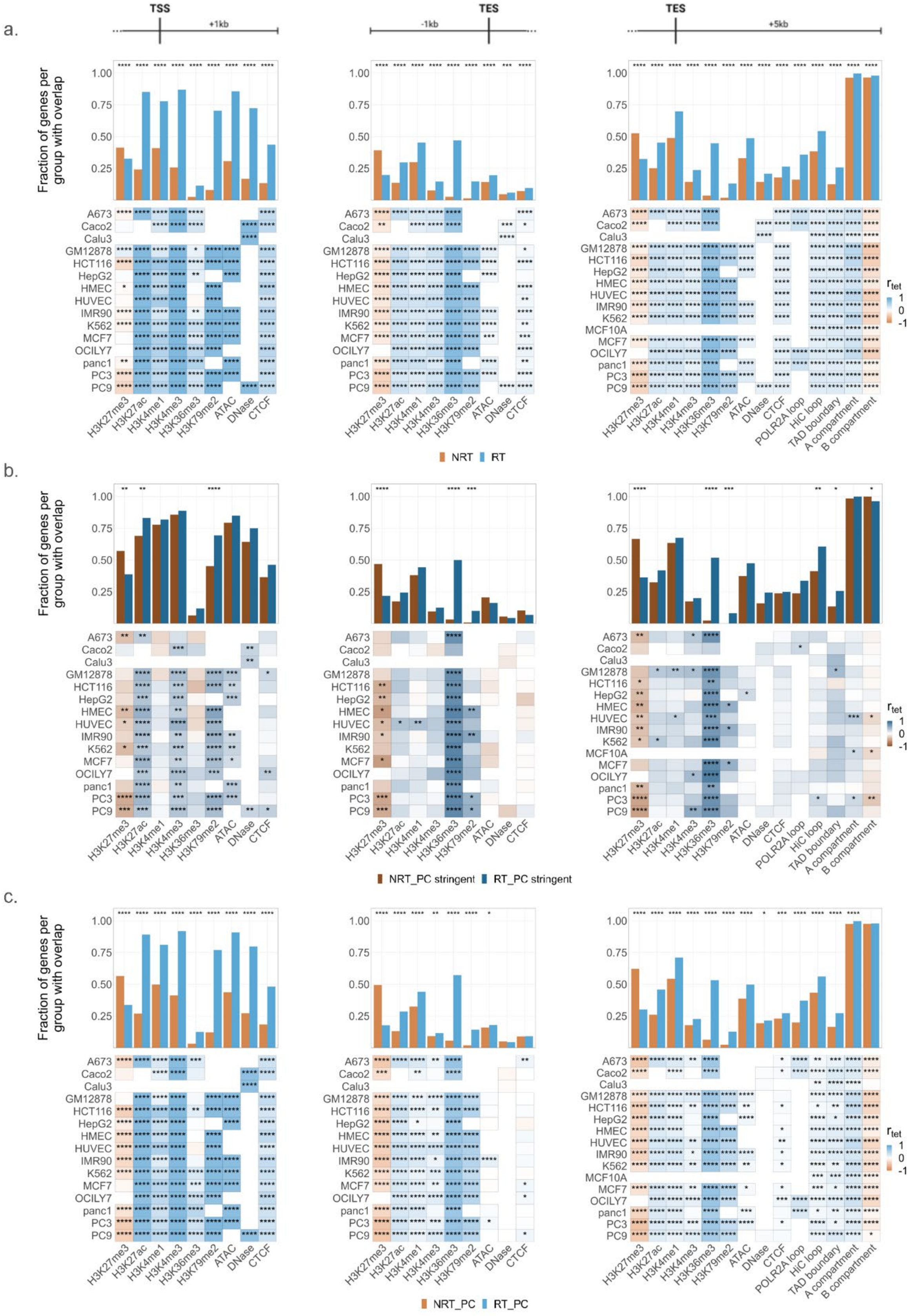

a.
b.
c.
TSS
+1kb
-1kb
TES
+5kb
Fraction of genes per group with overlap
A673
Caco2
Calu3
GM12878
HCT116
HepG2
HMEC
HUVEC
IMR90
K562
MCF10A
MCF7
OCILY7
panc1
PC3
PC9
H3K27me3
H3K27ac
H3K4me1
H3K4me3
H3K36me3
H3K79me2
ATAC
DNase
CTCF
POLR2A loop
HiC loop
TAD boundary
A compartment
B compartment
r tet
NRT
RT
NRT_PC stringent
RT_PC stringent
NRT_PC
RT_PC

**Supplementary Fig. 15 | Chromatin features of RT and NRT genes.** Overlap of stringent RT/NRT genes with various chromatin features in the 1 kb downstream of the TSS, the 1 kb upstream of the TES, and 5 kb downstream of the TES. This is shown for all (**a**), stringent protein-coding (**b**), and all protein-coding (**c**) RT/NRT gene subsets. The fraction of RT/NRT genes that had each of these features in any cell line is shown (top), along with the tetrachoric correlation ($r_{tet}$) between RT/NRT genes and the presence or absence of each feature per cell line (bottom). Fisher's exact p-values are designated where significant: *$p < 0.05$, **$p < 0.01$, ***$p < 0.001$, ****$p < 0.0001$.

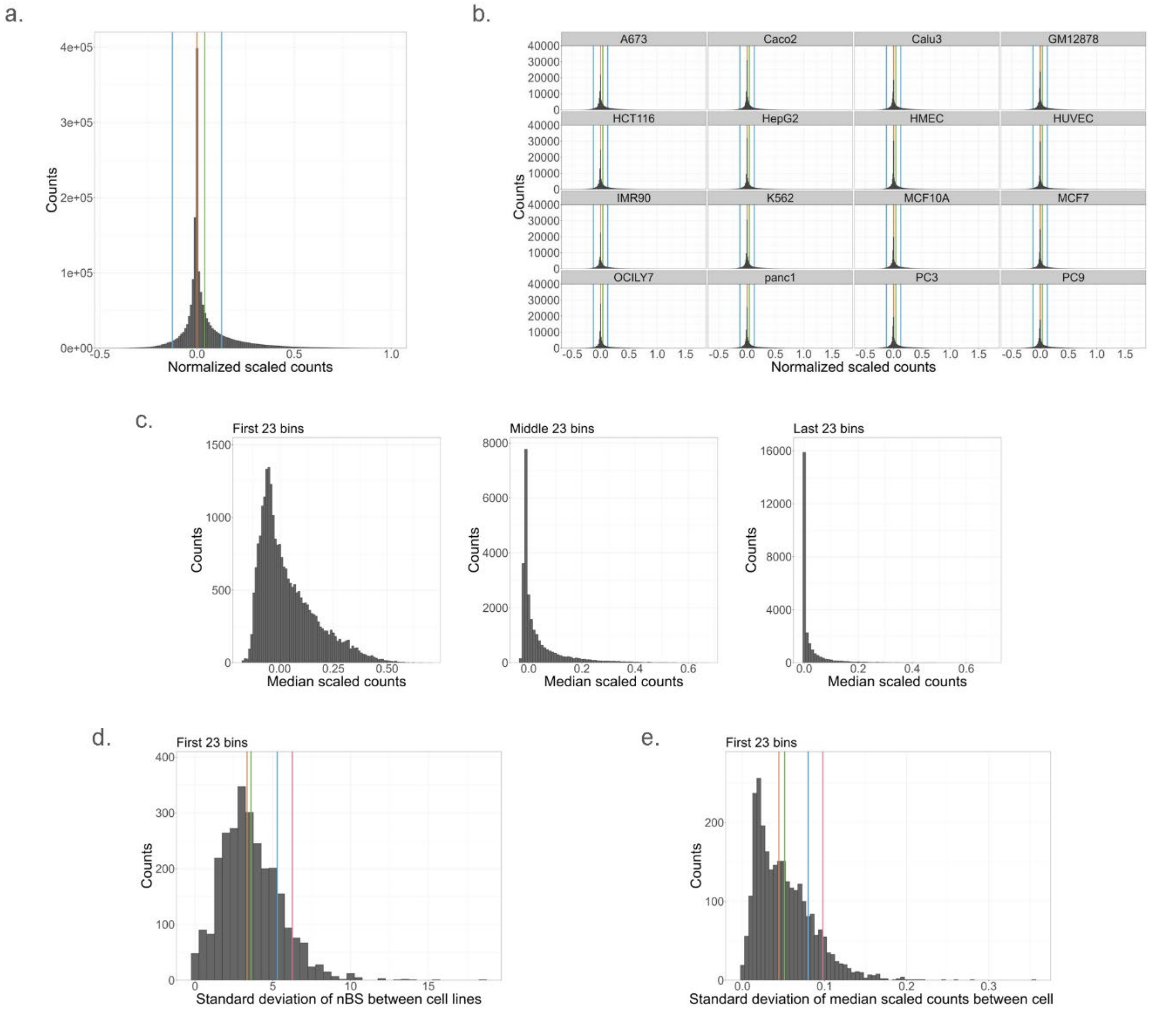

**Supplementary Fig. 16 | RT signal scaled counts and net bin score distributions. a-b.** Distribution of binned normalized scaled counts aggregated for all cell lines (**a**) or each cell line individually (**b**). The mean (green) and median (orange) of the distribution in a are indicated for all. The median is equal to zero due to the normalization of all binned counts to the median scaled signal. One standard deviation from the median (blue) in both directions is indicated, which was the threshold imposed to designate binned signals as significantly different from the median aggregated signal. **c.** Distribution of median scaled counts across cell lines for all 23 bins in each sector of the 17.4 kb RT signal region. **d-e.** Distribution of standard deviations between all cell lines for the net bin score (nBS, **d**) and median scaled counts (**e**). The mean (green), median (orange), and 1 (blue) and 1.5 (pink) standard deviations away from the median, which were used to find cell lines with highly variable nBSs and median scaled counts.

### Supplemental References

### ENCODE portal links

L1. Encode portal - https://www.encodeproject.org

L2. Deeply profiled cell line data matrix – https://www.encodeproject.org/deeply-profiled-uniform-batch-matrix/?type=Experiment&control_type!=*&status=released&replicates.library.biosample.biosample_ontology.term_id=EFO:0002106&replicates.library.biosample.biosample_ontology.term_id=EFO:0001203&replicates.library.biosample.biosample_ontology.term_id=EFO:0006711&replicates.library.biosample.biosample_ontology.term_id=EFO:0002713&replicates.library.biosample.biosample_ontology.term_id=EFO:0002847&replicates.library.biosample.biosample_ontology.term_id=EFO:0002074&replicates.library.biosample.biosample_ontology.term_id=EFO:0001200&replicates.library.biosample.biosample_ontology.term_id=EFO:0009747&replicates.library.biosample.biosample_ontology.term_id=EFO:0002824&replicates.library.biosample.biosample_ontology.term_id=CL:0002327&replicates.library.biosample.biosample_ontology.term_id=CL:0002618&replicates.library.biosample.biosample_ontology.term_id=EFO:0002784&replicates.library.biosample.biosample_ontology.term_id=EFO:0001196&replicates.library.biosample.biosample_ontology.term_id=EFO:0001187&replicates.library.biosample.biosample_ontology.term_id=EFO:0002067&replicates.library.biosample.biosample_ontology.term_id=EFO:0001099&replicates.library.biosample.biosample_ontology.term_id=EFO:0002819&replicates.library.biosample.biosample_ontology.term_id=EFO:0009318&replicates.library.biosample.biosample_ontology.term_id=EFO:0001086&replicates.library.biosample.biosample_ontology.term_id=EFO:0007950&replicates.library.biosample.biosample_ontology.term_id=EFO:0003045&replicates.library.biosample.biosample_ontology.term_id=EFO:0003042&replicates.library.biosample.internal_tags=Deeply%20Profiled

L3. OMNI ATAC-seq protocol – https://www.encodeproject.org/documents/74d62deb-b150-44ff-a53d-0ad141ff621b/@@download/attachment/ENCODE4_ATAC_Omni_CellLines_v1.pdf

L4. snATAC-seq nuclear isolation protocol – https://www.encodeproject.org/documents/25ed0d2e-ba3b-4951-8a20-711f5495c151/@@download/attachment/Snyder_Nuclei_Isolation_tissue_10xscATAC.pdf

L5. Single cell ATAC-seq protocol – https://www.encodeproject.org/documents/e69bfc96-1fc8-4b36-b9a7-9a3e7623c682/@@download/attachment/ENCODE4_scATAC_10x_v1.pdf

L6. Bru-seq protocol - https://www.encodeproject.org/documents/3d850e66-46e9-43ea-bf4c-650e65128123/@@download/attachment/Bru-seq_Experiment_Protocol_v1.0.pdf

L7. Bru-seq with ionizing radiation treatment protocol - https://www.encodeproject.org/documents/35fc1bc7-0655-4318-bb97-3f24304a7d98/@@download/attachment/Bru-seq_IR_Treatment_Experiment_Protocol_v1.0.pdf

L8. 2 hour and 6 hour BruChase-seq protocol - https://www.encodeproject.org/documents/7cb93b63-c366-4e71-9095-00ca7bab3f3f/@@download/attachment/BruChase-seq_Experiment_Protocol_v1.0.pdf

L9. BruUV-seq protocol - https://www.encodeproject.org/documents/38a4efcf-22a2-496b-826b-aa03751b3933/@@download/attachment/BruUV-seq_Experiment_Protocol_v1.0.pdf

L10. ChIA-PET protocol - https://www.encodeproject.org/documents/f7779792-9403-4396-9ff6-8e859edd608c/@@download/attachment/In-situ%20chiapet_Ruan%20Lab_final.pdf

L11. DNase-seq protocol - https://www.encodeproject.org/documents/980de32c-f7ec-4cd7-b736-61faf723c186/@@download/attachment/newnuclei_isolation_from_human_tissue-combined.pdf

L12. Intact Hi-C protocol - https://www.encodeproject.org/documents/768fa33e-3c32-4ce2-8f78-b9fafda06cbc/@@download/attachment/intacthi-c_modularprotocol-release.pdf

L13. PRO-cap protocol - https://www.encodeproject.org/documents/8a82200a-023f-439d-ae91-a1a1bf80eaa0/@@download/attachment/20220128_PROcap_Experimental_Protocol.pdf

L14. Long-read RNA-seq protocol - https://www.encodeproject.org/documents/3baa46d2-cb88-4608-8877-70596d200489/@@download/attachment/ENCODE_longread_wetlab_protocolv3.pdf

L15. miRNA-seq protocol - https://www.encodeproject.org/documents/49f43842-5ab4-4aa1-a6f4-2b1234955d93/@@download/attachment/Multiplexed%20miRNA%20Sequencing%20Library%20Generation%20Protocol-3.4.pdf

L16. Total RNA-seq protocol - https://www.encodeproject.org/documents/01a7807d-0d18-4a4a-ac90-f4056a25a681/@@download/attachment/manualE7760_Directional_RNA_library_kit_October30_2018.pdf

L17. ChIA-PET pipeline readme - https://www.encodeproject.org/documents/47cf3c63-12c4-4b32-afb5-0e8141918ffa/@@download/attachment/ENCODE_ChIA_PET_pipeline_readme_12172020.pdf

L18. Bru-seq data processing pipeline overview - https://www.encodeproject.org/documents/2f3ba718-5daf-4604-9d47-b886fe257143/@@download/attachment/Bru-seq%20Pipeline%20Overview%20Document%20v1.0.pdf

### References

82. Nurk, S. *et al.* The complete sequence of a human genome. *Science* **376**, 44-53 (2022).

83. Magnuson, B. *et al.* Identifying transcription start sites and active enhancer elements using BruUV-seq. *Sci Rep* **5**, 17978 (2015).

84. Tornaletti, S. & Hanawalt, P.C. Effect of DNA lesions on transcription elongation. *Biochimie* **81**, 139-46 (1999).
85. Chatterjee, N. & Walker, G.C. Mechanisms of DNA damage, repair, and mutagenesis. *Environ Mol Mutagen* **58**, 235-263 (2017).
86. Shen, J.C., Fox, E.J., Ahn, E.H. & Loeb, L.A. A rapid assay for measuring nucleotide excision repair by oligonucleotide retrieval. *Sci Rep* **4**, 4894 (2014).
87. Andrade-Lima, L.C., Veloso, A., Paulsen, M.T., Menck, C.F. & Ljungman, M. DNA repair and recovery of RNA synthesis following exposure to ultraviolet light are delayed in long genes. *Nucleic Acids Res* **43**, 2744-56 (2015).
88. Corces, M.R. *et al.* An improved ATAC-seq protocol reduces background and enables interrogation of frozen tissues. *Nat Methods* **14**, 959-962 (2017).
89. Li, X. *et al.* Long-read ChIA-PET for base-pair-resolution mapping of haplotype-specific chromatin interactions. *Nat Protoc* **12**, 899-915 (2017).
90. Mahat, D.B. *et al.* Base-pair-resolution genome-wide mapping of active RNA polymerases using precision nuclear run-on (PRO-seq). *Nat Protoc* **11**, 1455-76 (2016).
91. Mortazavi, A., Williams, B.A., McCue, K., Schaeffer, L. & Wold, B. Mapping and quantifying mammalian transcriptomes by RNA-Seq. *Nat Methods* **5**, 621-8 (2008).
92. Buenrostro, J.D., Wu, B., Chang, H.Y. & Greenleaf, W.J. ATAC-seq: A Method for Assaying Chromatin Accessibility Genome-Wide. *Curr Protoc Mol Biol* **109**, 21 29 1-21 29 9 (2015).
93. Amemiya, H.M., Kundaje, A. & Boyle, A.P. The ENCODE Blacklist: Identification of Problematic Regions of the Genome. *Sci Rep* **9**, 9354 (2019).
94. Lee, B. *et al.* ChIA-PIPE: A fully automated pipeline for comprehensive ChIA-PET data analysis and visualization. *Sci Adv* **6**, eaay2078 (2020).
95. Yao, L. *et al.* A comparison of experimental assays and analytical methods for genome-wide identification of active enhancers. *Nat Biotechnol* **40**, 1056-1065 (2022).
96. Wyman, D. *et al.* A technology-agnostic long-read analysis pipeline for transcriptome discovery and quantification. *bioRxiv*, 672931 (2020).
97. Preker, P. *et al.* PROMoter uPstream Transcripts share characteristics with mRNAs and are produced upstream of all three major types of mammalian promoters. *Nucleic Acids Res* **39**, 7179-93 (2011).
98. Gallego Romero, I. & Lea, A.J. Leveraging massively parallel reporter assays for evolutionary questions. *Genome Biol* **24**, 26 (2023).
99. Gschwind, A.R. *et al.* An encyclopedia of enhancer-gene regulatory interactions in the human genome. *bioRxiv* (2023).
100. Vilborg, A., Passarelli, M.C., Yario, T.A., Tycowski, K.T. & Steitz, J.A. Widespread Inducible Transcription Downstream of Human Genes. *Mol Cell* **59**, 449-61 (2015).
101. Drasgow, F. Polychoric and polyserial correlations. *The Encyclopedia of Statistics* **7**, 68-74 (1986).
102. Langmead, B. & Salzberg, S.L. Fast gapped-read alignment with Bowtie 2. *Nat Methods* **9**, 357-9 (2012).
103. Langmead, B., Trapnell, C., Pop, M. & Salzberg, S.L. Ultrafast and memory-efficient alignment of short DNA sequences to the human genome. *Genome Biol* **10**, R25 (2009).
104. Langmead, B., Wilks, C., Antonescu, V. & Charles, R. Scaling read aligners to hundreds of threads on general-purpose processors. *Bioinformatics* **35**, 421-432 (2019).
105. Danecek, P. *et al.* Twelve years of SAMtools and BCFtools. *Gigascience* **10**(2021).
106. Dobin, A. *et al.* STAR: ultrafast universal RNA-seq aligner. *Bioinformatics* **29**, 15-21 (2013).
107. Love, M.I., Huber, W. & Anders, S. Moderated estimation of fold change and dispersion for RNA-seq data with DESeq2. *Genome Biol* **15**, 550 (2014).
